# Chemical Tools for Profiling the Intracellular ADP-ribosylated Proteome

**DOI:** 10.1101/2023.08.30.555609

**Authors:** Simeon D. Draganov, Michael J. Gruet, Daniel Conole, Cristina Balcells, Alexandros P. Siskos, Hector C. Keun, Dorian O. Haskard, Edward W. Tate

## Abstract

The post-translational modification (PTM) ADP-ribosylation plays an important role in cell signaling and regulating protein function and has been implicated in the development of multiple diseases, including breast and ovarian cancers. Studying the underlying mechanisms through which this PTM contributes towards disease development, however, has been hampered by the lack of appropriate tools for reliable identification of physiologically relevant ADP-ribosylated proteins in a live-cell environment. Herein, we explore the applications of an alkyne-tagged proprobe, 6Yn-ProTide-Ad (6Yn-Pro) as a chemical tool for the identification of intracellular ADP-ribosylated proteins through metabolic labelling. We applied targeted metabolomics and chemical proteomics in HEK293T cells treated with 6Yn-Pro to demonstrate intracellular metabolic conversion of the probe into ADP-ribosylation cofactor 6Yn-NAD^+^, and subsequent labelling and enrichment of PARP1 and multiple known ADP-ribosylated proteins in cells under hydrogen peroxide-induced stress. We anticipate that the approach and methodology described here will be useful for future identification of novel intracellular ADP-ribosylated proteins.

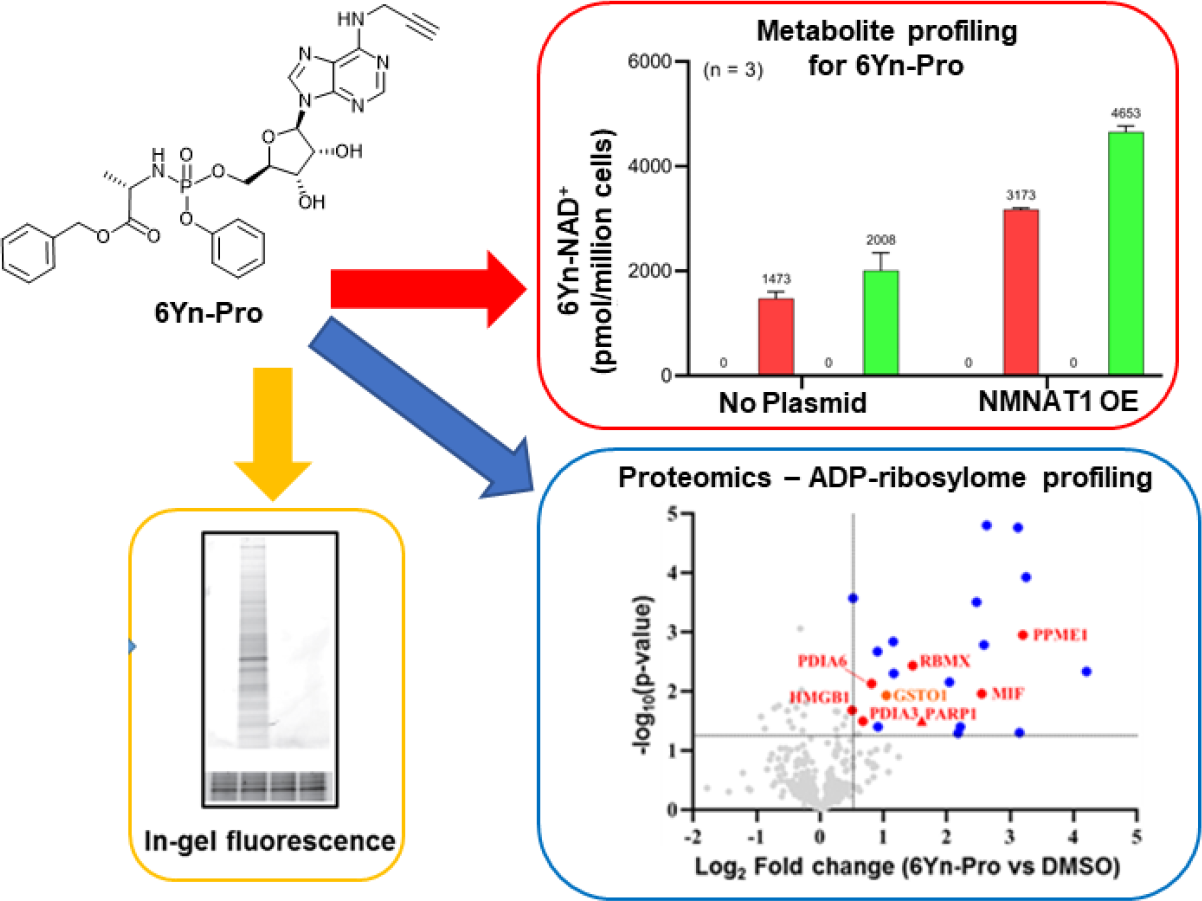

## INTRODUCTION

ADP-ribosylation is a type of post-translational modification most commonly found on proteins.^1,2^ It is catalyzed by poly(ADP-ribose) polymerases (PARPs), a family of 17 enzymes, which catalyze covalent coupling of ADP-ribose units, derived from the cellular cofactor nicotinamide adenine dinucleotide (NAD^+^), to protein substrates.^3,4^ PARPs are broadly divided into two classes: mono-ADP-ribosylating (MARylating) PARPs, which catalyze the addition of a single ADP-ribose unit to a protein target, a process known as mono-ADP-ribosylation (MARylation), and polyADP-ribosylating (PARylating) PARPs, which catalyze the addition of multiple (up to 200) ADP-ribose units to a protein target, a process known as poly-ADP-ribosylation (PARylation).^4,5^ PARPs have been implicated in a very wide range of biological processes including repair of single- and double-strand DNA breaks as part of the DNA damage response (DDR),^6-8^ RNA metabolism by altering localization and activity of RNA-binding proteins,^9^ antiviral response by repressing viral replication through degradation of viral RNA/proteins and amplification of the interferon response^10-14^, and neuronal development by regulating dendrite morphogenesis^15^. In addition to their roles in regulating essential biological processes PARPs have been implicated in development of disease states such as breast and ovarian cancer, through upregulation of ADP-ribosylation activity.^16-20^ Elucidating the underlying mechanisms linking PARP catalytic activity and disease development is of vital importance for development of targeted treatments, for which identification and study of the protein substrates of ADP-ribosylation in relevant disease models represents an important step.

In recent years, several approaches have been developed to study the biological consequences of ADP-ribosylation through identification of the ADP-ribosylated proteome, and for dissecting the biological roles of individual PARPs. NAD^+^ analogues incorporating a click chemistry tag allow for functionalization of labelled proteins *via* click chemistry conjugation to a fluorophore and/or biotin.^21-35^ This breakthrough enables the use of more complex analytical techniques to study ADP-ribosylation, including visualization of the distribution of ADP-ribosylation events in fixed cells *via* fluorescence microscopy.^24-27^ Furthermore, biotinylation of ADP-ribosylated proteins allows for their selective enrichment from the cellular proteome, and coupled with liquid chromatography-tandem mass spectrometry (LC-MS/MS), has led to the identification of hundreds of putative ADP-ribosylation targets.^23,30-35^

A major limitation on the use of NAD^+^ analogues in the study of ADP-ribosylation dynamics and identification of relevant protein substrates is their inability to passively cross cell membranes. Studies employing NAD^+^ analogues have therefore been largely confined to experiments involving cell lysates or involve the use of carrier peptides^24^, transfection reagents^26^ or transiently permeabilized cells^27^ to deliver NAD^+^ analogues intracellularly, each of which is limited by disruption of the cell membrane. Two previous studies employing clickable adenosine precursors for the identification of intracellular ADP-ribosylation targets via their metabolic conversion into the corresponding NAD^+^ analogues have been performed.^36,37^ However, evidence is currently lacking for intracellular conversion of precursor probes into the corresponding relevant metabolites: the intermediate adenosine monophosphate (AMP) and adenosine triphosphate (ATP) analogues, and essential NAD^+^ analogue cofactor.

Herein, we explore the suitability of a clickable adenosine-based ProTide proprobe, 6Yn-ProTide-Ad (6Yn-Pro), previously applied to study intracellular AMPylation^38^ as a potential precursor probe for intracellular labelling of ADP-ribosylated proteins. We demonstrate successful intracellular metabolic conversion of the precursor probe 6Yn-Pro into the ADP-ribosylation cofactor 6Yn-NAD^+^ by targeted metabolomics, and subsequent labelling and enrichment of auto-PARylated PARP1 and other known ADP-ribosylated proteins by proteomics. We provide the first evidence for the application of 6Yn-Pro for detection of intracellular protein ADP-ribosylation and anticipate that the probe and described methodology will serve as a basis for further optimization of the approach towards identification of novel intracellular ADP-ribosylated proteins, including non-canonical ADP-ribosylation events.^39^

## RESULTS AND DISCUSSION

6Yn-Pro (**Figure 1A**) has been previously shown to undergo metabolic conversion into 6Yn-ATP in HeLa cells.^38^ In the final step of the NAD^+^ biosynthesis pathway (**Figure 1B**), ATP and beta-nicotinamide mononucleotide (β-NMN) are coupled by nicotinamide mononucleotide adenylyltransferases (NMNAT1, 2, 3) to produce NAD^+^,^40^ and we hypothesized that 6Yn-ATP may also undergo intracellular metabolic conversion into the NAD^+^ analogue 6Yn-NAD^+^. Subsequently, PARPs might use metabolically generated 6Yn-NAD^+^ for intracellular protein ADP-ribosylation, in line with previous studies in cell lysates or with recombinant PARP1.^22-24^ We therefore sought to explore whether 6Yn-Pro could be used for successful intracellular labelling of ADP-ribosylated proteins. In addition, we sought to establish experimental conditions that promote metabolic conversion of 6Yn-ATP into 6Yn-NAD^+^ and thus favor protein ADP-ribosylation over competing AMPylation modification (**Figure S1A**) following probe treatment.

**Figure 1.**
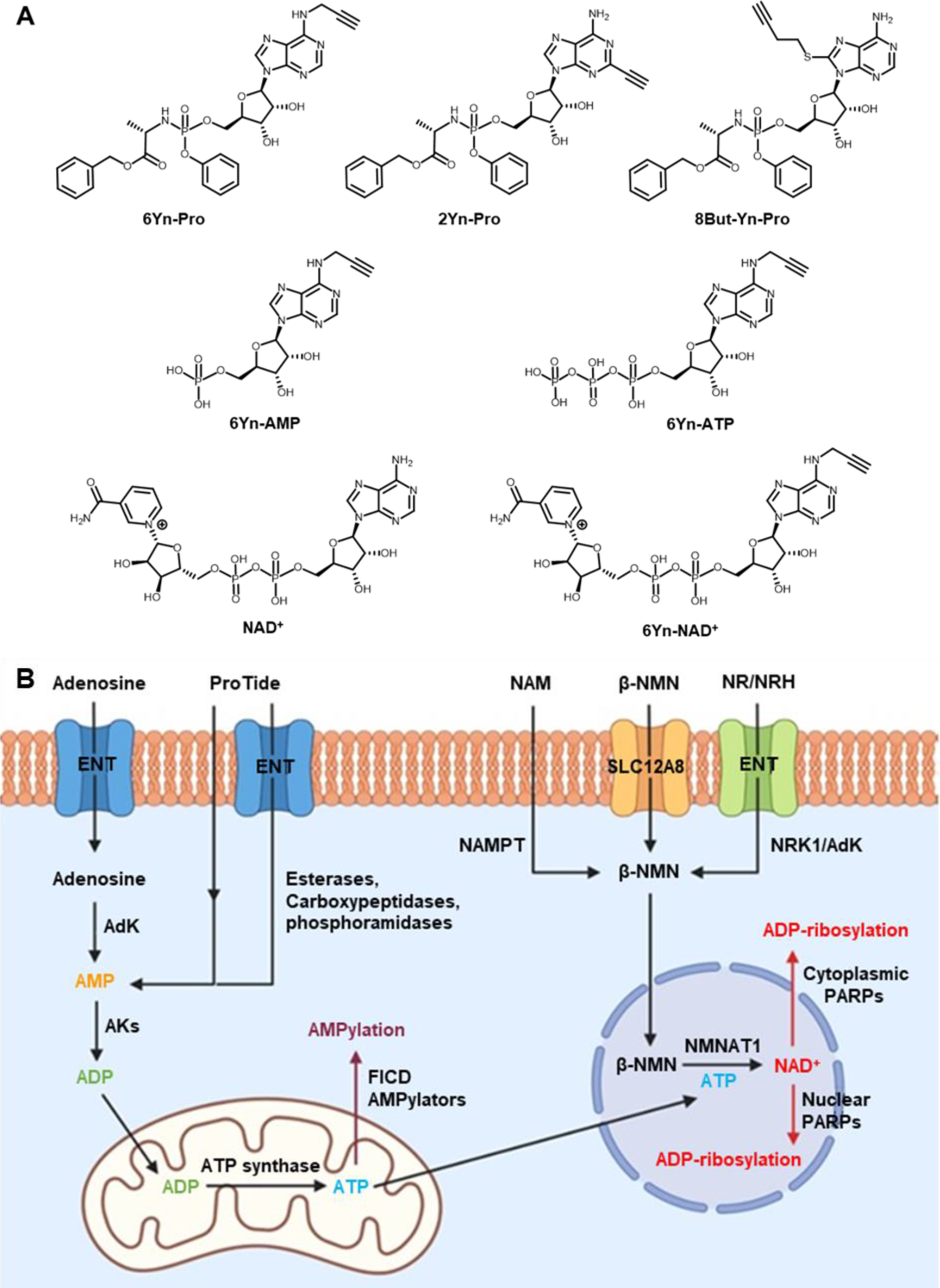
Chemical structures of probes and metabolites, and illustration of compartmentalization of intracellular ATP, β-NMN and nuclear NAD^+^ biosynthesis. (**A**) Chemical structures of the three ProTide probes, 6Yn-Pro, 2Yn-Pro and 8But-Yn-Pro, and metabolites 6Yn-AMP, 6Yn-ATP, 6Yn-NAD^+^ and NAD^+^. (**B**). Extracellular adenosine (or ProTide-Ad proprobe) is transported into cells through equilibrative nucleoside transporters (ENTs) and converted into AMP by adenosine kinase (AdK), and then ADP by adenylate kinases (AKs), in the cytoplasm. ADP is subsequently transported into the mitochondria and converted into ATP by ATP synthase. ATP is then used in protein AMPylation by AMPylators, such as FICD, or transported into the nucleus for nuclear NAD^+^ biosynthesis *via* coupling to β-NMN by NMNAT1, and subsequent ADP-ribosylation by PARPs. β-NMN is biosynthesized from nicotinamide (NAM) by nicotinamide phosphoribosyltransferase (NAMPT), or through metabolic conversion of nicotinamide riboside (NR) or reduced nicotinamide riboside (NRH) by nicotinamide riboside kinase 1 (NRK1) or adenosine kinase (AdK), respectively. Extracellular β-NMN can also be transported intracellularly by cells expressing the β-NMN transporter channel SLC12A8.

In order to explore the structure-activity relationship among related adenosine probe analogues, 6Yn-Pro, along with two novel ProTide probes, 2Yn-ProTide-Ad (2Yn-Pro) and 8But-Yn-ProTide-Ad (8But-Yn-Pro), and the two adenosine analogues, 6Yn-Ad and 2Yn-Ad,^37^ were synthesized (**Figure 1A**; see **Supplementary Methods**). Intracellular protein labelling was first investigated by in-gel fluorescence (IGF), by following the workflow outlined in Figure 2. Following treatment of HEK293T and T47D cells with each probe or DMSO (negative control), the cells were lysed, proteins were precipitated using methanol:chloroform, and probe-labelled proteins were conjugated to azide-5-carboxytetramethylrhodamine (TAMRA)-biotin (AzTB; **Figure S1B**) by copper-catalyzed azide-alkyne cycloaddition (CuAAC).

**Figure 2.**
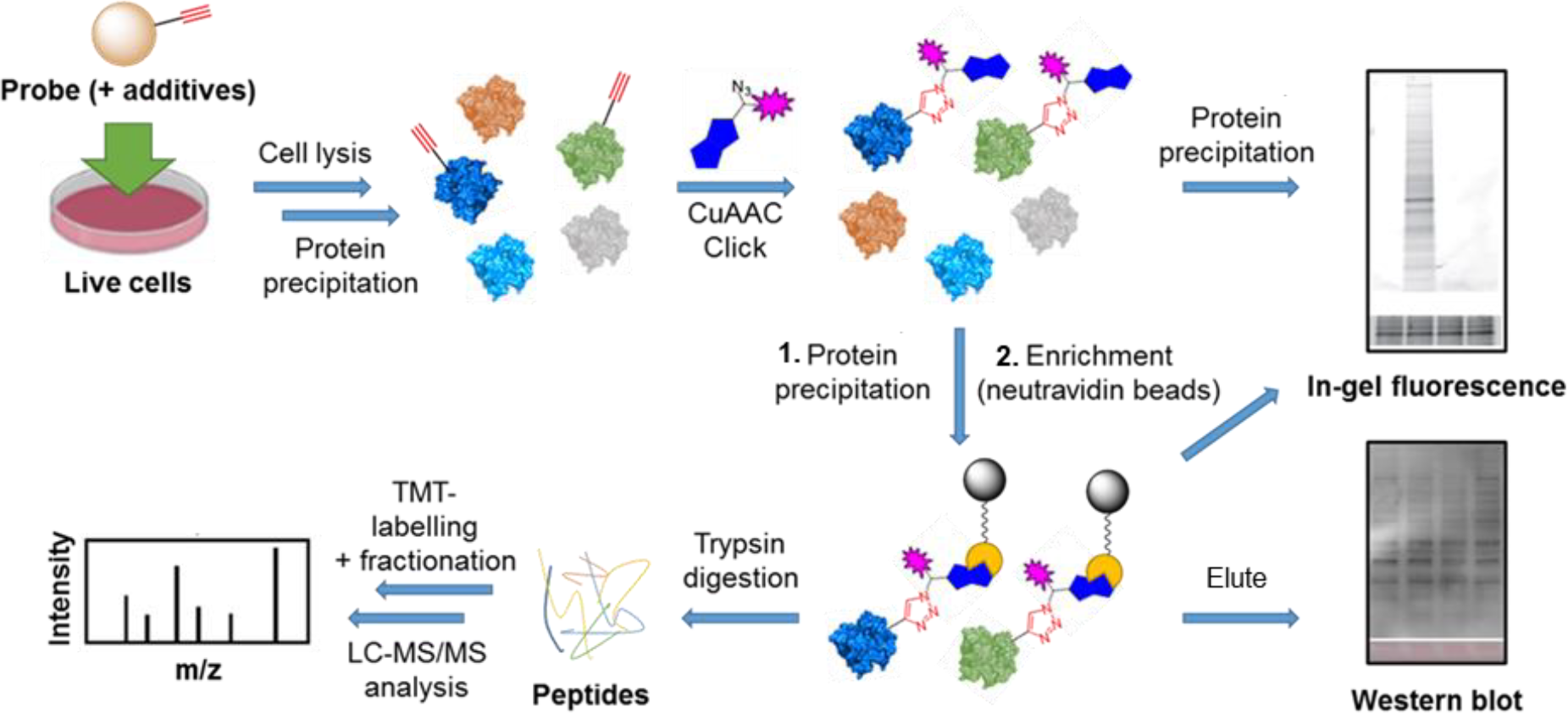
Overview of the workflow followed for sample preparation for analysis by in-gel fluorescence, western blot, and proteomics. Cell treatment with the relevant probe(s) and additive(s) is performed, typically for 24 h, followed by cell lysis and a protein precipitation step. Probe-labelled proteins are then captured by CuAAC click chemistry conjugation to a trifunctional azide-TAMRA-biotin capture reagent, allowing for visualization of protein labelling *via* its fluorophore by IGF, or enrichment (pull-down) of labelled proteins *via* its biotin moiety by streptavidin affinity chromatography. For analysis by IGF and western blot, enriched proteins are eluted from the streptavidin beads by boiling; for proteomics analysis, on-bead digestion of the enrichment proteins is performed, and the corresponding peptides analyzed by LC-MS/MS for protein identification.

### Clickable adenosine-based probes non-specifically label proteins in lysates

Upon performing IGF experiments we discovered that the adenosine-based analogue probes 6Yn-Ad and 2Yn-Ad, including 6Yn-Pro, non-specifically interacted with proteins in denatured, enzymatically inactive lysates (of MDA-MB-231 cells), resulting in protein labelling being observed in the absence of metabolic and enzymatic activity (**Figure S2A**). The observed protein labelling pattern was identical in probe-treated live MDA-MB-231 cells (**Figure S2B**), providing evidence that non-specific protein labelling also occurred in live-cell samples. These findings have important implications for downstream sample processing, including proteomics experiments where non-specifically labelled proteins may appear significantly enriched relative to untreated control samples, leading to false identification of protein targets. Indeed, the IGF labelling pattern observed in adenosine analogue-treated samples in the two previous proteomics studies^36,37^ was very similar to the pattern we observed in denatured lysates, suggesting that non-specifically labelled proteins were also analyzed in these previously reported proteomics experiments.

### Protein precipitation prior to click chemistry conjugation removes non-specific protein labelling

We discovered that adding a protein precipitation step in methanol:chloroform after cell lysis and prior to click chemistry conjugation resulted in removal of this non-specific protein labelling (**Figure S2B and S2C**). Protein precipitation in methanol:chloroform is a highly efficient process with minimal loss of protein, as evidenced by Coomassie stained loading controls (**Figure S2B and S2C**), confirming that loss of protein labelling was not due to overall loss of protein loading following protein precipitation. This led us to hypothesize that on cell treatment adenosine and ProTide-Ad probes enter cells and are released on cell lysis, leading to non-enzyme mediated chemical conjugation, which in turn results in the observed non-specific protein labelling. Protein precipitation removes residual probe released from cells on lysis and eliminates non-specific protein labelling, preventing subsequent capture by CuAAC.

By following this approach, subsequent comparison of probe-treated and DMSO-treated cells after protein enrichment revealed extensive protein labelling in probe-treated samples (**Figure S3A and S3B**), confirming that the probes were metabolically converted into biologically active cofactors and subsequently attached to proteins, possibly as part of a PTM. Performing the precipitation step also demonstrated that 6Yn-Pro was superior in downstream incorporation into PTMs, and hence protein labelling, compared to adenosine analogues 6Yn-Ad and 2Yn-Ad. This was evidenced by the presence of protein bands following removal of the non-specific labelling in 6Yn-Pro-treated cells (**Figure S3A and S3B**) by precipitation, in contrast to 6Yn-Ad and 2Yn-Ad-treated cells where protein labelling entirely disappeared following removal of non-specific labelling by precipitation (**Figure S2C**), except for a single band at ~75 kDa in 2Yn-Ad treated cells.

### Non-specific probe-protein interactions are facilitated by ribose ring in adenosine

To uncover the origin of the non-specific probe-protein interactions, 6Yn-adenine was synthesized, and lysates and live cells were treated with this probe. 6Yn-adenine did not result in non-specific protein labelling in either case (**Figure S2B**), demonstrating that this phenomenon is facilitated by the ribose ring in adenosine. These findings potentially imply that nucleoside-based clickable probes other than adenosine might also suffer from this same drawback, which should be considered when using such probes in protein labelling experiments and warrants further investigation to confirm whether this phenomenon occurs with other clickable nucleoside probes. Although not further investigated in the present study, reversible cross-linking of the 2’ and 3’ hydroxyl groups of the ribose ring of the probes to carbonyl moieties in proteins could be a plausible mechanism for the reversible, non-enzymatic non-specific labelling of probe-treated samples.

### 6Yn-Pro is metabolically converted into ADP-ribosylation cofactor 6Yn-NAD^+^

To determine the extent of metabolic incorporation of 6Yn-Pro in NAD^+^ biosynthesis in cells, we employed targeted liquid chromatography-mass spectrometry (LC-MS) metabolomics^41^ (**Figure S4**) to measure the abundance of the key metabolites expected to arise following 24-hour treatment with 6Yn-Pro: 6Yn-AMP, 6Yn-ATP, and 6Yn-NAD^+^. We also explored the effect of supplementation with reduced nicotinamide riboside (NRH), a β-NMN/NMNH and NAD^+^ precursor, on 6Yn-NAD^+^ levels, which has previously been shown to increase intracellular NAD^+^ levels 3-to 10-fold in mammalian cell lines through metabolic conversion into β-NMNH^42-44^. Finally, we investigated whether NMNAT1 overexpression (OE), responsible for conversion of ATP into NAD^+^ in the nucleus^40^, resulted in increased 6Yn-NAD^+^ production. Analysis of extracted metabolites in cells treated only with 6Yn-Pro revealed the presence of all three metabolite analogues (**Figure 3A-C**), demonstrating that the probe is cell-permeable, undergoes metabolic conversion into 6Yn-AMP (**Figure 3A**), and is used to produce 6Yn-ATP (**Figure 3B**) and 6Yn-NAD^+^ (**Figure 3C**). Following 6Yn-Pro probe co-treatment with NRH, β-NMN levels increased ~4-fold compared to 6Yn-Pro treatment alone (**Figure S5E**), whereas 6Yn-ATP levels decreased by 28%, (**Figure 3B**) and 6Yn-NAD^+^ levels increased by 36%, (**Figure 3C**). Collectively, the data indicate increased production of 6Yn-NAD^+^ possibly due to increased coupling of 6Yn-ATP and β-NMN/NMNH. We also measured the endogenous levels of ATP and NAD^+^ (**Figure S5A and S5B**) and calculated % incorporation of 6Yn-ATP and 6Yn-NAD^+^ in the endogenous ATP and NAD^+^ pools, which corresponded to 9.0% and 0.26% in the probe-only treated samples, respectively (**Figure S5C and S5D**). The observed disparity between % incorporation of 6Yn-ATP and 6Yn-NAD^+^ in their respective pools suggests that the final conversion step by NMNAT1 may be rate-limiting.

**Figure 3.**
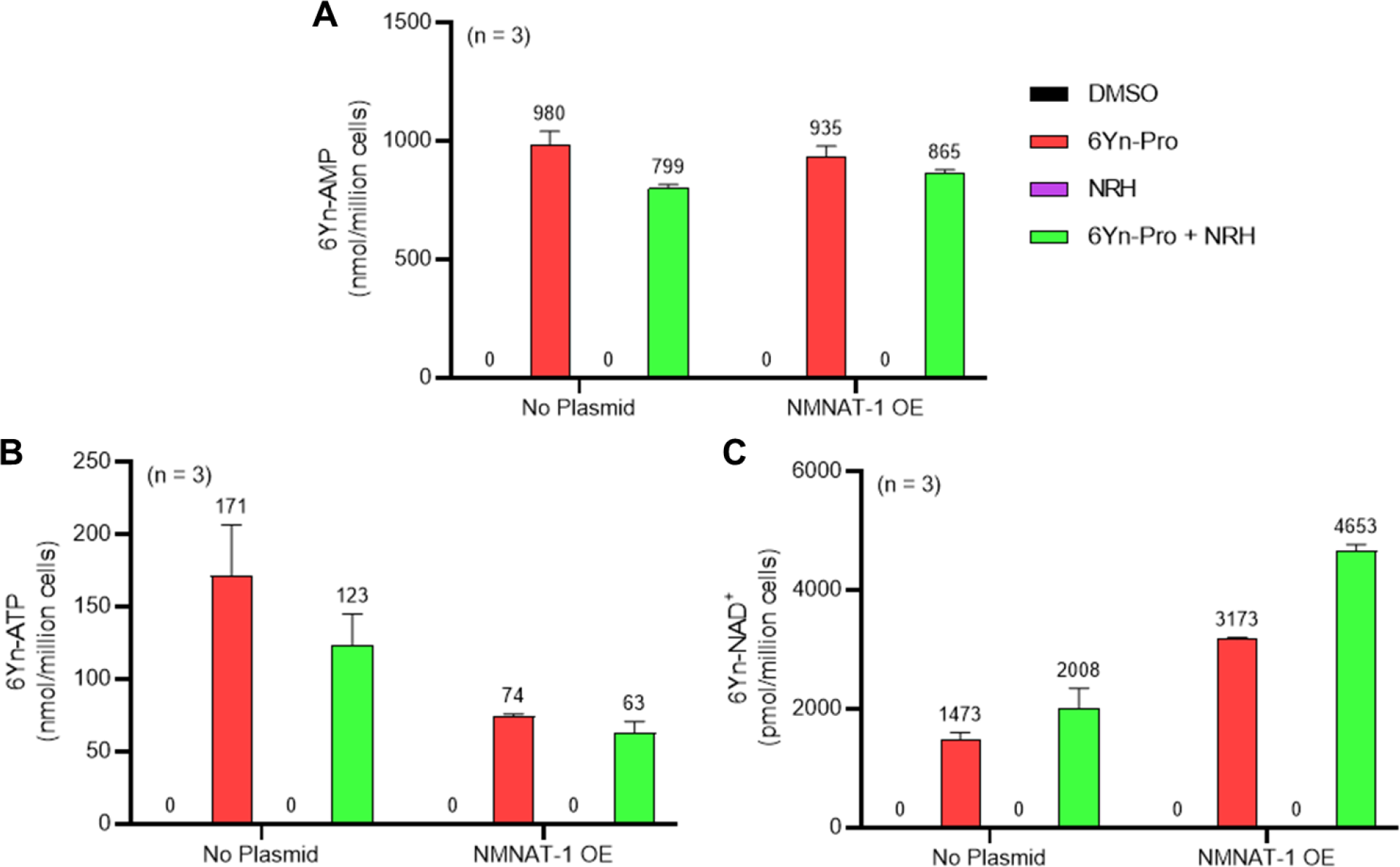
Metabolite profiling in HEK293T cells reveals intracellular conversion of 6Yn-Pro into 6Yn-ATP and 6Yn-NAD^+^. Metabolite profiling was performed following treatment with probe 6Yn-Pro (100 µM, 24 h), with/without NRH co-treatment (500 µM, 24 h), with/without NMNAT1 overexpression (24 h), or DMSO (negative control). Metabolite levels for (**A**) 6Yn-AMP, (**B**) 6Yn-ATP and (**C**) 6Yn-NAD^+^ were measured and quantified.

### NMNAT1 overexpression boosts intracellular production of 6Yn-NAD^+^

To explore whether higher NMNAT1 levels resulted in increased production of 6Yn-NAD^+^, NMNAT1 was overexpressed in HEK293T cells for 24 h prior to probe treatment (**Figure S6**). Although 6Yn-AMP levels were largely unaffected when NMNAT1 was overexpressed (**Figure 3A**), 6Yn-ATP levels decreased by 57% and 49%, in NRH-untreated and NRH-co-treated cells, respectively, when compared to no NMNAT1 overexpression (**Figure 3B**). Furthermore, 6Yn-NAD^+^ levels increased by 115% and 132%, in NRH-untreated and NRH-co-treated cells, respectively, when compared to cells without NMNAT1 overexpression (**Figure 3C**). The drop in 6Yn-ATP levels and the increase in 6Yn-NAD^+^ levels following NMNAT1 overexpression imply increased production of 6Yn-NAD^+^ through increased coupling of the precursor 6Yn-ATP to β-NMN/NMNH, with the greatest effect observed when cells were also supplemented with NRH. Collectively, the data provide the first evidence that the 6Yn-ATP to 6Yn-NAD^+^ conversion step can be successfully targeted to increase metabolic production of 6Yn-NAD^+^ and raise the overall intracellular 6Yn-NAD^+^ levels.

### Metabolically produced 6Yn-NAD^+^ labels PARP1 and ADP-ribosylated proteins intracellularly

Following metabolomics data confirming that 6Yn-Pro undergoes intracellular metabolic conversion into 6Yn-NAD^+^, and IGF experiments confirming intracellular protein labelling, we next sought to identify labelled protein targets in 6Yn-Pro-treated HEK293T cells by proteomics. By following the workflow outlined in **Figure 2**, comparison of protein enrichment was performed for probe-treated vs DMSO-treated samples. We performed a preliminary experiment where HEK293T cells were treated with probe only, co-treated with probe and NRH, treated with probe and hydrogen peroxide (H_2_O_2_) just prior to cell lysis, or co-treated with probe, NRH and H_2_O_2_ (prior to cell lysis), and compared the protein labelling profiles to the corresponding DMSO controls (**Figure 4A-D**). H_2_O_2_-treatment was included since ADP-ribosylation is known to be upregulated under oxidative stress/damage by PARP1/2-hyperactivation^2^. We observed labelling and relative enrichment of PARP1 in all four conditions (**Figure 4A-D**), demonstrating that metabolically generated 6Yn-NAD^+^, derived from 6Yn-Pro, is used by PARP1 for auto-PARylation. The extent of PARP1 labelling and enrichment was lowest in 6Yn-Pro only treated cells (absolute Fold change = 1.8), and highest in cells co-treated with 6Yn-Pro, NRH and H_2_O_2_ (absolute Fold change = 5.5) (**Table S1**). Similarly, the highest proportion of enriched proteins known to be ADP-ribosylated from previous studies was found in cells co-treated with 6Yn-Pro, NRH cells and H_2_O_2_ (51%), and lowest in 6Yn-Pro only treated cells (39%) (**Table S2**). Mirroring the higher 6Yn-NAD^+^ levels observed in cells co-treated with 6Yn-Pro and NRH (**Figure 3C and S5D**), collectively, the data suggest that boosting 6Yn-NAD^+^ levels through supplementation with NRH promoted protein ADP-ribosylation with the analogue. In addition, these data support application of H_2_O_2_ prior to cell lysis to stimulate 6Yn-NAD^+^ incorporation by PARPs.

**Figure 4.**
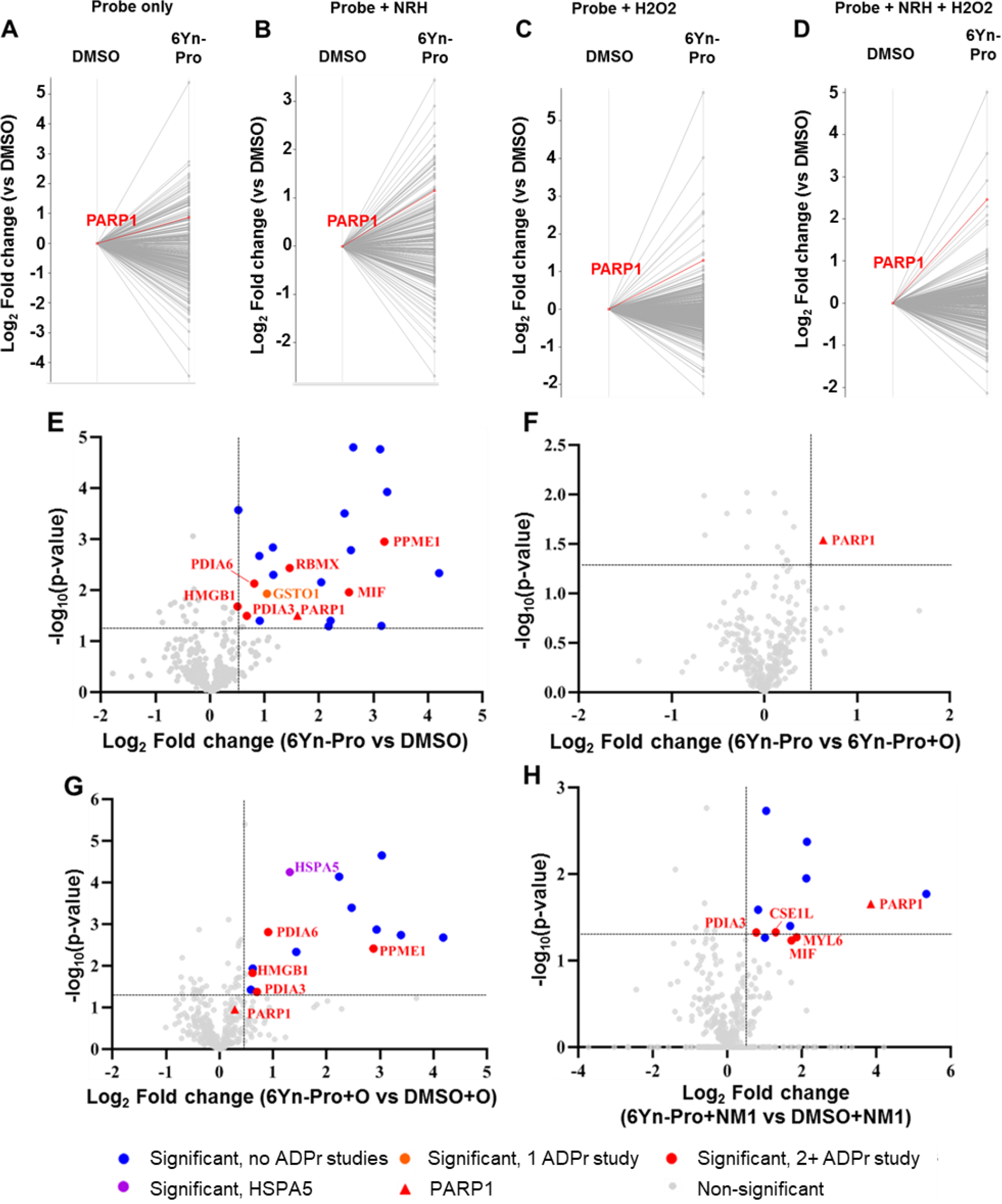
Proteomics analysis reveals protein labelling and enrichment of known ADPr proteins in 6Yn-Pro-treated HEK293T cells. Log_2_ Fold change represents the extent of protein enrichment/labelling in one treatment condition (typically probe treatment) relative to another. -Log_10_(p-value) represents the statistical significance between replicates. (**A-D**) Profile plots showing protein labelling and enrichment, relative to the corresponding DMSO controls, following treatment of HEK293T cells with (**A**) 6Yn-Pro only (100 µM, 24 h), (**B**) 6Yn-Pro and NRH (500 µM, 24 h) co-treatment, (**C**) 6Yn-Pro followed by H_2_O_2_ treatment (in PBS, 2 mM, 6 min) and (**D**) 6Yn-Pro and NRH co-treatment followed by H_2_O_2_ treatment. Highlighted lines in red show PARP1 enrichment. (**E-H**) Volcano plots showing protein labelling and enrichment, and their statistical significance, for one treatment condition relative to another treatment condition. All samples were co-treated with NRH (500 µM, 24 h) followed by H_2_O_2_ treatment (in PBS, 2 mM, 6 min) prior to cell lysis. Significantly enriched known ADP-ribosylated proteins are shown in red, HSPA5 is shown in purple and all other significantly enriched proteins are shown in blue. Comparison of protein labelling in (**E**) 6Yn-Pro treated samples relative to DMSO control, (**F**) 6Yn-Pro treated samples relative to 6Yn-Pro and Olaparib co-treated samples, (**G**) 6Yn-Pro and Olaparib co-treated samples relative to DMSO and Olaparib co-treated samples and (**H**) 6Yn-Pro treated samples relative to DMSO control, following NMNAT1 overexpression (24 h).

We next co-treated HEK293T cells with 6Yn-Pro and NRH and included an H_2_O_2_ treatment step prior to cell lysis in triplicate and quantified significantly enriched proteins in probe-treated vs DMSO-treated samples (**Figure 4E**).

A total of 22 proteins were significantly enriched (**Table S3**), 8 of which are known ADP-ribosylated proteins, including PARP1 (**Figure 4E**), by comparison to the ADP-ribosylation database generated by Buch-Larsen *et al*.^45^ (see **Supplementary File Table S3** in Buch-Larsen *et al*.^45^; derived from the ADP-ribosylation (ADPr) data sets identified in the following 7 ADPr studies^45-51^) and Hendriks *et al*.^52^ (see **Supplementary File Data 7** in Hendriks *et al*.^52^). A comparison of probe-treated vs probe and Olaparib (O; PARP1/2 inhibitor) co-treated samples revealed that PARP1 was the only significantly enriched protein in the absence of Olaparib (**Table S4**). This suggested that the other 7 significantly enriched known ADP-ribosylated proteins (in orange/red) may be ADP-ribosylation targets of other PARPs, and the remaining 14 proteins (in blue) are potentially AMPylated proteins (**Figure 4F**). In samples co-treated with Olaparib, we observed the appearance of HSPA5 (**Table S5**), a known AMPylated protein^53^, as a significantly enriched protein (**Figure 4G**), which was not enriched in the absence of Olaparib. A possible explanation for this observation is that protein AMPylation (*via* 6Yn-ATP) is more likely when poly-ADP-ribosylation (and 6Yn-NAD^+^ consumption) is largely inhibited in the presence of Olaparib. These data suggest that AMPylation and ADP-ribosylation may compete for intermediates in the NAD^+^ synthesis pathway and illustrate the significance of optimizing experimental conditions that can favor production of modified NAD^+^ cofactor and consequently protein ADP-ribosylation over AMPylation, when using an upstream precursor probe such as 6Yn-Pro.

### Protein ADP-ribosylation levels negatively correlate with high NMNAT1 expression levels

Based on the earlier observation that NMNAT1 overexpression resulted in increased levels of 6Yn-NAD^+^ (**Figure 3C and S5D**), we investigated the effect of NMNAT1 overexpression (NM1) on protein labelling and enrichment. HEK293T cells were transfected with NMNAT1 plasmid for 24 h prior to probe-treatment and then co-treated with 6Yn-Pro (or DMSO) and NRH for 24 h. All samples were treated with H_2_O_2_ prior to cell lysis. Comparison of probe-treated vs DMSO-treated samples in the presence of NMNAT1 overexpression revealed that, compared to no NMNAT1 overexpression, only 12 (vs 22) proteins were significantly enriched, and only 5 (vs 8) of those were known ADP-ribosylated proteins (**Table S6**), including PARP1 (**Figure 4H**). Surprisingly, although the number of significantly enriched known ADP-ribosylated proteins decreased, the extent of PARP1 labelling and enrichment increased following NMNAT1 overexpression (Log_2_ Fold change = 4 vs 1.6 without NMNAT1 overexpression), suggesting increased PARP1 auto-ADP-ribosylation. The former observation was contrary to our expectation that higher 6Yn-NAD^+^ levels observed following NMNAT1 overexpression (**Figure 3F and 3G**) would result in an increase in the number of significantly enriched known ADP-ribosylated proteins. However, the observation correlated well with overall MAR/PAR levels obtained by western blotting, which showed that the overall ADP-ribosylation levels decreased following NMNAT1 overexpression (**Figure S7**). These findings point to the presence of a negative feedback mechanism between high NMNAT1 expression and overall ADP-ribosylation. Optimization of NMNAT1 levels warrant future investigation to explore whether intermediate levels of NMNAT1 lead to increased PARP1 activation without negatively impacting overall ADP-ribosylation levels.

### 6Yn-Pro probe enables identification of ADP-ribosylated proteins under oxidative stress

Since H_2_O_2_ significantly upregulates ADP-ribosylation, we determined the impact of H_2_O_2_ treatment on the enrichment of ADP-ribosylated proteins in probe-treated samples. In probe- and H_2_O_2_-treated samples (**Figure 4H**), 35 known ADP-ribosylated proteins were significantly enriched (Log_2_ Fold change > 0.5; **Figure 5A and 5C**; **Tables S7 and S8**), 21 of which were also identified in the corresponding probe-treated samples in the absence of H_2_O_2_ treatment (**Figure 5B and 5D**; **Tables S9 and S10**). In the absence of H_2_O_2_, the enrichment of all 21 matching ADP-ribosylated proteins (highlighted in green) decreased, consistent with reduced ADP-ribosylation. By comparing the extent of enrichment of matching proteins in H_2_O_2_-treated and untreated samples, the same methodology could therefore be applied for the identification of novel ADP-ribosylation protein targets using the probe 6Yn-Pro.

**Figure 5.**
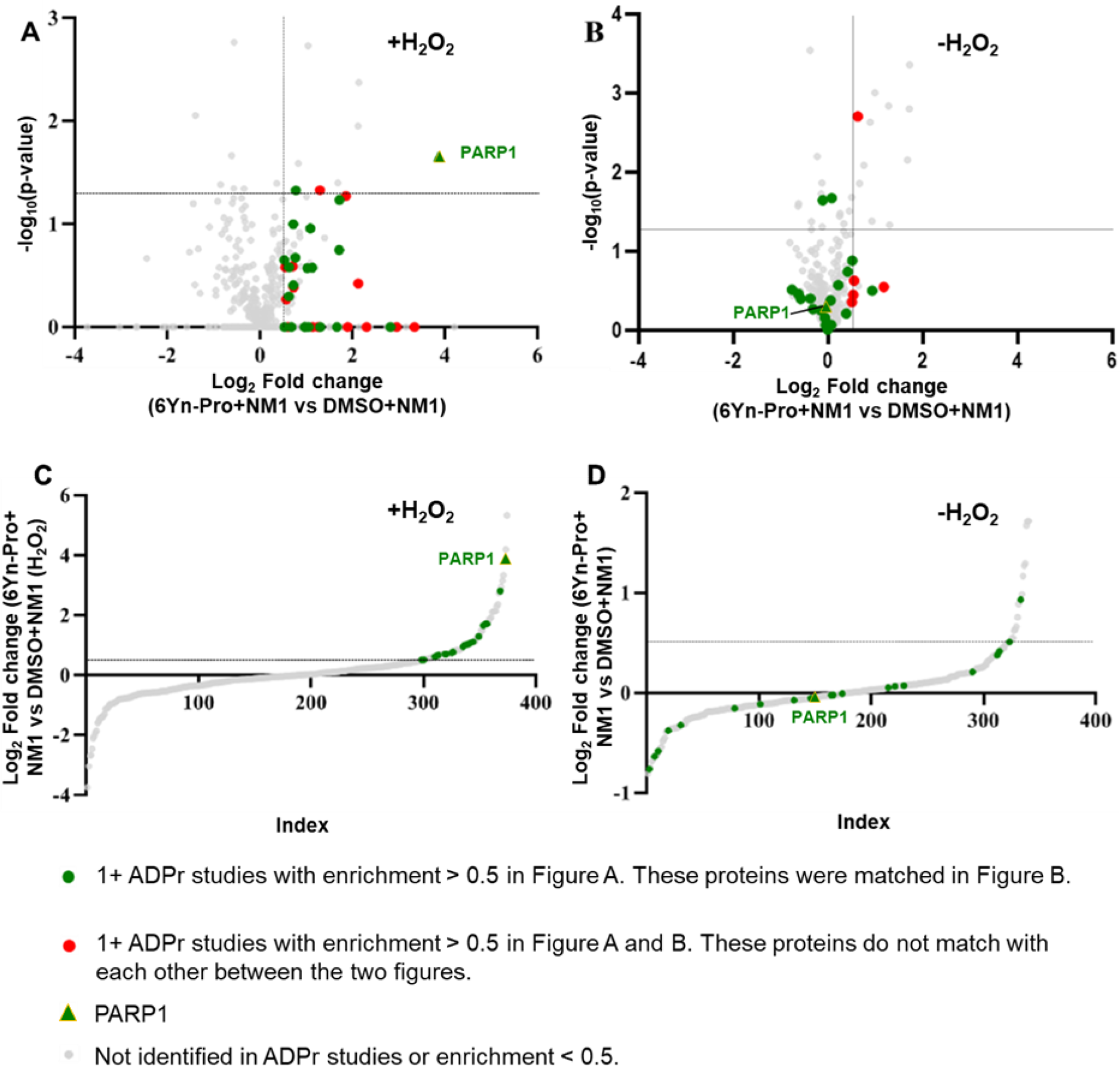
The presence of H_2_O_2_ upregulates labelling of ADPr proteins following treatment with 6Yn-Pro. Comparison of protein labelling profiles of 6Yn-Pro-treated HEK293T cells following NMNAT1 overexpression, (**A**) with H_2_O_2_ treatment and (**B**) without H_2_O_2_ treatment prior to cell lysis. All known ADP-ribosylated proteins with an enrichment value (Log_2_ Fold change) > 0.5 in **A** that matched the data set in **B** are highlighted in green. All known ADP-ribosylated proteins with an enrichment value (Log_2_ Fold change) > 0.5 in **A** that do not match the data set in **B**, and vice versa, are highlighted in red. (**C-D**) Comparison of the extent of enrichment of known ADP-ribosylated proteins in H_2_O_2_ treated samples and non-treated samples. (**C**) Data in volcano plot **A** visualized on an S-plot (significance is omitted). (**D**) Data in volcano plot **B** visualized on an S-plot (significance is omitted).

## CONCLUSION

To our knowledge, the present study is the most comprehensive to date on the metabolic incorporation of cell-permeable precursor probes for identification of intracellular ADP-ribosylation protein targets. We demonstrate successful metabolic incorporation of clickable precursor probe 6Yn-Pro in the NAD^+^ biosynthesis pathway for intracellular metabolic production of the ADP-ribosylation cofactor 6Yn-NAD^+^, describe several experimental conditions for boosting intracellular 6Yn-NAD^+^ levels, and show its subsequent use for labelling and enrichment of intracellular ADP-ribosylated proteins. We also identify a significant issue of non-specific labelling with nucleoside analogues which appears to have impacted previously reported studies and provide conditions for its removal. We anticipate that these probes and methodology will serve as a basis for further optimization of clickable NAD^+^ precursor probes for studying intracellular ADP-ribosylation events, including non-canonical ADP-ribosylation^39^.

## ASSOCIATED CONTENT

The supporting information is available free of charge at:

Methods and Materials used, NMR compound data, tables and additional figures, including chemical structures, western blots and plasmid maps (PDF)

Extended Data S1 (XLSX)

Extended Data S2 (XLSX)

### Notes

All authors declare no competing financial interest.

## ACKNOWLEDGMENT

The authors would like to thank L. Haigh at the Department of Chemistry, Imperial College London for managing the departmental Mass Spectrometry Facility, A. Coulson at the Department of Chemistry, Imperial College London for maintaining the departmental Tissue Culture Facility, and P. Haycock at the Department of Chemistry, Imperial College London for managing the departmental NMR Facility. This work was jointly funded by the British Heart Foundation (RE/13/4/30184) and the Engineering and Physical Sciences Research Council (EPSRC) through the Institute of Chemical Biology (EP/L015498/1). M.J.G and H.C.K acknowledge funding from EPSRC (EP/S023518/1) and Cancer Research UK (CRUK) (CANTAC721\100021). Work in the Tate laboratory was supported by CRUK Programme Award to EWT (DRCNPGNov21\100001 and C29637/A20183).

