## Supporting Information for "Chemical Tools for Profiling the Intracellular ADP-ribosylated Proteome"

This document includes:

Supplementary Methods

Figures S1 to S7

Tables S1 to S10

1-((2R,3R,4S,5R)-3,4-dihydroxy-5-(hydroxymethyl)tetrahydrofuran-2-yl)-1,4-dihydropyridine-3-carboxamide (NRH) ... 10
Synthesis of Adenosine ProTide Proprobe Analogues ... 10
Benzyl ((((2R,3S,4R,5R)-3,4-dihydroxy-5-(6-(prop-2-yn-1-ylamino)-9H-purin-9-yl)tetrahydrofuran-2-yl)methoxy)(phenoxy)phosphoryl)-L-alaninate (6Yn-Pro) ... 11
Benzyl ((((2R,3S,4R,5R)-5-(6-amino-2-ethynyl-9H-purin-9-yl)-3,4-dihydroxy tetrahydrofuran-2-yl)methoxy)(phenoxy)phosphoryl)-L-alaninate (2Yn-Pro) ... 11
Benzyl ((((2R,3S,4R,5R)-5-(6-amino-8-(but-3-yn-1-ylthio)-9H-purin-9-yl)-3,4-dihydroxytetrahydrofuran-2-yl)methoxy)(phenoxy)phosphoryl)-L-alaninate (8But-Yn-Pro) ... 12
Synthesis of Phosphorylated Adenosine (Nucleotide) Analogues ... 12
((2R,3S,4R,5R)-3,4-dihydroxy-5-(6-(prop-2-yn-1-ylamino)-9H-purin-9-yl)tetra hydrofuran-2-yl)methyl dihydrogen phosphate (6Yn-AMP) ... 12
((2R,3S,4R,5R)-3,4-dihydroxy-5-(6-(prop-2-yn-1-ylamino)-9H-purin-9-yl)tetra hydrofuran-2-yl)methyl tetrahydrogen triphosphate (6Yn-ATP) ... 13
3-carbamoyl-1-((2R,3R,4S,5R)-5-(((((((2R,3S,4R,5R)-3,4-dihydroxy-5-(6-(prop-2-yn-1-ylamino)-9H-purin-9-yl)tetrahydrofuran-2-yl)methoxy)(hydroxy)phosphoryl) oxy)(hydroxy)phosphoryl)oxy)methyl)-3,4-dihydroxytetrahydrofuran-2-yl)pyridin-1-ium (6Yn-$NAD^+$) ... 13
Biological Methods ... 14
Sample Preparation for In-gel Fluorescence and Western Blotting ... 14
Sample Preparation for Proteomics ... 16
Analysis of Proteomics Samples ... 17
Data Processing and Data Analysis ... 17
Sample Preparation for Metabolomics ... 18
Cloning of NMNAT1 Sequence and NMNAT1 Overexpression ... 19
NMR Spectra ... 35
$^1$H NMR of N-(prop-2-yn-1-yl)-9H-purin-6-amine (6Yn-adenine) ... 35
$^{13}$C NMR of N-(prop-2-yn-1-yl)-9H-purin-6-amine (6Yn-adenine) ... 36
$^1$H NMR of (2R,3S,4R,5R)-2-(hydroxymethyl)-5-(6-(prop-2-yn-1-ylamino)-9H-purin-9-yl)tetrahydrofuran-3,4-diol (6Yn-Ad) ... 37
$^{13}$C NMR of (2R,3S,4R,5R)-2-(hydroxymethyl)-5-(6-(prop-2-yn-1-ylamino)-9H-purin-9-yl)tetrahydrofuran-3,4-diol (6Yn-Ad) ... 38
$^1$H NMR of (2R,3R,4S,5R)-2-(6-amino-2-ethynyl-9H-purin-9-yl)-5-(hydroxymethyl)tetrahydrofuran-3,4-diol (2Yn-Ad) ... 39
$^{13}$C NMR of (2R,3R,4S,5R)-2-(6-amino-2-ethynyl-9H-purin-9-yl)-5-(hydroxymethyl)tetrahydrofuran-3,4-diol (2Yn-Ad) ... 40
$^1$H NMR of (2R,3R,4S,5R)-2-(6-amino-8-bromo-9H-purin-9-yl)-5-(hydroxymethyl)tetra hydrofuran-3,4-diol (8Br-Ad) ... 41

$^{13}$C NMR of (2R,3R,4S,5R)-2-(6-amino-8-bromo-9H-purin-9-yl)-5-(hydroxymethyl)tetra hydrofuran-3,4-diol (8Br-Ad) ... 42

$^{1}$H NMR of (2R,3R,4S,5R)-2-(6-amino-8-(but-3-yn-1-ylthio)-9H-purin-9-yl)-5-(hydroxy methyl)tetrahydrofuran-3,4-diol (8But-Yn-Ad) ... 43

$^{13}$C NMR of (2R,3R,4S,5R)-2-(6-amino-8-(but-3-yn-1-ylthio)-9H-purin-9-yl)-5-(hydroxy methyl)tetrahydrofuran-3,4-diol (8But-Yn-Ad) ... 44

$^{1}$H NMR of 3-carbamoyl-1-((2R,3R,4R,5R)-3,4-diacetoxy-5-(acetoxymethyl)tetrahydrofuran-2-yl)pyridin-1-ium (NR-OAc) ... 45

$^{13}$C NMR of 3-carbamoyl-1-((2R,3R,4R,5R)-3,4-diacetoxy-5-(acetoxymethyl)tetrahydrofuran-2-yl)pyridin-1-ium (NR-OAc) ... 46

$^{1}$H NMR of 1-((2R,3R,4S,5R)-3,4-dihydroxy-5-(hydroxymethyl)tetrahydrofuran-2-yl)-1,4-dihydropyridine-3-carboxamide (NRH) ... 47

$^{13}$C NMR of 1-((2R,3R,4S,5R)-3,4-dihydroxy-5-(hydroxymethyl)tetrahydrofuran-2-yl)-1,4-dihydropyridine-3-carboxamide (NRH) ... 48

$^{1}$H NMR of Benzyl ((((2R,3S,4R,5R)-3,4-dihydroxy-5-(6-(prop-2-yn-1-ylamino)-9H-purin-9-yl)tetrahydrofuran-2-yl)methoxy)(phenoxy)phosphoryl)-L-alaninate (6Yn-Pro) ... 49

$^{13}$C NMR of Benzyl ((((2R,3S,4R,5R)-3,4-dihydroxy-5-(6-(prop-2-yn-1-ylamino)-9H-purin-9-yl)tetrahydrofuran-2-yl)methoxy)(phenoxy)phosphoryl)-L-alaninate (6Yn-Pro) ... 50

$^{31}$P NMR of Benzyl ((((2R,3S,4R,5R)-3,4-dihydroxy-5-(6-(prop-2-yn-1-ylamino)-9H-purin-9-yl)tetrahydrofuran-2-yl)methoxy)(phenoxy)phosphoryl)-L-alaninate (6Yn-Pro) ... 51

$^{1}$H NMR of Benzyl ((((2R,3S,4R,5R)-5-(6-amino-2-ethynyl-9H-purin-9-yl)-3,4-dihydroxy tetrahydrofuran-2-yl)methoxy)(phenoxy)phosphoryl)-L-alaninate (2Yn-Pro) ... 52

$^{13}$C NMR of Benzyl ((((2R,3S,4R,5R)-5-(6-amino-2-ethynyl-9H-purin-9-yl)-3,4-dihydroxy tetrahydrofuran-2-yl)methoxy)(phenoxy)phosphoryl)-L-alaninate (2Yn-Pro) ... 53

$^{31}$P NMR of Benzyl ((((2R,3S,4R,5R)-5-(6-amino-2-ethynyl-9H-purin-9-yl)-3,4-dihydroxy tetrahydrofuran-2-yl)methoxy)(phenoxy)phosphoryl)-L-alaninate (2Yn-Pro) ... 54

$^{1}$H NMR of Benzyl ((((2R,3S,4R,5R)-5-(6-amino-8-(but-3-yn-1-ylthio)-9H-purin-9-yl)-3,4-dihydroxytetrahydrofuran-2-yl)methoxy)(phenoxy)phosphoryl)-L-alaninate (8But-Yn-Pro) ... 55

$^{13}$C NMR of Benzyl ((((2R,3S,4R,5R)-5-(6-amino-8-(but-3-yn-1-ylthio)-9H-purin-9-yl)-3,4-dihydroxytetrahydrofuran-2-yl)methoxy)(phenoxy)phosphoryl)-L-alaninate (8But-Yn-Pro) ... 56

$^{31}$P NMR of Benzyl ((((2R,3S,4R,5R)-5-(6-amino-8-(but-3-yn-1-ylthio)-9H-purin-9-yl)-3,4-dihydroxytetrahydrofuran-2-yl)methoxy)(phenoxy)phosphoryl)-L-alaninate (8But-Yn-Pro) ... 57

### Supplementary Methods

#### Chemical Synthesis

General Synthetic Methods

All reagents and solvents were purchased from VWR, Sigma-Aldrich UK, Fluorochem, Tokyo Chemical Industry and Alfa Aesar and used without further purification, unless otherwise stated. All reactions under anhydrous conditions were performed under a nitrogen atmosphere in oven-dried glassware. All anhydrous solvents were obtained from the departmental SolvTM solvent towers (Innovative Technology Inc.) and freshly used. For reversed-phase purifications, HPLC grade acetonitrile (VWR, Merck) and ultrapure water from MilliQ Millipore purification system were used.

Analytical Techniques

Reaction monitoring for non-phosphorylated compounds was performed by thin-layer chromatography (TLC) on Merck aluminium plates pre-coated with silica gel 60 F254. Visualization of spots was performed either by using a UV lamp (at 254 nm) or by TLC plate staining using potassium permanganate, vanillin or ninhydrin. Monitoring of reactions with phosphorylated compounds was performed by LC-MS on a Waters 3100 Mass Detector system equipped with a photodiode array and an XBridge reverse phase C18 column (5 μm, 4.6 × 100 mm, operating at a flow rate of 1.2 mL/min). Unless otherwise stated, LC-MS analysis was performed using MeCN (solvent B) and $H_2O$ (solvent A) as the eluents (both containing 0.1% FA) over 18 mins using the following linear gradient: 5-98% MeCN (in $H_2O$)0-10 mins, 98% MeCN 10-12 mins, 98-5% 12-13 mins and 5% MeCN 13-18 mins. $^{1}H$ and $^{13}C$ NMR spectra were recorded on a 400 MHz Bruker AV NMR instrument at room temperature in chloroform-*d*, methanol-$d_4$, $D_2O$ or DMSO-*d*6 at 400 MHz and 101 MHz for $^{1}H$ and $^{13}C$ NMR respectively. NMR data is reported in the following manner: chemical shift (δ) recorded in ppm relative to trimethylsilane ($\delta_H$ and $\delta_C$ set at 0 ppm); multiplicity (s = singlet, d = doublet, t = triplet, q = quartet, m = multiplet); coupling constant (*J*) stated in Hz; number of protons (integration).

Purification Techniques

Purification was generally performed by manual column chromatography over normal phase silica gel 60 F254. For phosphorylated compounds, selected nucleosides and all ProTide analogues, purification was performed on a Biotage Isolera Prime automated system equipped with a reverse-phase Biotage SNAP Ultra C18 column (12 g, column volume = 17 mL). Unless otherwise stated, reverse-phase purification was performed using MeCN and $H_2O$ as the eluents (both containing 0.1% FA) over 25 column volumes (CV) at 12 mL/min using the following linear gradient: 5% MeCN (in $H_2O$) over 3 CV, 5-95% MeCN over 20 CV, 95% over 2 CV. AMP and ATP analogues were purified by anion exchange chromatography (AXC). AXC purifications were performed on a Biotage Isolera Prime automated system equipped with a SNAP cartridge (12 g, column volume = 17 mL), manually packed with activated DEAE Sephadex A-25 (chloride form) weak anion exchange resin. Activation of the resin was performed by dissolving the resin (7.5 g) in 1 M $NaHCO_{3\,(aq)}$ (100 mL) at 4 ˚C with mild shaking over 24 hours. The swollen resin was then washed with distilled $H_2O$ and packed into the empty SNAP cartridge described above. Unless otherwise stated, AXC purification was performed using 0.4 M $NH_4HCO_{3\,(aq)}$ and $H_2O$ as the eluents over 30 column volumes (CV) at 12 mL/min using the following linear gradient: 0-100% $NH_4HCO_{3\,(aq)}$ over 20 CV and then 100% $NH_4HCO_{3\,(aq)}$ over 10 CV. At the end, the column was flushed and the resin reactivated by washing subsequently with 2 M $NaCl_{(aq)}$ (100 mL), $H_2O$ (100 mL) and 1 M $NH_4HCO_{3\,(aq)}$ (50 mL) in that order, and the column was stored at 4 ˚C.

#### Synthesis of Adenosine Analogues and Nucleosides

**N-(prop-2-yn-1-yl)-9H-purin-6-amine (6Yn-adenine)**

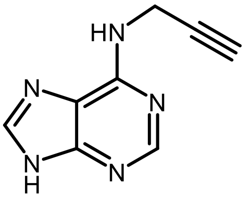

To a solution of 6-chloropurine (200 mg, 1.29 mmols) in EtOH (2 mL) was added propargylamine (414 μL, 6.47 mmols) and the reaction mixture was heated under reflux for 3 hours. The pale-yellow precipitate was filtered and washed with $H_2O$ and EtOH to afford **6Yn-adenine** as a pale-yellow powder (156 mg, 0.903 mmols, 70%). $R_f$ = 0.15 (DCM:MeOH 9:1); $^1$H NMR (400 MHz, DMSO-*d*$_6$) δ 13.01 (s, 1H), 8.26 (s, 1H), 8.15 (s, 1H), 8.02 (s, 1H), 4.28 (s, 2H), 3.04 (s, 1H); $^{13}$C NMR (101 MHz, DMSO-*d*$_6$) δ 154.2, 152.7, 150.0, 139.8, 119.3, 82.5, 72.9, 29.6; LC-MS (ESI$^+$) m/z found [M+H]$^+$ = 174.09 at $R_t$ = 2.33 mins.

**(2R,3S,4R,5R)-2-(hydroxymethyl)-5-(6-(prop-2-yn-1-ylamino)-9H-purin-9-yl)tetrahydrofuran-3,4-diol (6Yn-Ad)**

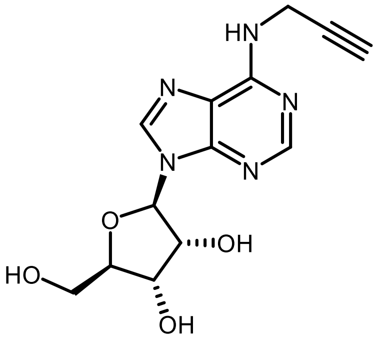

To a solution of 6-chloropurine riboside (1.00 g, 3.49 mmols) in EtOH (10 mL) was added propargylamine (1.12 mL, 17.4 mmols) and the reaction mixture was heated under reflux for 1 hour. The yellow-orange precipitate was filtered and washed with $H_2O$ and EtOH to afford **6Yn-Ad** as a white powder (799 mg, 2.61 mmols, 75%). $R_f$ = 0.2 (DCM:MeOH 9:1); $^1$H NMR (400 MHz, DMSO-*d*$_6$) δ 8.42 (s, 1H), 8.29 (s, 1H), 5.91 (d, *J* = 6.0, 1H), 5.49 (s, 1H), 5.37 (s, 1H), 5.23 (s, 1H), 4.61 (t, *J* = 5.5, 1H), 4.26 (s, 2H), 4.16 (t, *J* = 4.0, 1H), 3.97 (q, *J* = 3.5, 1H), 3.68 – 3.65 (m, 1H), 3.60 – 3.52 (m, 1H), 3.05 (s, 1H); $^{13}$C NMR (101 MHz, DMSO-*d*$_6$) δ 154.4, 152.7, 147.6, 140.7, 120.4, 88.3, 86.3, 82.3, 78.3, 74.0, 71.1, 62.1, 28.7; LC-MS (ESI$^+$) m/z found [M+H]$^+$ = 306.08 at $R_t$ = 7.55 mins; HRMS (ESI+) found 306.1193 ($C_{13}H_{15}N_5O_4$, [M+H]$^+$ requires 306.1202).

**(2R,3R,4S,5R)-2-(6-amino-2-((trimethylsilyl)ethynyl)-9H-purin-9-yl)-5-(hydroxy methyl)tetrahydrofuran-3,4-diol (2Yn-TMS-Ad)**

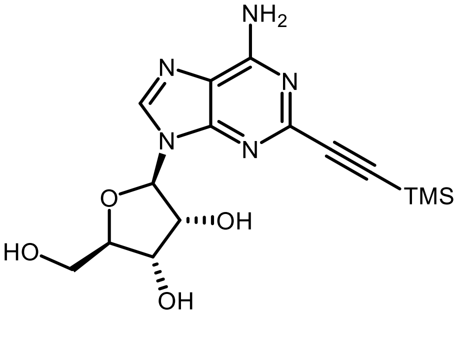

To a solution of 2-iodoadenosine (200 mg, 0.509 mmols) in dry and degassed DMF (2.5 mL) under nitrogen were added CuI (19.3 mg, 0.101 mmols), $Pd(PPh_3)_2Cl_2$ (35.6 mg, 0.0508 mmols) and dry and degassed trimethylamine (426 µL, 3.06 mmols). To the resulting dark orange solution degassed trimethylsilylacetylene (106 µL, 0.762 mmols) was added and the reaction mixture was stirred at r.t. for 16 hours. Upon addition of trimethylsilylacetylene, the reaction mixture immediately changed colour to clear yellow and gradually (over 30 mins) changed to black. Removal of solvent under reduced pressure afforded a black residue which was purified by manual column chromatography (DCM:MeOH 9:1) to afford **2Yn-TMS-Ad** as a dark yellow-brown residue. The presence of trace amounts and black inorganic impurities was noted and the crude product was used in the subsequent TMS deprotection reaction without further purification. $R_f$ = 0.5 (DCM:MeOH 9:1); LC-MS (ESI$^+$) m/z found $[M+H]^+$ = 364.36 at $R_t$ = 10.18 mins.

**(2R,3R,4S,5R)-2-(6-amino-2-ethynyl-9H-purin-9-yl)-5-(hydroxymethyl)tetrahydrofuran-3,4-diol (2Yn-Ad)**

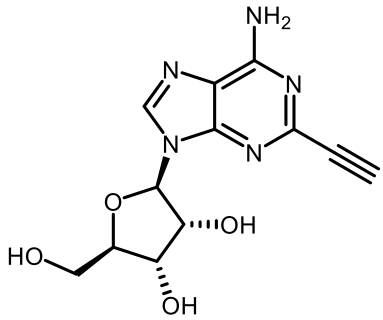

To a solution of **2Yn-TMS-Ad** in dry MeOH (3 mL) was added $K_2CO_3$ (491 mg, 3.56 mmols) and the reaction was stirred for 20 mins at r.t. (>30 mins reaction time leads to product degradation). The reaction mixture was diluted with DCM (27 mL) and purified (DCM:MeOH 9:1) to afford **2Yn-Ad** as a yellow powder (87 mg, 0.299 mmols, 59% over 2 steps). $R_f$ = 0.2 (DCM:MeOH 9:1); $^1$H NMR (400 MHz, DMSO-$d_6$) δ 8.44 (s, 1H), 7.54 (s, 2H), 5.87 (d, $J$ = 6.1, 1H), 5.48 (d, $J$ = 6.3, 1H), 5.20 (d, $J$ = 4.8, 1H), 5.19 (d, $J$ = 4.8, 1H), 4.56 (q, $J$ = 5.9, 1H), 4.14 (ddd, $J$ = 7.3 and 5.2 and 2.8, 1H), 4.03 (s, 1H), 3.96 (q, $J$ = 3.6, 1H), 3.68 (dt, $J$ = 12.0 and 4.4, 1H), 3.56 (ddd, $J$ = 12.1 and 6.8 and 3.8, 1H); $^{13}$C NMR (101 MHz, DMSO-$d_6$) δ 156.4, 149.6, 145.1, 141.2, 119.5, 88.0, 86.2, 83.7, 75.5, 74.1, 70.9, 61.9; LC-MS (ESI$^+$) m/z found $[M+H]^+$ = 292.03 at $R_t$ = 7.22 mins; HRMS (ESI$^+$) found 292.1040 ($C_{12}H_{16}N_8O_4$, $[M+H]^+$ requires 292.1046).

**(2R,3R,4S,5R)-2-(6-amino-8-bromo-9H-purin-9-yl)-5-(hydroxymethyl)tetra hydrofuran-3,4-diol (8Br-Ad)**

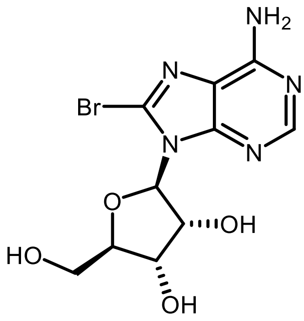

To a solution of 0.5 M NaOAc $_{(aq)}$ (22 mL) and AcOH (2.2 mL) was added adenosine (560 mg, 2.10 mmols) and the mixture was heated to 50 ˚C and stirred until complete dissolution of adenosine. Bromine solution (161 µL, 3.15 mmols) was then added to the reaction mixture at r.t. and the reaction was stirred at r.t. for 3 hours. The resulting dark red solution changed colour to yellow and was quenched by addition of excess sodium bisulphite, and neutralized (to pH 7) by the addition of 5M NaOH $_{(aq)}$ (~10 mL). The orange precipitate was filtered and washed with $H_2O$ and acetone to afford

**8Br-Ad** as a pale-yellow powder (253 mg, 0.730 mmols, 35%). $R_f$ = 0.15 (DCM:MeOH 9:1); $^1$H NMR (400 MHz, DMSO-*d*$_6$) δ 8.12 (s, 1H), 5.85 (d, *J* = 6.9, 1H), 5.08 (q, *J* = 6.9, 1H), 4.20 (dd, *J* = 5.2 and 2.2, 1H), 4.00 (q, *J* = 3.6, 1H), 3.69 (dd, *J* = 12.3 and 3.7, 1H), 3.53 (dd, *J* = 12.2 and 4.0, 1H); $^{13}$C NMR (101 MHz, DMSO-*d*$_6$) δ 155.5, 152.8, 150.3, 127.6, 120.1, 90.9, 87.2, 71.6, 71.3, 62.5; LC-MS (ESI$^+$) m/z found $[M+H]^+$ = 346.2, 348.2 (isotope) at $R_t$ = 7.12 mins.

**But-3-yn-1-yl carbamimidothioate**

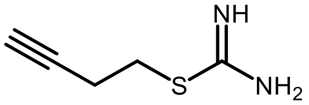

To a solution of 4-bromo-1-butyne (93.9 µL, 1.00 mmols) in dimethoxyethane (2.0 mL) was added thiourea (76.0 mg, 1.00 mmols) and the reaction mixture was heated at 50 ˚C for 16 hours. The solvent was evaporated under reduced pressure, the resulting yellow slurry was washed with $Et_2O$, and the crude product **but-3-yn-1-yl carbamimidothionate** was used in the next step without further purification. LC-MS (ESI$^+$) m/z found $[M+H]^+$ = 129.11 at $R_t$ = 1.96 mins.

**(2R,3R,4S,5R)-2-(6-amino-8-(but-3-yn-1-ylthio)-9H-purin-9-yl)-5-(hydroxy methyl)tetrahydrofuran-3,4-diol (8But-Yn-Ad)**

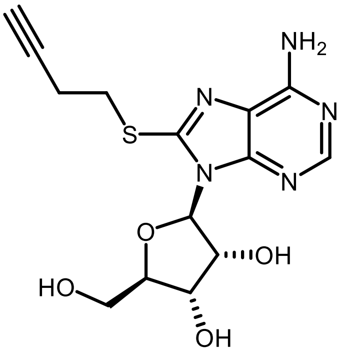

To a solution of 8-bromoadenosine **(8Br-Ad)** (100 mg, 0.289 mmols) in DMF (2 mL) was added a solution of **but-3-yn-1-yl carbamimidothioate** (76.9 mg, 0.6 mmols) in $H_2O$ (600 µL). 5 M NaOH $_{(aq)}$ (115.6 µL, 0.578 mmols) was added and the reaction mixture was stirred at 40 ˚C for 2 hours. The reaction was neutralized by addition of formic acid and the crude mixture was purified by automated reverse-phase column chromatography as described in section Purification Techniques. Removal of solvent under reduced pressure, followed by lyophilization afforded **8But-Yn-Ad** as a white powder (51 mg, 0.144 mmols, 50%). $R_f$ = 0.2 (DCM:MeOH 9:1); $^1$H NMR (400 MHz, DMSO-*d*$_6$) δ 8.07 (s, 1H), 7.35 (s, 2H), 5.75 (d, *J* = 6.9, 1H), 5.68 (d, *J* = 8.8, 1H), 5.46 (d, *J* = 6.3, 1H), 5.24 (d, *J* = 4.4, 1H), 5.00 (q, *J* = 5.9, 1H), 4.16 (t, *J* = 4.3, 1H), 3.98 (q, *J* = 3.4, 1H), 3.68 (dt, *J* = 12.4 and 3.7, 1H), 3.56 – 3.44 (m, 1H, 3), 2.97 (t, *J* = 2.7, 1H), 2.71 (td, *J* = 7.0 and 2.7, 2H); $^{13}$C NMR (101 MHz, DMSO-*d*$_6$) δ 155.1, 151.8, 150.9, 148.6, 120.1, 89.3, 87.1, 82.9, 73.3, 71.8, 71.5, 62.7, 31.6, 19.3; LC-MS (ESI$^+$) m/z found $[M+H]^+$ = 352.14 at $R_t$ = 7.23 mins; HRMS (ESI$^+$) found 352.1078 ($C_{14}H_{17}N_5O_4S$, $[M+H]^+$ requires 352.1079).

**3-carbamoyl-1-((2R,3R,4R,5R)-3,4-diacetoxy-5-(acetoxymethyl)tetrahydrofuran-2-yl)pyridin-1-ium (NR-OAc)**

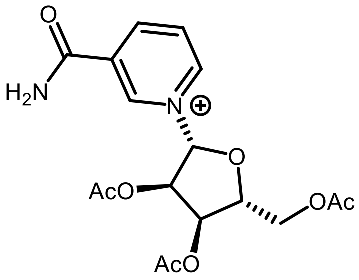

To a solution of nicotinamide (208 mg, 1.65 mmols) in dry MeCN (11 mL) under nitrogen was added β-D-ribofuranse 1,2,3,5-tetraacetate (350 mg, 1.10 mmols) followed by dropwise addition of TMS-OTf (239 µL, 1.32 mmols) over 10 mins and the reaction mixture was stirred at r.t. for 24 hours. The solvent was removed under reduced pressure, the crude residue was dissolved in MeOH (25 mL) and extracted with hexane (3 × 25 mL). Removal of the MeOH layer under reduced pressure afforded a pale-yellow residue which was dissolved in $H_2O$ (3 mL) and purified by automated reverse-phase column chromatography as described in section Purification Techniques. Removal of solvent under reduced pressure, followed by lyophilization afforded **NR-OAc** as a fluffy white powder (218 mg, 0.572 mmols, 52%). $^1H$ NMR (400 MHz, $D_2O$) δ 9.40 (t, $J$ = 1.6, 1H), 9.16 (dt, $J$ = 6.6 and 1.4, 1H), 8.94 (dt, $J$ = 8.1 and 1.5, 1H), 8.22 (dd, $J$ = 8.1 and 6.3, 1H), 6.54, (d, $J$ = 3.9, 1H), 5.52 (dd, $J$ = 5.4 and 3.8, 1H), 5.40, (t, $J$ = 5.4, 1H), 4.84 (dt, $J$ = 5.4 and 2.7, 1H), 4.48 (t, $J$ = 2.5, 2H), 2.11 (s, 3H), 2.08 (s, 3H), 2.04 (s, 3H); $^{13}C$ NMR (101 MHz, $D_2O$) δ 173.3, 172.4, 172.4, 165.5, 146.2, 143.1, 140.4, 134.2, 128.6, 97.3, 82.6, 76.4, 69.4, 62.6, 20.1, 19.8, 19.8; LC-MS (ESI$^+$) m/z found $[M+H]^+$ = 381.25 at $R_t$ = 7.31 mins.

**(2R,3R,4R,5R)-2-(acetoxymethyl)-5-(3-carbamoylpyridin-1(4H)-yl)tetrahydro furan-3,4-diyl diacetate (NRH-OAc)**

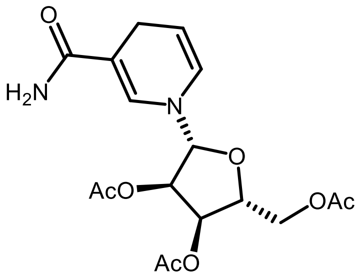

To a solution of **NR-OAc** (800 mg, 2.10 mmols) in $H_2O$ (15 mL) was added $NaHCO_3$ (1.15 g, 13.7 mmols) followed by sodium dithionite (915 mg, 5.26 mmols). The reaction mixture was stirred at r.t. for 2 mins, EtOAc (20 mL) was then added and the reaction mixture was stirred at r.t. for 2 hours. The EtOAc layer was extracted, followed by extraction of the aqueous layer with EtOAc (3 × 25 mL) and all EtOAc layers were combined. Acetone (25 mL) was added to completely dissolve the product, due to borderline solubility of the desired product in EtOAc, and the combined EtOAc and acetone layers were dried over anhydrous $Na_2SO_4$, filtered and evaporated under reduced pressure to afford crude **NRH-OAc** as a yellow residue. The product was used in the subsequent reaction step without further purification. LC-MS (ESI$^+$) m/z found $[M+H]^+$ = 383.01 at $R_t$ = 9.93 mins.

**1-((2R,3R,4S,5R)-3,4-dihydroxy-5-(hydroxymethyl)tetrahydrofuran-2-yl)-1,4-dihydropyridine-3-carboxamide (NRH)**

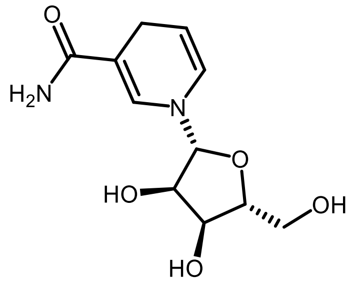

Crude **NRH-OAc** (assuming quantitative yield ~800 mg, 2.10 mmols) was dissolved in 4 M ammonia in MeOH solution (17 mL) and stirred at 4 ˚C for 20 hours. Removal of solvent under reduced pressure afforded a yellow residue which was dissolved in $H_2O$ (2 mL) and purified by automated reverse-phase column chromatography, eluting with MeCN and 25 mM $NH_4HCO_{3\ (aq)}$ over 25 CV at 12 mL/min using the following linear gradient: 2-5% MeCN (in 25 mM $NH_4HCO_{3\ (aq)}$) over 3 CV, 5-70% MeCN over 20 CV and 70% MeCN over 2 CV. The use of basic pH (>7) for the elution gradient was necessary to prevent the product from hydrating at position **4**, which occurs at pH <7. Removal of solvent under reduced pressure, followed by lyophilization afforded **NRH** as a bright yellow powder (56 mg, 0.219 mmols, 5% over three reaction steps). $^1$H NMR (400 MHz, DMSO) δ 6.98 (d, *J* = 1.3, 1H), 6.67 (s, 2H), 6.11 (dd, *J* = 8.3 and 1.8, 1H), 4.73 – 4.65 (m, 1H), 4.60 (d, *J* = 6.7, 1H), 4.16 – 3.92 (m, 3H), 3.92 – 3.86 (m, 1H), 3.83 (dd, *J* = 5.4 and 2.6, 1H), 3.70 – 3.64 (m, 1H), 3.51 – 3.40 (m, 2H), 2.95 (d, *J* = 3.4, 2H); $^{13}$C NMR (101 MHz, $D_2O$) δ 169.6, 136.3, 127.1, 102.9, 102.1, 95.7, 84.4, 71.7, 70.8, 62.3, 23.3; LC-MS (ESI$^+$) m/z found $[M+H]^+$ = 257.13 at $R_t$ = 2.00 mins.

#### Synthesis of Adenosine ProTide Proprobe Analogues

General Procedure for the Synthesis of ProTide Analogues

All ProTide analogues were synthesized according to the following general procedure: To a solution of 50 mg (1 eq.) of the corresponding adenosine analogue in freshly distilled trimethyl phosphate $PO(OMe)_3$ (500 µL) under nitrogen was added dry $NEt_3$ (4 eq.) at 0 ˚C. After 2 – 3 mins, phenyl dichlorophosphate (2 eq.) was added and the reaction mixture was stirred at 0 ˚C for 30 mins - 1 hour. Disappearance of the starting material and formation of the (hydrolyzed) activated intermediate was monitored by LC-MS. In a separate flask, to a solution of L-alanine benzyl ester hydrochloride (4 eq.) in freshly distilled $PO(OMe)_3$ (800 µL) under nitrogen was added dry $NEt_3$ (4 eq.) and the solution was transferred to the reaction mixture dropwise using a syringe. The flask was washed with $PO(OMe)_3$ (500 µL) and the washing was also transferred to the reaction mixture. The reaction was allowed to proceed at 0 ˚C for 10 mins and then at r.t. for 30 mins. The reaction was quenched with $H_2O$ (3 mL) and the resulting solution was purified by automated reverse-phase column chromatography as described in section Purification Techniques. Removal of solvent under reduced pressure, followed by lyophilization afforded all ProTide products as fluffy white powders.

**Benzyl ((((2R,3S,4R,5R)-3,4-dihydroxy-5-(6-(prop-2-yn-1-ylamino)-9H-purin-9-yl)tetrahydrofuran-2-yl)methoxy)(phenoxy)phosphoryl)-L-alaninate (6Yn-Pro)**

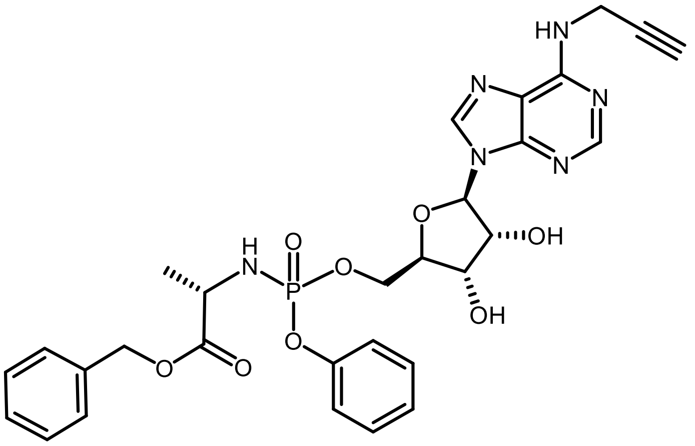

Synthesized according to the general procedure described. **6Yn-Pro** (23 mg, 0.0376 mmols, 23%). $R_f$ = 0.7 (DCM:MeOH 9:1); $^1$H NMR (400 MHz, methanol-$d_4$) δ 8.31 (s, 1H), 8.28 (s, 1H), 8.24 (s, 1H), 7.35 – 7.27 (m, 7H), 7.22 – 7.16 (m, 3H), 6.06 (t, *J* = 4.9, 1H), 5.12 (d, *J* = 3.3, 1H), 5.09 (d, *J* = 7.5, 1H), 4.65 (dt, *J* = 12.8 and 5.0, 1H), 4.44 – 4.37 (m, 4H), 4.37 – 4.29 (m, 1H), 4.25 (q, *J* = 3.4, 1H), 4.03 – 3.89 (m, 1H), 2.66 (t, *J* = 2.9, 1H), 1.30 (dd, *J* = 19.8 and 7.1, 3H); $^{13}$C NMR (101 MHz, methanol-$d_4$) δ 174.8, 155.1, 153.4, 152.0, 141.1, 137.2, 130.8, 129.6, 129.3, 129.3, 126.2, 121.4, 121.4, 121.0, 90.0, 84.4, 81.0, 75.4, 72.4, 71.5, 67.9, 67.1, 51.7, 30.9, 20.2; 31P NMR (162 MHz, methanol-$d_4$) δ 3.91, 3.65; LC-MS (ESI$^+$) m/z found [M+H]$^+$ = 623.24 at $R_t$ = 11.28 mins; HRMS (ESI$^+$) found 623.2014 ($C_{29}H_{31}N_6O_8P$, [M+H]$^+$ requires 623.2019).

**Benzyl ((((2R,3S,4R,5R)-5-(6-amino-2-ethynyl-9H-purin-9-yl)-3,4-dihydroxy tetrahydrofuran-2-yl)methoxy)(phenoxy)phosphoryl)-L-alaninate (2Yn-Pro)**

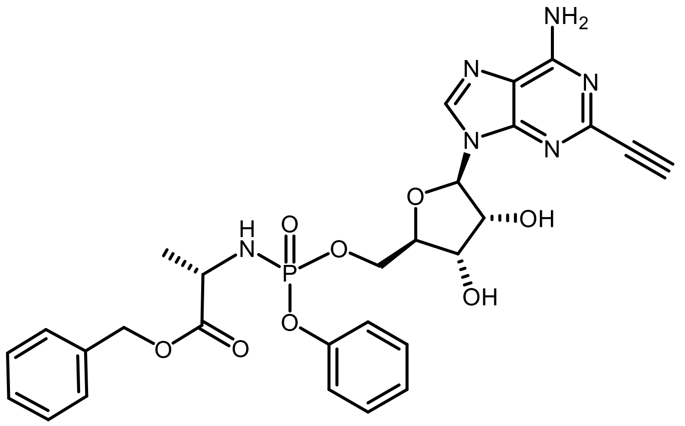

Synthesized according to the general procedure described. **2Yn-Pro** (2.1 mg, 0.00345 mmols, 2%). $R_f$ = 0.7 (DCM:MeOH 9:1); $^1$H NMR (500 MHz, methanol-$d_4$) δ 8.29 (s, 1H), 8.25 (s, 1H), 7.30 – 7.22 (m, 8H), 7.21 – 7.12 (m, 3H), 6.00 (d, *J* = 4.6, 1H), 5.10 (d, *J* = 6.0, 1H), 5.08 – 5.05 (m, 1H), 4.55 (dt, *J* = 14.8 and 4.9, 1H), 4.39 – 4.34 (m, 2H), 4.33 – 4.26 (m, 1H), 4.25 – 4.19 (m, 1H), 4.03 – 3.91 (m, 1H), 1.32 – 1.24 (m, 3H); $^{13}$C NMR (101 MHz, methanol-$d_4$) δ 173.6, 155.5, 150.6, 149.1, 140.5, 135.9, 129.4, 128.1, 128.0, 127.9, 124.8, 120.1, 120.0, 118.9, 88.7, 82.8, 74.2, 69.9, 66.5, 66.5, 66.0, 65.6, 50.0, 22.8; 31P NMR (162 MHz, methanol-$d_4$) δ 3.91, 3.66; LC-MS (ESI$^+$) m/z found [M+H]$^+$ = 609.25 at $R_t$ = 10.97 mins; HRMS (ESI$^+$) found 609.1863 ($C_{29}H_{31}N_6O_8P$, [M+H]$^+$ requires 609.1862).

**Benzyl ((((2R,3S,4R,5R)-5-(6-amino-8-(but-3-yn-1-ylthio)-9H-purin-9-yl)-3,4-dihydroxytetrahydro-furan-2-yl)methoxy)(phenoxy)phosphoryl)-L-alaninate (8But-Yn-Pro)**

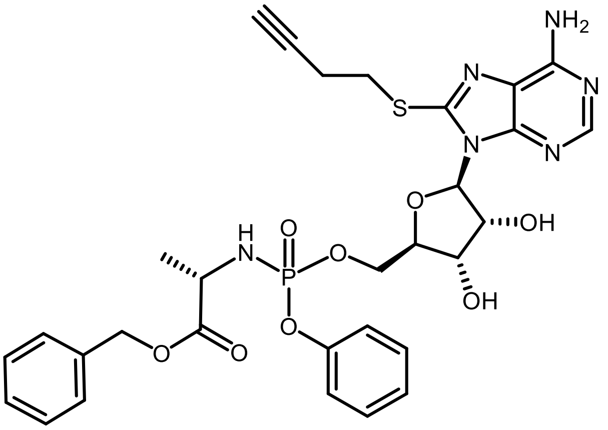

Synthesized according to the general procedure described. **8But-Yn-Pro** (8.0 mg, 0.0120 mmols, 8%). $R_f$ = 0.75 (DCM:MeOH 9:1); $^1$H NMR (500 MHz, methanol-$d_4$) δ 8.11 (s, 1H), 7.32 – 7.19 (m, 9H), 7.17 – 6.99 (m, 3H), 5.93 (d, $J$ = 4.7, 1H), 5.09 – 5.01 (m, 2H), 4.59 – 4.52 (m, 1H), 4.42 – 4.33 (m, 1H), 4.32 – 4.21 (m, 1H), 4.19 – 4.10 (m, 1H), 3.99 – 3.85 (m, 1H), 3.51 – 3.41 (m, 2H), 2.70 (td, $J$ = 7.1 and 2.6, 2H), 2.38 (t, $J$ = 2.6, 1H), 1.27 (dd, $J$ = 7.1 and 1.1, 3H); $^{13}$C NMR (101 MHz, methanol-$d_4$) δ 174.6, 155.2, 153.9, 152.3, 150.2, 137.2, 130.7, 129.6, 129.3, 129.3, 126.0, 121.4, 121.3, 121.0, 91.1, 84.4, 82.6, 73.0, 71.6, 71.5, 67.9, 67.6, 51.5, 32.5, 24.2, 20.2; 31P NMR (162 MHz, methanol-$d_4$) δ 3.53, 3.42; LC-MS (ESI$^+$) m/z found [M+H]$^+$ = 669.23 at $R_t$ = 11.16 mins; HRMS (ESI$^+$) found 669.1911 ($C_{30}H_{33}N_6O_8PS$, [M+H]$^+$ requires 669.1896).

#### Synthesis of Phosphorylated Adenosine (Nucleotide) Analogues

**((2R,3S,4R,5R)-3,4-dihydroxy-5-(6-(prop-2-yn-1-ylamino)-9H-purin-9-yl)tetra hydrofuran-2-yl)methyl dihydrogen phosphate (6Yn-AMP)**

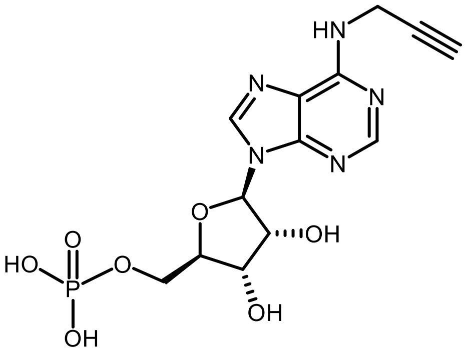

To a solution of **6Yn-Ad** (200 mg, 0.656 mmols) in freshly distilled $PO(OMe)_3$ (2.4 mL) under nitrogen was added dry DIPEA (340 μL, 1.96 mmols) at 0 ˚C. After 5 mins, $POCl_3$ (120 μL, 1.30 mmols) was added and the reaction mixture was stirred 0 ˚C for 3 hours. Dry DIPEA (85.0 μL, 0.490 mmols) and $POCl_3$ (30.0 μL, 0.325 mmols) were subsequently added and the reaction was further stirred at 0 ˚C for 1 hour. The reaction was quenched with $H_2O$ (7.5 mL) and the resulting solution was purified by automated reverse-phase column chromatography, eluting with MeCN and 25 mM $NH_4HCO_{3\,(aq)}$ over 23 CV at 12 mL/min using the following linear gradient: 2% MeCN (in 25 mM $NH_4HCO_{3\,(aq)}$) over 3 CV, 2-5% MeCN over 5 CV, 5-10% MeCN over 5 CV and 10-50% MeCN over 10 CV. Removal of solvent under reduced pressure, followed by lyophilization afforded **6Yn-AMP** as a white powder (90 mg, 0.235 mmols, 36%). $^1$H NMR (400 MHz, $D_2O$) δ 8.38 (s, 1H), 8.12 (s, 1H), 5.98 (d, $J$ = 5.8, 1H), 4.63 (t, $J$ = 5.4, 1H), 4.37 (dd, $J$ = 5.1 and 3.6, 1H), 4.28 – 4.22 (m, 1H), 4.21 – 4.13 (m, 2H), 3.98 – 3.87 (m, 2H), 2.49 (t, $J$ = 2.5, 1H); $^{13}$C NMR (101 MHz, $D_2O$) δ 153.8, 152.6, 148.3, 139.8, 119.0, 86.9, 84.3,

80.3, 74.4, 71.8, 70.5, 63.8, 30.1; $^{31}$P NMR (162 MHz, $D_2O$) δ 2.16; LC-MS (ESI$^+$) m/z found $[M+H]^+$ = 386.29 at $R_t$ = 2.7 mins; HRMS (ESI$^+$) found 386.0870 ($C_{13}H_{16}N_5O_7P$, $[M+H]^+$ requires 386.0787).

**((2R,3S,4R,5R)-3,4-dihydroxy-5-(6-(prop-2-yn-1-ylamino)-9H-purin-9-yl)tetra hydrofuran-2-yl)methyl tetrahydrogen triphosphate (6Yn-ATP)**

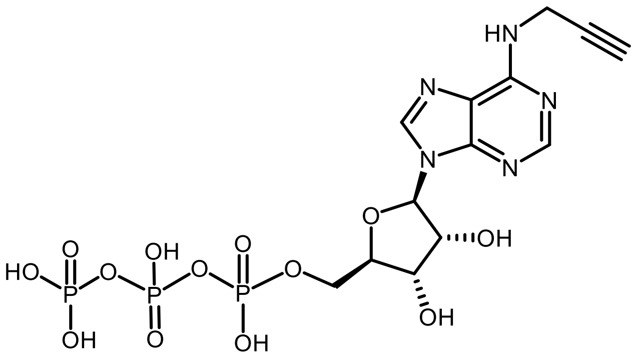

To a solution of **6Yn-Ad** (50 mg, 0.164 mmols) in freshly distilled $PO(OMe)_3$ (0.75 mL) under nitrogen was added dry DIPEA (85.6 µL, 0.493 mmols) at 0 ˚C. After 5 mins, $POCl_3$ (30.5 µL, 0.328 mmols) was added and the reaction mixture was stirred at 0 ˚C for 1 hour 45 mins. In a separate flask, to a solution of tributylammonium pyrophosphate (539 mg, 0.984 mmols) in freshly distilled $PO(OMe)_3$ (1.25 mL) under nitrogen was added dry DIPEA (85.6 µL, 0.493 mmols) and the solution was added dropwise to the reaction mixture at 0 ˚C over 5 mins. The reaction was allowed to proceed at 0 ˚C for 2 hours 30 mins and was quenched by the addition of 0.1 M $TEAB_{(aq)}$ (2 mL) at 0 ˚C for 30 mins. The resulting solution was purified by automated anion exchange chromatography as described in section Purification Techniques. Removal of solvent under reduced pressure, followed by lyophilization afforded **6Yn-ATP** as a white crystalline solid (19 mg, 0.0347 mmols, 21%). $^1$H NMR (400 MHz, $D_2O$) δ 8.26 (s, 1H), 8.03 (s, 1H), 5.91 (d, *J* = 5.5, 1H), 4.59 – 4.57 (m, 1H), 4.39 (t, *J* = 4.7, 1H), 4.25 – 4.18 (m, 1H), 4.11 – 4.07 (m, 3H), 3.90 (t, *J* = 13.4, 1H), 2.46 (s, 1H); $^{13}$C NMR (101 MHz, $D_2O$) δ 153.6, 152.5, 148.2, 139.5, 118.9, 86.8, 83.6, 80.3, 74.2, 71.8, 70.1, 65.1, 30.1; $^{31}$P NMR (162 MHz, $D_2O$) δ -10.68, -11.39, -23.01; LC-MS (ESI$^+$) m/z found $[M+H]^+$ = 546.1 at $R_t$ = 2.34 mins; HRMS (ESI$^+$) found 546.0170 ($C_{13}H_{18}N_5O_{13}P_3$, $[M+H]^+$ requires 546.0192).

**3-carbamoyl-1-((2R,3R,4S,5R)-5-(((((((2R,3S,4R,5R)-3,4-dihydroxy-5-(6-(prop-2-yn-1-ylamino)-9H-purin-9-yl)tetrahydrofuran-2-yl)methoxy)(hydroxy)phosphoryl) oxy)(hydroxy)phosphoryl)oxy)methyl)-3,4-dihydroxytetrahydrofuran-2-yl)pyridin-1-ium (6Yn-NAD$^+$)**

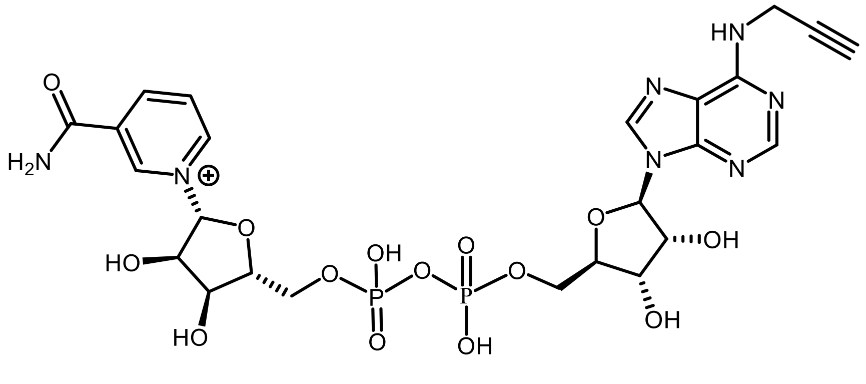

**6Yn-AMP** (3.8 mg, 0.0100 mmols) and β-nicotinamide mononucleotide (β-NMN) (10.2 mg, 0.0300 mmols) were added to separate flasks followed by addition of dry $NEt_3$ (1.0 mL) and stirred for 16 – 20 hours under nitrogen to afford the corresponding triethylammonium salts. After removal of $NEt_3$ under reduced pressure, to a solution of β-NMN.$NEt_3$ in dry DMF (210 µL) and dry 1,4-dioxane (150 µL) under nitrogen was added dry $NEt_3$ (30.0 µL, 0.216 mmols) followed by diphenyl phosphoryl chloride (12.5 µL, 0.0600 mmols) and benzyl tributyl ammonium chloride (9.4 mg, 0.0300 mmols). The yellow reaction mixture was stirred at r.t. for 3 hours and formation of the activated β-NMN intermediate was monitored by LC-MS (m/z $[M]^+$ 335). The reaction mixture was transferred into a 15 mL falcon tube, precipitated by addition of dry $Et_2O$ (10 mL) and centrifuged at 4000 g at 4 ˚C for 2 mins. A pale-yellow pellet formed at the bottom of the tube and the supernatant was discarded. Washing of the pellet was repeated with $Et_2O$, centrifuged at 4000 g at 4 ˚C for 2 mins and then dried under a flow of nitrogen for 15 mins and under vacuum for 3 hours. In a separate flask, a solution of **6Yn-AMP.NEt$_3$** in dry DMF (150 µL) under nitrogen was prepared and added to the dried pellet under nitrogen. A solution of dry pyridine (150 µL) in dry DMF (200 µL) under nitrogen was prepared and added to the reaction mixture, and the reaction was allowed to proceed at r.t. for 16 hours. The reaction mixture was transferred into a 15 mL falcon tube, precipitated by the addition of dry $Et_2O$ (12 mL) and centrifuged at 4000 g at 4 ˚C for 2 mins. A pale-yellow pellet formed at the bottom of the tube and the supernatant was discarded. Washing of the pellet was repeated with $Et_2O$, centrifuged at 4000 g at 4 ˚C for 2 mins and then dried under a flow of nitrogen for 15 mins to afford crude **6Yn-NAD$^+$** as a pale-yellow powder. Because of the highly polar nature of **6Yn-NAD$^+$**, it could not be successfully separated from other components in the reaction mixture, such as excess **6Yn-AMP**, and was not further purified. LC-MS ($ESI^+$) m/z found $[M+H]^+$ = 702.37 at $R_t$ = 2.2 mins. There was insufficient compound for full characterization.

#### Biological Methods

General Biological Methods

HEK293T and MDA-MB-231 cells were cultured in high glucose (4.5 g/L D-glucose, 4.0 mM L-glutamine, 1.0 mM sodium pyruvate) Dulbecco's Modified Eagle's Medium (DMEM) supplemented with 10% (v/v) Fetal Bovine Serum (FBS). T47D cells were cultured in high glucose (4.5 g/L D-glucose, 4.0 mM L-glutamine, 1.0 mM sodium pyruvate, 17.9 mM $NaHCO_3$, 10 mM HEPES) Roswell Park Memorial Institute (RPMI)-1640 medium supplemented with 10% (v/v) Fetal Bovine Serum (FBS). All cell types were cultured at 37 ˚C in a humidified 5% $CO_2$ incubator. Cells were passaged up to 30 times (for proteomics experiments) or up to 40 times for IGF and WB experiments.

**Sample Preparation for In-gel Fluorescence and Western Blotting**
For IGF and WB experiments with the ProTide analogues, typically 500 000 cells were seeded per well in a 6-well plate in 2 mL of the appropriate culture media for 16 – 20 h. After seeding, the media was removed and replaced with 1 mL of fresh culture media. 1 µL of the appropriate 100 mM ProTide stock (in DMSO), or DMSO only (negative control), was then added to the respective wells (to a final concentration of 100 µM) and treatment was performed for 24 h. After incubation, the culture media was aspirated, cells were washed twice with PBS and then lysed either with RIPA lysis buffer (25 mM Tris, 150 mM NaCl, 1% (v/v) Triton X100, 0.1% (w/v) SDS, 1% (w/v) sodium deoxycholate, protease inhibitor cocktail (1X, cOmplete), benzonase (50 units/mL of buffer, 10 mM olaparib, pH 7.6)) or 4% (w/v) SDS lysis buffer (50 mM HEPES, 4% (w/v) SDS, 150 mM NaCl, protease inhibitor (1X), benzonase (50 units/mL of buffer), 10 mM olaparib) on ice. Lysates were collected and cleared by centrifugation (16 000 g, 10 mins).

For samples where protein precipitation was performed after cell lysis, and prior to click chemistry conjugation, protein precipitation was performed in the following manner: MeOH (4 vol), $H_2O$ (2 vol) and chloroform $CHCl_3$ (1 vol) were sequentially added to each sample, where vol represents the volume of the lysis buffer. Samples were then centrifuged at 4 ˚C (for lysates generated in RIPA lysis buffer) or at r.t. (for lysates generated in 4% (w/v) SDS lysis buffer) for 5 mins at 6000 g. The supernatant (containing the MeOH/ $H_2O$ layer) was aspirated for each sample without disturbing the protein pellet (forming at the $CHCl_3$ – MeOH/$H_2O$ interface) and discarded. MeOH (500 µL) was added to each sample, the samples were sonicated to break up the pellets and the proteins were pelleted by centrifugation at r.t. (16 000 g, 5 mins). The supernatants were discarded and each protein pellet washed twice more with MeOH (500 µL per washing). The pellets were then air-dried, dissolved in 0.2% (w/v) SDS in PBS and protein concentrations were estimated in triplicates using Bio-Rad DC Protein Assay kit.

For samples where click chemistry conjugation by CuAAC was performed, a freshly prepared “click mix” was assembled by mixing the following stock reagents in a ratio 2:2:1:1 (v/v/v/v) – tris(2-carboxy-ethyl)phosphine hydroxide (TCEP, 50 mM in $H_2O$), copper sulphate ($CuSO_4$, 50 mM in $H_2O$), tris(benzyltriazolylmethyl)amine (TBTA, 10 mM in DMSO) and the appropriate capture reagent (AzT or AzTB (both prepared in-house), 10 mM in DMSO). In a typical click reaction, 6 µL of the “click mix” was added to every 100 µL of protein sample (at a 1 – 2 mg/mL protein concentration), and a minimum of 100 µg of protein sample were clicked per reaction. Samples were briefly vortexed and the click reaction was allowed to proceed for 2 – 3 hours at r.t. with shaking at 750 – 800 rpm. Quenching of the reaction was performed by the addition of 0.5 M EDTA $_{(aq)}$ stock to a final concentration of 5 mM, with shaking at r.t. for 2 mins. Protein precipitation was subsequently performed, as described in the previous paragraph, where vol represents the volume of the sample click reaction. The protein pellets were then dissolved in 0.2% (w/v) SDS in PBS to a final protein concentration of 1 mg/mL

The protein samples were prepared for gel electrophoresis by adding a loading buffer mixture, consisting of 4X (v/v) Laemmli loading buffer and 20% (v/v) β-mercaptoethanol, to 20 µg of protein sample in a ratio 3:1 (v/v) of protein sample:loading buffer mixture. Samples were then boiled by heating at 95 ˚C for 10 mins, allowed to cool down to r.t. and briefly centrifuged. Subsequently, the molecular weight ladder (MWL; Precision Plus Protein™ All Blue Prestained Protein Standards (Bio-Rad)) and the samples were loaded on a 10% or 12% sodium dodecyl sulphate-polyacrylamide (SDS-PAGE) gel, prepared in-house (with a 4% stacking gel on top), and the gel was run at 90 V for 10 mins and then at 150 V for 50 mins – 1 hour. Running of the gel was performed on a PowerPac™ Basic Power Supply (Bio-Rad) equipped with a Mini-PROTEAN Tetra Cell (Bio-Rad). The gel was briefly rinsed with $H_2O$ and subsequently imaged for TAMRA fluorescence ($\lambda_{abs}$ = 532 nm; $\lambda_{em}$ = 583 nm) on a Typhoon™ FLA 9500 imager (GE Healthcare). The gel was then either stained by InstantBlue Coomassie Protein Stain (Abcam) or Coomassie Brilliant Blue (Sigma-Aldrich), to check for protein loading, or used for western blotting as described in the next paragraph. Gels that were stained were imaged by trans-illumination on an ImageQuant LAS 4000 (Fujifilm) instrument and MWL was imaged by detecting for Cy5 ($\lambda_{abs}$ = 635 nm; $\lambda_{em}$ = 670 nm) fluorescence. Unless otherwise stated, all MWL values were shown in kDa units.

For western blotting, SDS-PAGE gels that were run (and not Coomassie stained) were transferred onto a nitrocellulose membrane. This was performed in a wet transfer buffer (25 mM Tris (base), 192 mM glycine, 20% (v/v) MeOH) for 1 hour at 100 V on a PowerPac™ Basic Power Supply (Bio-Rad) equipped with a Mini-PROTEAN Tetra Cell (Bio-Rad). To check for protein loading and confirm that the wet transfer was successful, the membrane was stained by Ponceau S stain (Sigma-Aldrich) for 5 mins at r.t. The membrane was then rinsed with $H_2O$ and the membrane was blocked in 5% (w/v) BSA in TBS-T (1X Tris-buffered saline, 0.1% (v/v) Tween-20) blocking buffer for 1 hour at r.t. with mild shaking. After 1 hour, the membrane was rinsed once with blocking buffer and incubated with primary antibody (appropriate dilution (v/v) of antibody stock in blocking buffer) for 16 – 20 hours at 4 ˚C. Following incubation, the membrane was washed with TBS-T (3 × 5 mins). The membrane was then either incubated with a secondary antibody (appropriate dilution (v/v) of antibody stock in blocking buffer) for 1 hour at r.t. (for non-HRP-linked primary antibodies), or directly imaged (for HRP-linked primary antibodies). After incubation with a secondary antibody the membrane was washed with TBS-T (3 × 5 mins) and then imaged. Imaging was performed by chemiluminescence following addition of Immobilon Crescendo Western HRP onto the entire surface of the membrane on an ImageQuant LAS 4000 (Fujifilm) instrument. MWL was imaged by detecting for Cy5 ($\lambda_{abs}$ = 635 nm; $\lambda_{em}$ = 670 nm) fluorescence. Unless otherwise stated, all MWL values were shown in kDa units. List of used antibodies: α-Poly/Mono-ADP ribose (anti-MAR/PAR antibody #83732, dilution 1:1000 (v/v), Cell Signaling Technology); α-pan-ADP-ribose binding reagent (MABE1016, dilution 1:1000 (v/v), Sigma-Aldrich); α-PARP1 (sc-8007, dilution 1:1000 (v/v), Santa Cruz Biotechnology); α-PARP1 (#9542, dilution 1:1000 (v/v), Cell Signaling Technology); α-NMNAT1 (sc-271557, dilution 1:500 (v/v), Santa Cruz Biotechnology); α-AMPylated tyrosine (ABS184, dilution 1:1000 (v/v), Sigma-Aldrich); anti-β-actin (A5316, dilution 1:10 000 (v/v), Sigma-Aldrich); α-mouse-HRP (secondary antibody R-05071-500, dilution 1:10 000 (v/v), Advansta); α-rabbit-HRP (secondary antibody R-05072-500, dilution 1:10 000 (v/v), Advansta).

For sample analysis by western blotting where protein enrichment (pull-down) was performed, protein pull-down was typically performed by incubating 200 µg (200 µL) of protein sample (0.2% (w/v) SDS in PBS pH 7.4, 1 mg/ml) with 20 µL streptavidin magnetic beads (Thermo Scientific Pierce™) for 3 hours at r.t., with shaking (750 – 800 rpm). Prior to pull-down incubation, the streptavidin magnetic beads were washed three times with 0.2% (w/v) SDS in PBS. Following pull-down incubation, the bead samples were placed on a magnetic rack, supernatants were transferred into clean Eppendorf tubes (the supernatants were then stored at -80 ˚C) and the bead samples were washed with 0.2% (w/v) SDS in PBS (3 × 500 µL). The washings were discarded and the enriched proteins were eluted from the beads by adding 15 µL of 2X Laemmli loading buffer and 10% (v/v) β-mercaptoethanol mixture to each bead sample and subsequent boiling by heating at 95 ˚C for 10 mins. Samples were allowed to cool down to r.t., briefly centrifuged and directly loaded on an SDS-PAGE gel for subsequent analysis by western blot.

**Sample Preparation for Proteomics**

All proteomics experiments were performed with HEK293T cells in biological triplicates, unless otherwise stated. Cell seeding was performed for 24 hours in 6 cm dishes at a concentration of $1 \times 10^6$ cells in 5 mL DMEM media (+10% (v/v/) FBS) per dish. For cells where NMNAT1 overexpression was induced, transfection with the CMV NMNAT1 plasmid (1000 ng per mL of media) was performed for 24 h, prior to probe treatment (see section **Cloning of NMNAT1 Sequence and NMNAT1 Overexpression**). Prior to metabolic labelling, the culture (or transfection) media was replaced with 3 mL of fresh media. 3 µL of the appropriate 100 mM ProTide stock (in DMSO), or DMSO only (negative control), was then added to the respective dish (to a final concentration of 100 µM) and treatment was performed for 24 h. For cells co-treated with NRH, 3 µL of a 500 mM NRH stock (in DMSO) was added to the appropriate dish at the same time as the ProTide and incubated for 24 h. After incubation, media was aspirated and cells were washed with PBS in the following manner: for cells (to be) treated with 2 mM $H_2O_2$ (in PBS), cells were washed once with PBS (5 mL) and subsequently incubated with 2 mM $H_2O_2$ (in PBS) for 6 mins prior to lysis. For cells not treated with 2 mM $H_2O_2$, cells were washed twice with PBS (5 mL) and directly lysed. Lysates were generated using RIPA lysis buffer and incubation on ice for 15 mins. Clearing of the lysates was performed by centrifugation at 13 000 g at 4 ˚C for 5 mins and protein precipitation of the lysates was performed with MeOH:$H_2O$:$CHCl_3$, as previously described (see previous section). Protein pellets were dissolved in 0.2% (w/v) SDS in PBS (500 µL per sample) and protein concentrations were measured using the Bio-Rad DC Protein Assay kit.

For proteomics samples, CuAAC click reaction was performed using 500 – 1000 µg of protein per sample (2 mg/mL final concentration) and AzB (Sigma-Aldrich, 762074) or AzTB (prepared in-house) for 2 hours at r.t., as described in the previous section. Following incubation and quenching of the reaction, protein precipitation was performed and pelleted proteins were dissolved in 0.2% SDS in PBS (500 µL or 1000 µL per sample, respectively, to a 1 mg/mL final concentration). Protein pull-down was then performed by incubating the 500 – 1000 µg (500 – 1000 µL, 1 mg/ml) of clicked protein sample (0.2% (w/v) SDS in PBS pH 7.4) with 50 – 100 µL Neutravidin agarose beads (Thermo Scientific Pierce™) for 3 hours at r.t., with shaking (750 – 800 rpm). Prior to the pull-down incubation, the Neutravidin agarose beads were washed five times with 0.2% (w/v) SDS in PBS, pelleted by centrifugation (3500 g, 2 mins) and supernatant aspirated by pipetting. Beads were suspended to their original volume and the appropriate volume of beads added to each sample (1:10 beads volume:sample volume). Following the incubation, the bead samples were pelleted by centrifugation (3500 g, 3 mins), supernatants aspirated, samples washed with 0.2% (w/v) SDS in 50 mM HEPES (pH 8.0) four times (1 mL each) and then washed three more times with 50 mM HEPES (pH 8.0). Each sample was subsequently made up to a volume of ~200 µL by addition of 150 µL 50 mM HEPES (pH 8.0) and on-bead sample reduction and alkylation was performed in the following manner: Stock solutions of TCEP (50 mM) and 2-chloroacetamide (CAA, 150 mM) were prepared in 50 mM HEPES (pH 8.0). Equal volumes of the stocks were combined and 40 µL of this mixture were added to each sample, giving final concentrations of 5 mM (TCEP) and 15 mM (CAA). Samples were then incubated at r.t. with shaking for 15 mins and peptides from each sample were released from the beads by digestion with trypsin. For trypsin digestion, 5 µL of trypsin stock (0.2 µg/µL) were added to each sample and subsequently incubated for 16 – 18 hours at 37 ˚C with shaking (900 – 1000 rpm). Digestion was quenched by the addition of EDTA-free protease inhibitor cocktail (4 µL, 50X) and incubation for 5 mins at r.t. with shaking. Beads were pelleted by centrifugation (3500 g, 3 mins) and supernatants were transferred into clean Eppendorf tubes. Each beads sample was washed once with 50 mM HEPES (pH 8.0, 100 µL), beads were pelleted (3500 g, 3 mins) and each washing combined with its respective supernatant from the first centrifugation step. Peptide concentration was measured for each sample using the Pierce™ Quantitative Fluorometric Peptide Assay (Thermo Scientific), in accordance with the manufacturer's protocol, and TMT-labelling was performed subsequently.

For the TMT-labelling reactions, 6 µg of each peptide sample were labelled and a different TMT10plex-channel was used for each treatment condition (biological triplicates were labelled with the same TMT-channel). TMT-channels were equilibrated to r.t., dissolved in anhydrous acetonitrile, one/fifth of the volume of each channel transferred into clean tubes (one/fifth of a TMT vial is enough to label ~20 µg of peptides) and the volume of each TMT-channel aliquot made up to 100 µL. TMT-labelling was then performed by adding 33 µL of a TMT-channel aliquot to each triplicate treatment condition (giving ~2:1 (v/v) ratio of sample volume:TMT-channel volume per labelling reaction) and incubating for 3 h at r.t. with shaking (750 rpm). Following incubation, the labelling reactions were quenched with 5% (w/v) hydroxylamine in 50 mM HEPES (pH 8.0, 2 µL per sample) and incubating for 15 mins at r.t. with shaking (750 rpm). All samples within a TMT set were combined, acidified by the addition of trifluoroacetic acid (2% (v/v) final concentration) and the combined samples were dried under vacuum. 4-Layer sample fractionation was subsequently performed using the Pierce™ High pH Reversed-Phase Fractionation Kit (Thermo Scientific) in accordance with the manufacturer's protocol, and using the following acetonitrile:triethylamine % gradients: fraction 1 – 12.5:87.5 (v/v); fraction 2 – 17.5:82.5; fraction 3 – 22.5:77.5; fraction 4 – 50:50. Following, fractionation, the samples were dried under vacuum. Finally, the dried fractionated peptide samples were dissolved in 15 µL of LC-MS grade H2O:MeCN:TFA (97.5:2:0.5 v/v/v) reagent mixture and filtered through a three-layer polyvinylidene fluoride (PVDF) Durapore 0.1 µm membrane (Millipore) directly into LC-MS polypropylene micro-vials (Kinesis) by centrifugation. The samples were then stored at 4 ˚C for same day or next day analysis.

**Analysis of Proteomics Samples**

Typically, 1 µg of peptides were loaded per sample injection on an EASY-Spray™ Acclaim PepMap $C_{18}$ column with an inner diameter of 50 × 75 µm. Separation of peptides was performed using a 3-hour linear gradient of 0-100% solvent B and a flow rate of 250 nL/min. The following solvent system was used: 80% MeCN supplemented with 0.1% FA (solvent B):2% MeCN supplemented with 0.1% FA (solvent A). Coupling of the LC to a QExactive MS instrument was performed *via* an EASY-Spray™ source (Thermo Fischer Scientific). Operation was set to data-dependent mode. Acquisition of survey scans was performed at a resolution of 70 000 at m/z 200 and in the range 350 – 1800 m/z. The most abundant isotope patterns (upper limit = 10) from the survey scan with a charge +2 or higher were selected for fragmentation by HCD. A 1.6 m/z isolation window was applied and a normalized collision energy for HCD fragmentation was set to 25. Survey scan maximum ion injection time was set to 20 ms. Acquisition of MS/MS scans was performed at a resolution of 35 000 at m/z 200 with a maximum ion injection time set to 120 ms. MS acquisition ion target value was set to $10^6$ and MS/MS acquisition ion target value was set to $10^5$.

**Data Processing and Data Analysis**

<u>Data Processing in MaxQuant</u>

Processing of mass spectrometry raw files was performed in MaxQuant (v1.6.5.0) for all experiments. Peptide searching was performed against human proteome reference list obtained as a FASTA file from UniProt (Taxonomy 9606, downloaded on 20 February 2020). The following were set as variable modifications: Oxidation (M), Acetyl (Protein N-term). The following were set as fixed modifications: Carbamidomethyl (C). Digestion mode was set to Trypsin (maximum missed cleavages = 2, minimum peptide length = 7 residues). For all analyses involving TMT-labelled samples, reporter ion $MS^2$ option was selected under ‘Type’ and the appropriate pre-defined TMT10plex labels were selected for lysine residues and N-termini. For protein quantification, ‘unique + razor’ peptide option was selected, with minimum label ratio count set to 2. All other parameters were applied as set by default.

Data Analysis in Perseus

Analysis of processed data from MaxQuant was performed in Perseus (v1.6.15.0) for all experiments. For TMT-labelling, all ‘reporter intensity corrected’ values for a given experiment were imported into Perseus from the ‘proteinGroups’ text file generated by MaxQuant. For experiments performed in triplicates, ‘reporter intensity corrected’ values were grouped (‘Categorical Annotation Rows’ option) into three categories: per TMT-set, per triplicates and per condition. Data filtering was performed by removing rows identified as ‘potential contaminants’ and ‘reverse’ (‘Filter Rows Based on Categorical Column’ option). $Log_2$ transformation was performed for all values (‘Basic -> Transform’ option). Normalization of protein abundance was performed by first subtracting the mean from each row in each TMT-set (‘Normalization -> Subtract -> Rows, TMT-set, Mean’) and then subtracting the median from each column (‘Normalization -> Subtract -> Columns, Median’). A final filtering step was applied to only include rows with at least two valid values (or at least one, where relevant) per triplicate condition (‘Filter Rows Based on Valid Values -> Number (set to 2), Mode -> In At Least One Group (set to Replicates)’). Summary statistics data was visualized in GraphPad Prism (v9.2.0). Processed data was visualized by either profile plots (in Perseus, log2 values plotted against their corresponding condition) or volcano plots (in GraphPad Prism (v9.2.0), significance (log(P)) plotted against difference in log2 values for two conditions) generated using Student’s t-test (paired two-sample t-test), with the following parameters set: False discovery rate (FDR) = 0.05, $S_o$ = 0.1, Number of randomizations = 250. For principal component analysis (PCA), missing values were replaced by ‘Imputation -> Replace Missing Values From Normal Distribution’.

**Sample Preparation for Metabolomics**

All metabolomics experiments were performed with HEK293T cells in biological triplicates, unless otherwise stated. Cell seeding was performed for 24 hours in 6-well dishes at a concentration of 400 000 cells per well in 2 mL DMEM media (+10% (v/v/) FBS). For cells where NMNAT1 overexpression was induced, transfection with the CMV NMNAT1 plasmid (1000 ng per mL of media) was performed for 24 h, prior to probe treatment (see section **Cloning of NMNAT1 Sequence and NMNAT1 Overexpression**). Prior to metabolic labelling, the culture (or transfection) media was replaced with 1 mL of fresh media. 1 µL of the appropriate 100 mM ProTide stock (in DMSO), or DMSO only (vehicle/negative control), was then added to the respective well (to a final concentration of 100 µM) and treatment was performed for 24 h. For cells treated with NRH, 1 µL of a 500 mM NRH stock (in DMSO) was added to the appropriate wells alone or at the same time as the ProTide (for co-treated cells) and incubated for 24 h. On the day of metabolite extraction, cell counts were performed on a Vi-CELL XR Cell Viability Analyzer (Beckman coulter). The remaining plates were transferred onto cooling blocks (cooled on ice), aspirated and washed once with 1 mL of cooled Ringers buffer (Sigma-Aldrich, 96724). After aspiration, 1 mL of cold 80:20 methanol:water (v/v) was added to each well. After 20 mins, cells were scrapped and transferred into 2 mL Eppendorf tubes. The previous step was repeated with 500 µL of 100% methanol, the washings were combined with their respective extraction samples, vortexed briefly and then centrifuged at 4 ˚C (18,000 g, 20 mins). 1 mL of each sample was transferred into a high recovery LC-MS vial, samples were dried by nitrogen flow and then stored at -80ºC until analysis. On the day of analysis, the dried samples were resuspended in 50 µL of proteomics grade $H_2O$ and 5 µL of a sample were loaded on the column per injection. Metabolic profiling and MS/MS analysis was carried out using a QTRAP4000 triple quadrupole mass spectrometer (AB Sciex, Dahaner Corporation) coupled to a 1290 Infinity UPLC system (Agilent Technologies). A 15 min normal-phase chromatographic method was used for both positive and negative ionization modes. For ATP and 6Yn-ATP molecules a HILIC (Hydrophilic Interaction Chromatography) method was used. LC separation was performed on an ACQUITY UPLC BEH amide column (Waters) with dimensions 3 × 150 mm and particle size 1.7 µm.

Column temperature was set to 40 ˚C. In $ESI^-$ mode, the following solvent system was used: 20 mM ammonium acetate and 10 mM ammonium hydroxide, both in $H_2O$ (solvent B):MeCN and 10 mM ammonium hydroxide in $H_2O$ (solvent A). Analysis was performed using a 15 mins gradient of 10-55% solvent B (10.0% 0-1 mins, 10.0-55.0% 1-8 mins, 55.0% 8-9 mins, 55.0-10.0% 9-9.10 mins, 10.0% 9.10-15 mins) and a flow rate of 500 µL/min. In $ESI^+$ mode, the following solvent system was used: 20 mM ammonium formate supplemented with 0.1% FA in $H_2O$ (solvent B):MeCN supplemented with 0.1% FA (solvent A). Analysis was performed using a 15 mins gradient of 5-50% solvent B (5.0% 0-1 mins, 5.0-50.0% 1-8 mins, 50.0% 8-9 mins, 50.0-5.0% 9-9.10 mins, 5.0% 9.10-15 mins) and a flow rate of 500 µL/min. Wash steps in-between sample analysis were performed using the same gradients. Data for the remaining metabolites was acquired using Reversed-Phase (RP) Liquid Chromatography. LC separation was performed on a ACQUITY UPLC® HSS (High Strength Silica) T3 Column (Waters) with dimensions 2.1 × 100 mm and particle size 1.8 µm. Column temperature was set to 40 ˚C. The following solvent system was used: 100% MeCN supplemented with 0.2% FA (solvent B):100% $H_2O$ supplemented with 0.2% FA (solvent A). Analysis was performed using a 15 mins gradient of 0.5-99.5% solvent B (0.5% 0-2 mins, 0.5-15.0% 2-5 mins, 15.0-99.5% 5-10 mins, 99.5% 10-13 mins, 99.5-0.5% 13-13.1 mins, 0.5% 13.1-15 mins) and a flow rate of 600 µL/min. Wash steps in-between sample analysis were performed using the same gradients.

Targeted metabolomics was performed by pre-selecting the desired precursor ion masses of metabolites of interest for fragmentation (where m/z mass 1 → mass 2 indicates m/z precursor ion → daughter ion). The mass spectrometer was operated in positive ion mode for the detection of β-NMN (m/z 335.0 → 123.0), 6Yn-Ad (m/z 306.3 → 174.3), 6Yn-Pro (m/z 623.1 → 432.5), 6Yn-AMP (m/z 386.3 → 174.3), 6Yn-$NAD^+$ (m/z 702.5 → 174.2) and $NAD^+$ (m/z 644.0 → 136.0). Negative ion mode was used for the detection of 6Yn-ATP (m/z 544.2 → 159.0) and ATP (m/z 506.2 → 408.2). To take into account metabolic degradation over time, the sample order was randomized. Pooled quality control (QC) samples were also injected throughout the run. Multiple reaction monitoring transitions were used for all analytes and internal standards. Retention time, exact mass and MS/MS spectra of quantified compounds were matched to an authentic standard. Data were processed using Analyst 1.6.2 (SCIEX) software. This involved peak detection, integration, calibration curve regression, and analyte quantification. Smoothing and peak-splitting factors were set as 3 and 2, respectively.

**Cloning of NMNAT1 Sequence and NMNAT1 Overexpression**

Human NMNAT1 DNA sequence was extracted by PCR from pMXs_FLAG-NMNAT1 vector (Addgene #133259) by designing and using the following primers: forward – 5'-ATCTACcatggaaaattccgagaagactgaag-3'; reverse – 5'-CGAATTCTCatgtcttagcttctgcagtg-3'. This was performed by following the 'Q5 High-Fidelity DNA Polymerase (M0491)' protocol (New England Biolabs) and using the reagents and amounts stated in the protocol, and using 1 ng of the plasmid. The following PCR cycle conditions were used: initial denaturation step – 98 ˚C for 30 s; 30 cycles of 98 ˚C for 10 s, (primer annealing at) 63 ˚C for 25 s, and 72 ˚C for 30 s; final elongation step – 72 ˚C for 2 mins. Extraction of NMNAT1 DNA sequence was confirmed by DNA gel electrophoresis by running the PCR reaction product on a TAE 1% agarose gel. The NMNAT1 fragment was then cloned into an in-house generated empty CMV vector. For preparation of the empty CMV vector, the following primers were designed and used: forward – 5'-agctaagacaTGAGAATTCGACTCTAGAGGATC-3'; reverse – 5'-cggaattttcCATGGTAGATCGATCTGAATTAATTC-3'. PCR cycle conditions were used: initial denaturation step – 98 ˚C for 30 s; 30 cycles of 98 ˚C for 10 s, (primer annealing at) 64 ˚C for 186 s, and 72 ˚C for 30 s; final elongation step – 72 ˚C for 2 mins. Preparation of the empty CMV vector was confirmed by DNA gel electrophoresis by running the PCR reaction product on a TAE 1% agarose gel.

Template DNA Digestion and PCR Product Purification

To ensure the DNA templates were removed from the PCR products prior to Gibson Assembly, DNA template digestion was performed with DpnI enzyme. This was performed by incubating 43 µL of each PCR product with 10X Cutsmart$^R$ Buffer (5 µL, New England Biolabs) and DpnI enzyme (2 µL, 40 units,

New England Biolabs) at 37 ˚C for 2 hours. After the incubation, DpnI was inactivated by heating at 80 ˚C for 20 mins. The samples were subsequently purified using the QIAquick PCR Purification Kit (QIAGEN).

Gibson Assembly

Each pair of purified PCR fragments, NMNAT1 DNA insert + empty CMV plasmid, was assembled into a single plasmid vector via Gibson Assembly. This was performed by following the 'Gibson Assembly (E5510)' protocol (New England Biolabs), and using the reagents and amounts stated in the protocol. Overall, 27 ng of the NMNAT1 DNA fragment were ligated to 100 ng of vector backbone by incubating the two fragments at 50 ˚C for 1 hour with the Gibson Assembly Master Mix (2X) (New England Biolabs). Successful generation of the ligated product was subsequently confirmed by Genewiz sequencing.

Transfection of HEK293T Cells for NMNAT1 Overexpression

Transfection of HEK293T cells was performed using Lipofectamine 2000, in accordance with the manufacturer's protocol, and 1000 ng of plasmid DNA were used per 1 mL of seeding/transfection media. For 6-well plates and 6 cm dishes, 2 mL and 5 mL of seeding/transfection media were used per well and per dish, respectively. Transfection was performed for 24 h and the media was replaced with fresh culture media (1 mL per well for 6-well plates and 3 mL per 6 cm dish), prior to treatment with the relevant probes. Overexpression of NMNAT1 was confirmed by western blotting using an α-NMNAT1 antibody (sc-271557, dilution 1:500 (v/v), Santa Cruz Biotechnology).

#### Supplementary Figures and Tables

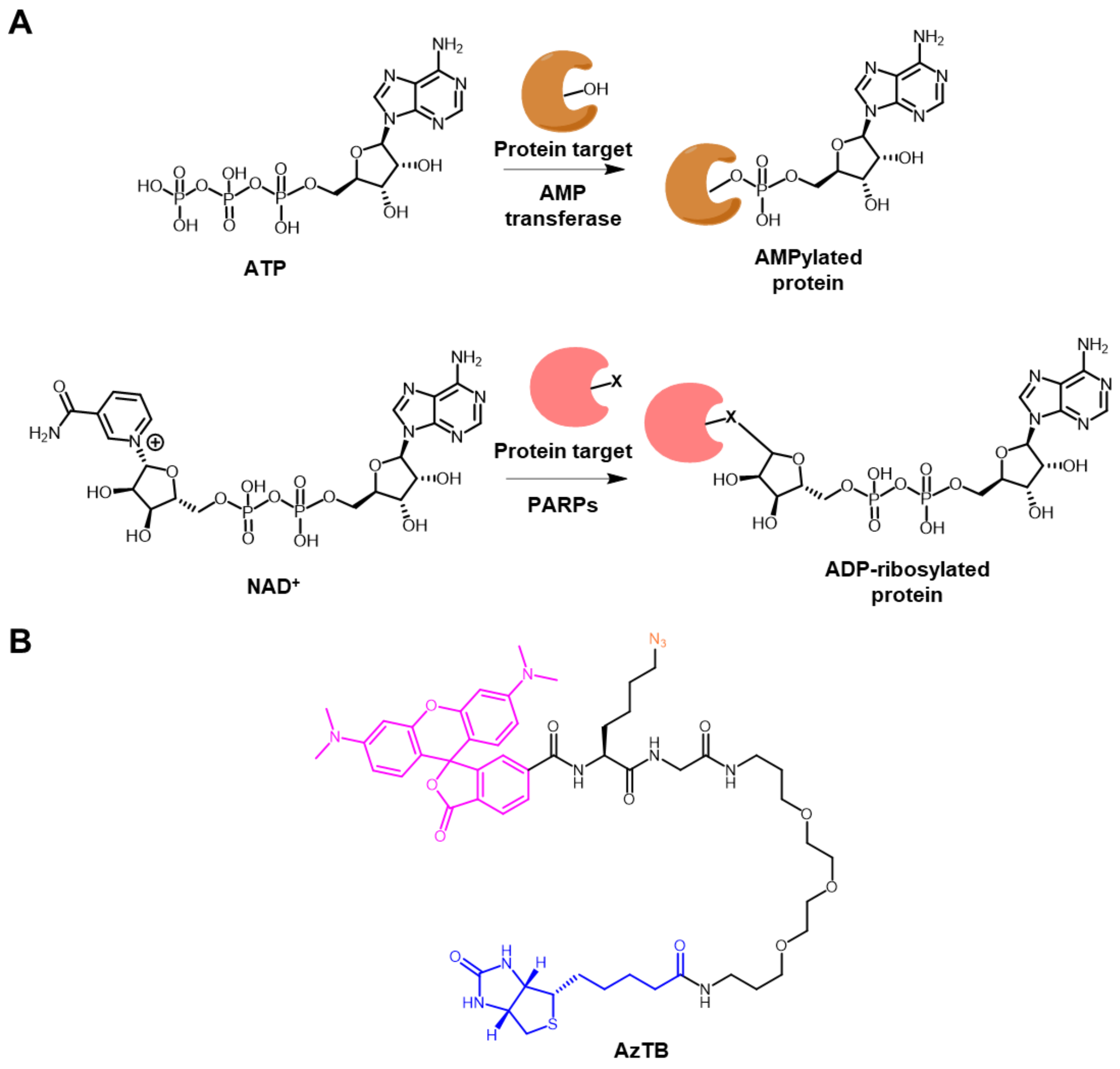

**Figure S1. Overview of protein AMPylation and protein ADP-ribosylation, and chemical structure of the capture reagent azide-TAMRA-biotin (AzTB) used in click chemistry CuAAC ligation reactions.** (**A**) ATP is used by AMP transferases as a cofactor in protein AMPylation on S, Y, and T residues. $NAD^+$ is used by PARPs as a cofactor in protein ADP-ribosylation on C, H, K, S, Y, T, R, E and D residues. (**B**) Chemical structure of azide-TAMRA-biotin. TAMRA fluorophore (5-carboxytetramethyl-rhodamine) is shown in pink, biotin moiety is shown in blue and the azide click chemistry group is shown in orange.

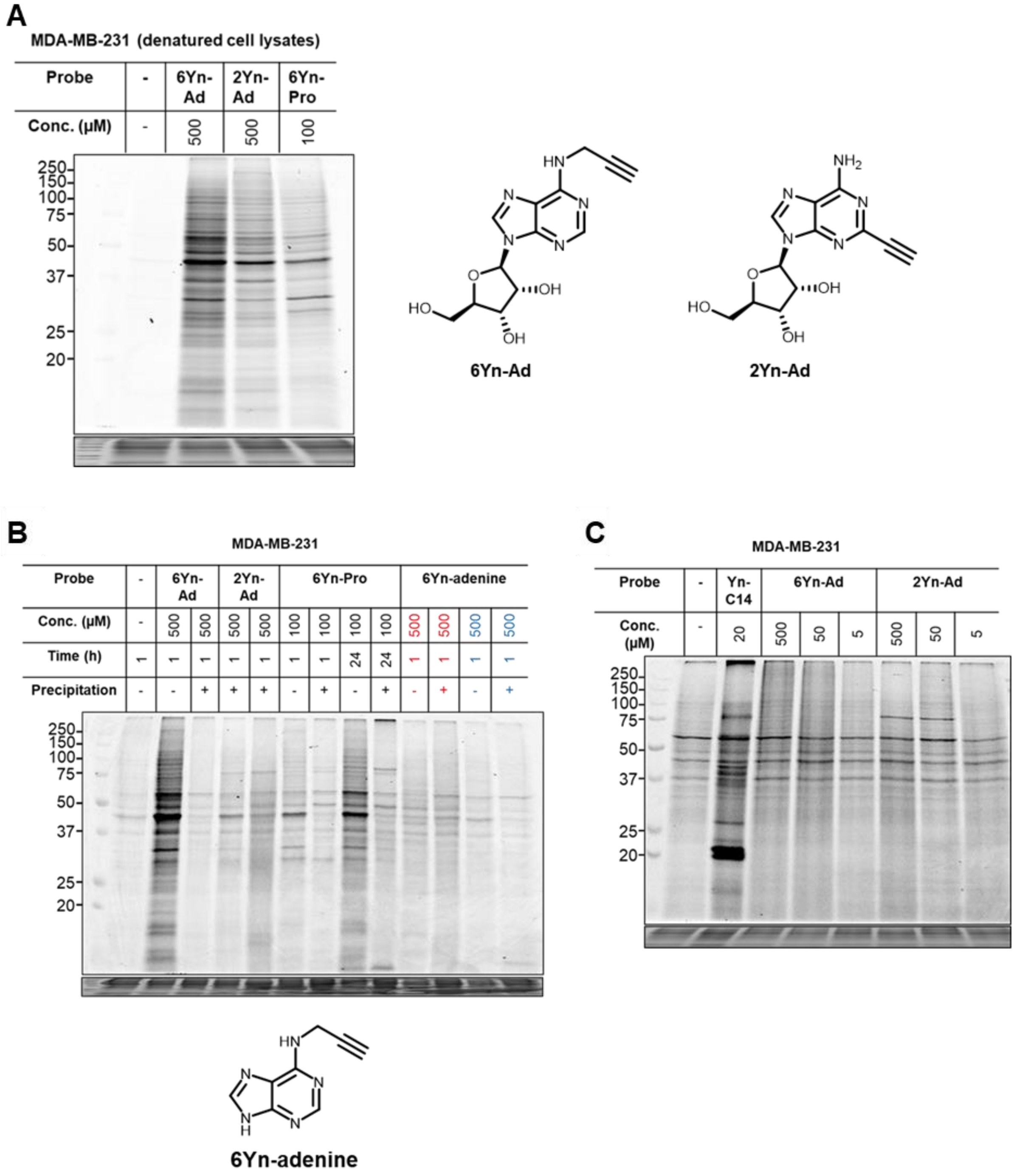

**Figure S2. Protein labelling observed in probe-treated live MDA-MB-231 cell samples.** (**A**) Protein labelling in denatured, enzymatically inactive MDA-MB-231 cell lysates by IGF (top panel). Protein loading was visualized by Ponceau S stain (bottom panel). Lysates were treated with 6Yn-Ad (500 µM, 1 h), 2Yn-Ad (500 µM, 1 h) or 6Yn-Pro (100 µM, 1 h) at r.t. (750 rpm) and ligation of labelled proteins to AzTB was performed by CuAAC click chemistry conjugation. Coomassie staining (bottom panel) = loading control**.** (**B**) Protein labelling in MDA-MB-231 cells by IGF (top panel) and protein loading by Coomassie staining (bottom panel). Cell treatment was performed with DMSO (negative control), 6Yn-Ad (500 µM, 1 h), 2Yn-Ad (500 µM, 1 h), 6Yn-Pro (100 µM, 1 h and 24 h) and 6Yn-adenine. For 6Yn-adenine-treated samples, for the lanes highlighted in red treatment was performed in live cells and for the lanes highlighted in blue treatment was performed in denatured, enzymatically inactive lysates. Live cell samples were lysed with 4% (w/v) SDS lysis buffer. Samples for which protein precipitation has been performed after cell lysis and prior to CuAAC click chemistry conjugation is indicated by +. (**C**) Protein labelling in

MDA-MB-231 cells by IGF (top panel) and protein loading by Coomassie staining (bottom panel). Cell treatment was performed with DMSO (negative control), Yn-C14 (myristic acid alkyne (CuAAC positive control), 20 µM, 24 h), 6Yn-Ad (500, 50 or 5 µM, 1 h) or 2Yn-Ad (500, 50 or 5 µM, 1 h). Treated MDA-MB-231 were lysed with 4% (w/v) SDS lysis buffer, protein precipitation was performed for all samples and probe-tagged proteins were ligated to azide-TAMRA by CuAAC.

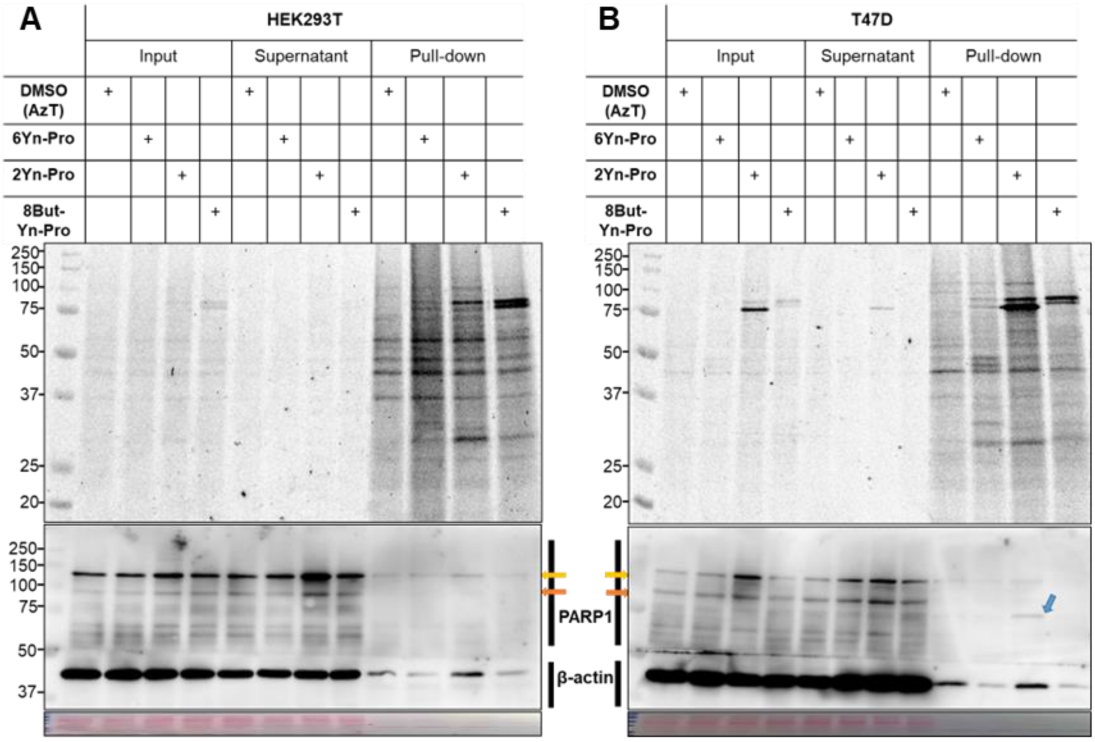

**Figure S3. Protein labelling observed in probe-treated live HEK293T and T47D cell samples.** (**A**) Protein labelling in HEK293T cells by IGF (top panel) and western blot (middle panel). Protein loading was visualized by Ponceau S stain (bottom panel). Cell treatment was performed with DMSO (0.1% v/v), 6Yn-Pro (100 µM, 24 h), 2Yn-Pro (100 µM, 24 h) or 8But-Yn-Pro (100 µM, 24 h) followed by cell lysis (4% (w/v) SDS lysis buffer), protein precipitation and ligation of tagged proteins to AzTB by CuAAC. Input shows protein labelling before enrichment, pull-down shows protein labelling after enrichment, supernatant shows depletion of protein labelling from input samples after the enrichment step. Western blotting was performed using PARP1 antibody (Santa Cruz Biotechnology, sc-8007). Yellow and orange arrows indicate full-length (~113 kDa) and cleaved (~89 kDa) PARP1 fragment, respectively; blue arrow indicates a potential alternative cleaved PARP1 fragment. β-actin and Ponceau S staining (bottom panel) = loading controls. (**B**) Protein labelling in T47D cells by IGF (top panel) and western blot (middle panel). Cell treatment and western blotting were performed as described in (**A**).

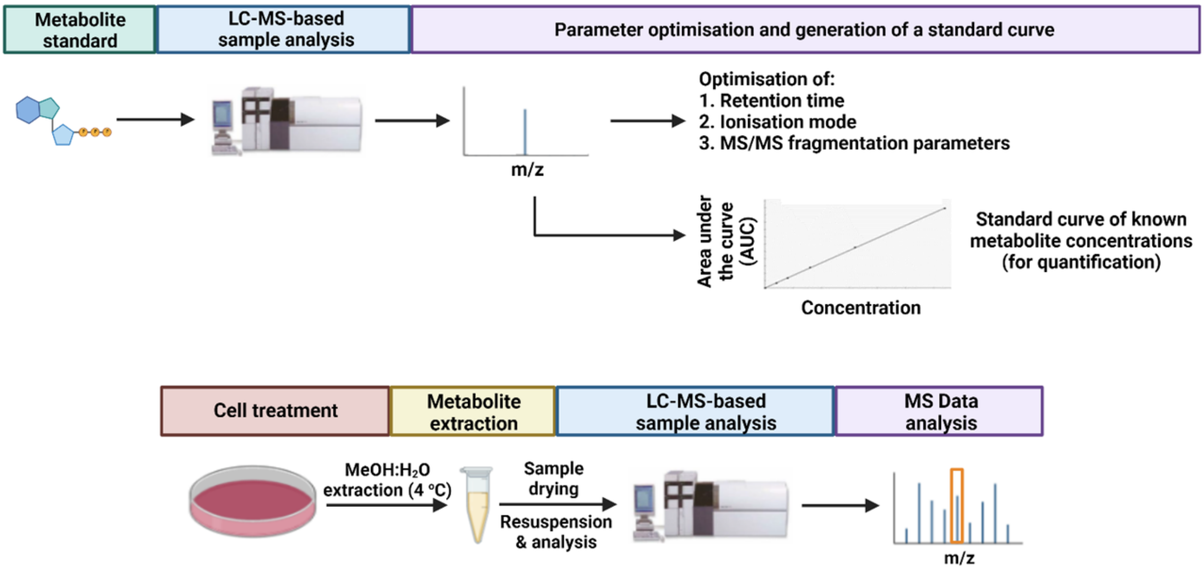

**Figure S4. Overview of the workflow followed for sample preparation for analysis by targeted metabolomics.** Prior to generation of cell samples, LC-MS method development and metabolite parameter optimizations were performed for each individual metabolite of interest using pure stock solutions of known concentrations of the metabolites (6Yn-AMP, 6Yn-ATP, 6Yn-NAD$^+$, ATP, NAD$^+$, β-NMN). A standard curve was then generated and used as a reference for quantification of metabolite levels from biological samples. For preparation of cell samples for analysis by targeted metabolomics, cells were treated with 6Yn-Pro (500 µM, 24 h), and where relevant, with NRH (500 µM, 24 h) and/or in the presence of NMNAT1 overexpression (24 h). Metabolite extraction was performed in ice-cold MeOH:$H_2O$ 80:20 (v/v) mixture on ice and samples were subsequently dried. Following samples processing and analysis by LC-MS, metabolites were identified based on m/z ratio, retention time and fragmentation patterns.

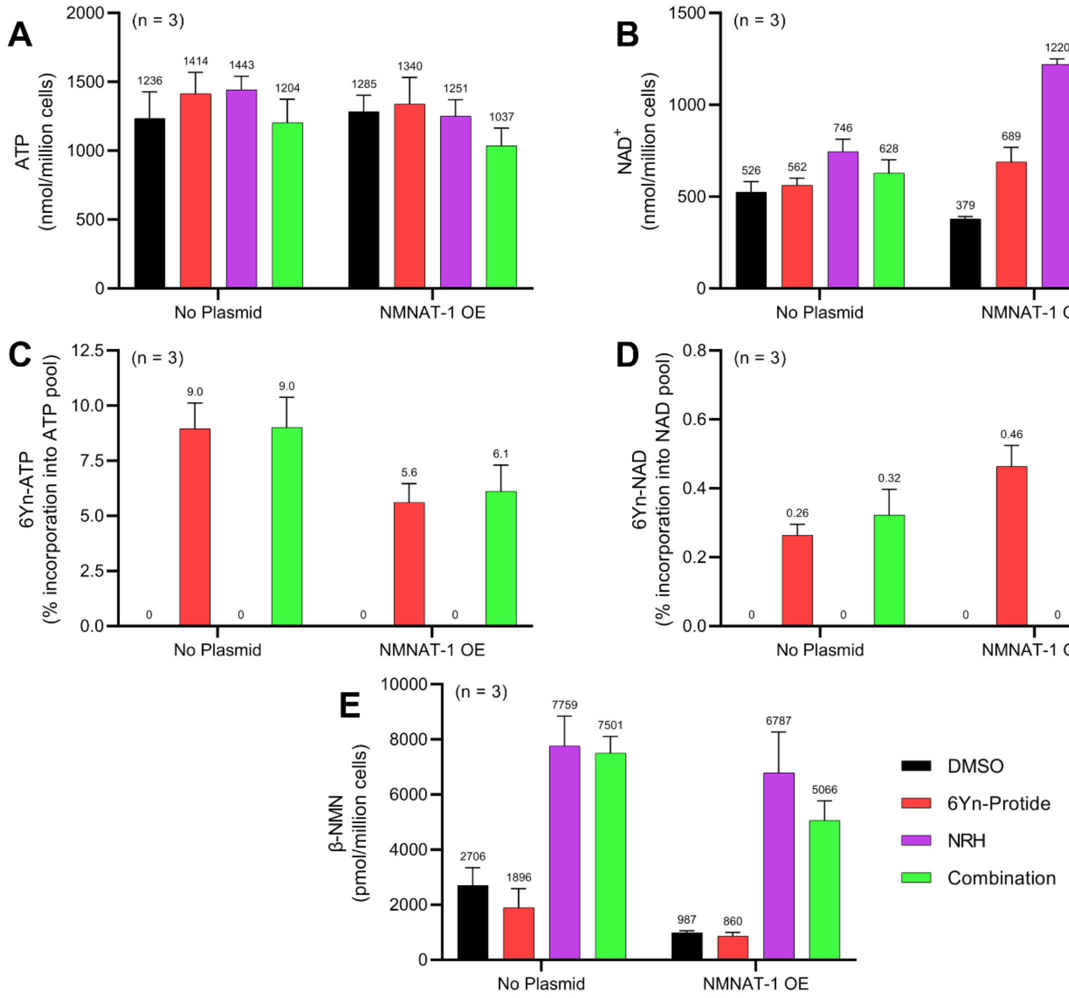

**Figure S5. Metabolite profiling in HEK293T cells reveals increased 6Yn-ATP consumption and increased intracellular 6Yn-NAD$^+$ production following overexpression of NMNAT1.** Metabolite profiling in HEK293T cells following treatment with probe 6Yn-Pro (100 µM, 24 h), with/without NRH co-treatment (500 µM, 24 h), with/without NMNAT1 overexpression (24 h), or DMSO (negative control). This was performed for the same samples shown in **Figure 3A-C**. Metabolite levels for β-NMN, ATP and NAD$^+$ were measured and quantified as pmol/million cells (for β-NMN), or nmol/million cells (for ATP and NAD$^+$). % incorporation of 6Yn-ATP and 6Yn-NAD$^+$ were calculated for the data shown in **Figure 3B** and **3C**, respectively. (**A**) Measured levels of ATP. (**B**) Measured levels of NAD$^+$. (**C**) % incorporation of 6Yn-ATP into the overall ATP pool. (**D**) % incorporation of 6Yn-NAD$^+$ into the overall NAD$^+$ pool. (**E**) Measured levels of β-NMN.

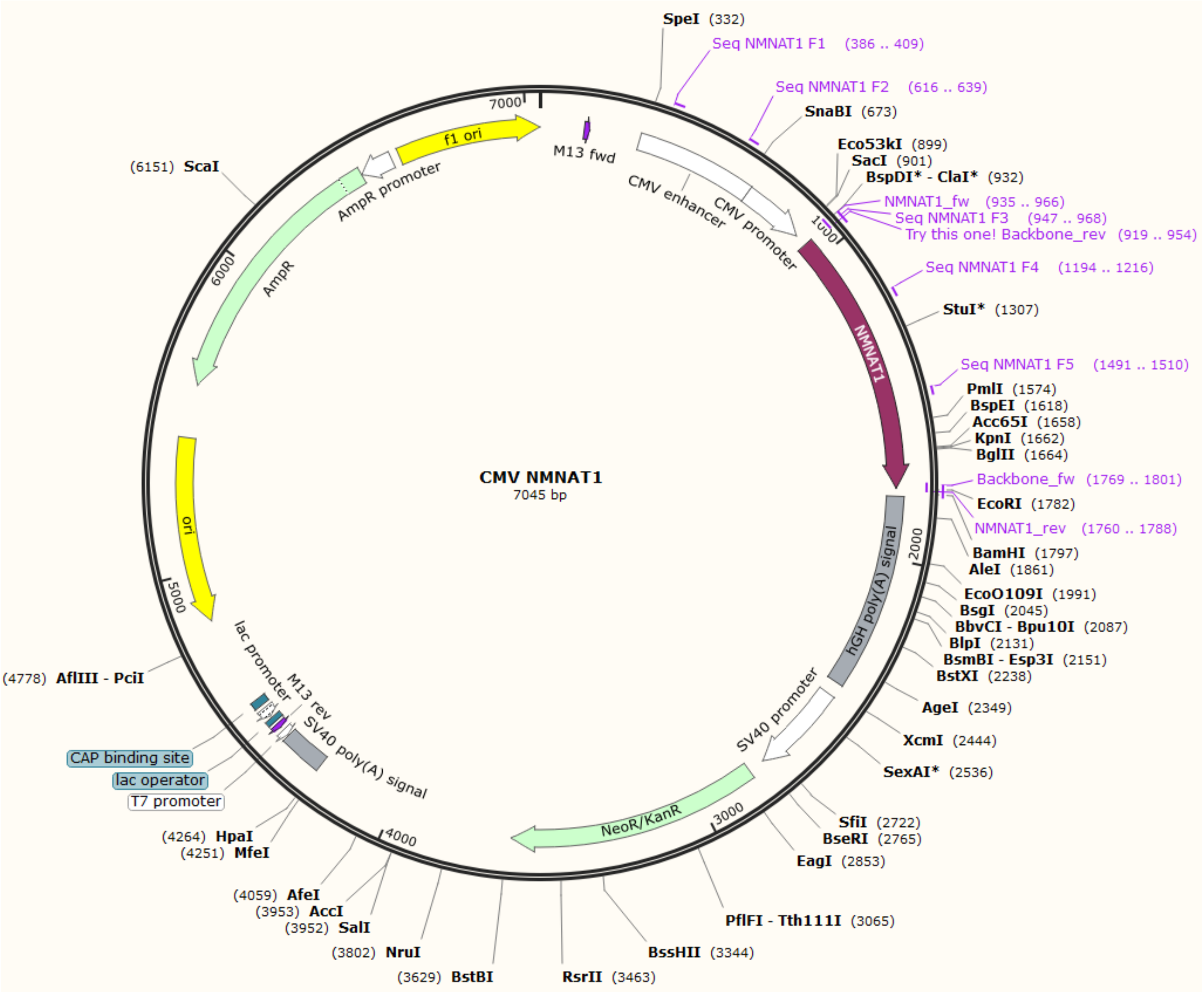

**Figure S6. Plasmid map for CMV NMNAT1 plasmid.**

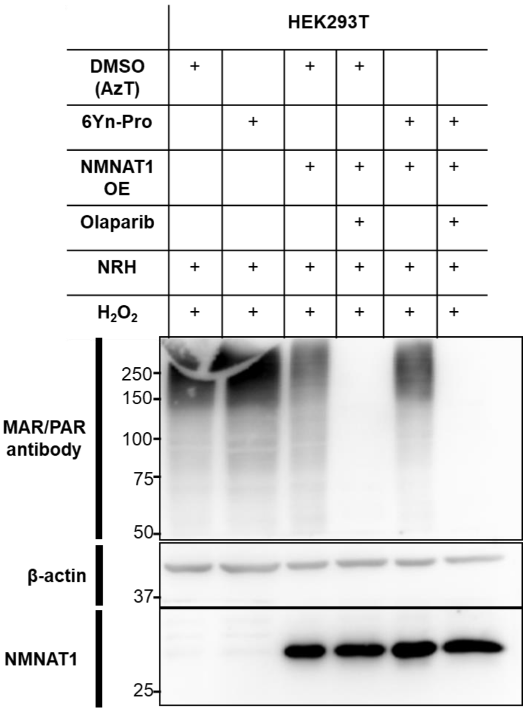

**Figure S7. Overexpression of NMNAT1 leads to a reduction in overall cellular protein ADP-ribosylation levels.** Western blot images showing overall ADP-ribosylation (MAR/PAR) protein levels in HEK293T cells at the indicated experimental conditions. For the relevant samples, NMNAT1 was over-expressed for 24 h prior to cell treatment with DMSO or 6Yn-Pro (100 µM, 24 h), with/without olaparib pre-treatment (10 µM, 2 h). All cells were co-treated with NRH (500 µM, 24 h) followed by $H_2O_2$ (in PBS, 2 mM, 10 min) prior to cell lysis. Blotting for MAR/PAR protein levels was performed with α-Poly/Mono-ADP ribose (anti-MAR/PAR) antibody (Cell Signaling Technology, #83732). Blotting for NMNAT1 expression levels was performed with α-NMNAT1 antibody (Santa Cruz Biotechnology, sc-271557). β-actin antibody = loading control.

**Table S1.** Summary of the extent of PARP1 enrichment, shown as $Log_2$ Fold change and Absolute Fold increase**,** in 6Yn-Pro treated samples compared to the corresponding DMSO control samples for the data shown in **Figure 4A-D**.

| Condition | $Log_2$ Fold change (PARP1) | Absolute Fold increase (PARP1) |
|---|---|---|
| 6Yn-Pro | 0.883 | 1.84 |
| 6Yn-Pro + NRH | 1.15 | 2.22 |
| 6Yn-Pro + $H_2O_2$ | 1.30 | 2.46 |
| 6Yn-Pro + NRH + $H_2O_2$ | 2.46 | 5.51 |

**Table S2.** Summary of the total number of proteins enriched ($Log_2$ Fold change > 0) in 6Yn-Pro treated samples compared to the corresponding DMSO control samples for the data shown in **Figure 4A-D**. The number of proteins identified as ADP-ribosylated proteins in previous studies is shown, alongside the % fraction they represent in each sample.

| | 6Yn-Pro | 6Yn-Pro + NRH | 6Yn-Pro + $H_2O_2$ | 6Yn-Pro + NRH + $H_2O_2$ |
|---|---|---|---|---|
| **Total number of proteins enriched** | 104 | 114 | 155 | 189 |
| **No. of proteins identified in at least one ADPr study** | 41 (39%) | 51 (45%) | 79 (51%) | 97 (51%) |
| **No. of proteins identified in two or more ADPr stud-ies** | 31 (30%) | 38 (33%) | 56 (36%) | 71 (38%) |

**Table S3.** Significantly enriched proteins in volcano plot **Fig. 4E**. Enrichment ($Log_2$ Fold change) and significance ($-Log_{10}$(p-value)) values for each protein are shown.

| **Protein** | **$Log_2$ Fold change** | **$-Log_{10}$(p-value)** |
|---|---|---|
| ACP2 | 4.20174 | 2.335943 |
| CTSA | 3.247465 | 3.926593 |
| PPME1 | 3.198154 | 2.952403 |
| NCEH1 | 3.142584 | 1.302133 |
| SLC25A3 | 3.119327 | 4.763065 |
| PLD3 | 2.62413 | 4.801288 |
| PFKP | 2.582147 | 2.784853 |
| MIF | 2.547378 | 1.960166 |
| HEXB | 2.465707 | 3.508306 |
| TPP1 | 2.208282 | 1.404106 |
| CTSB | 2.175413 | 1.295072 |
| RPLP1 | 2.037982 | 2.155392 |
| PARP1 | 1.603592 | 1.500945 |
| RBMX | 1.461025 | 2.434493 |
| PREP | 1.161517 | 2.30356 |
| CTSD | 1.15094 | 2.839743 |
| GSTO1 | 1.044719 | 1.929222 |
| TALDO1 | 0.909082 | 1.401506 |
| RALGAPA1 | 0.902733 | 2.673543 |
| PDIA6 | 0.811351 | 2.131374 |
| PDIA3 | 0.675349 | 1.498881 |
| CCT6A | 0.517229 | 3.571692 |
| HMGB1 | 0.504435 | 1.680521 |

**Table S4.** Significantly enriched proteins in volcano plot **Fig. 4F**. Enrichment ($Log_2$ Fold change) and significance ($-Log_{10}$(p-value)) values for each protein are shown.

| **Protein** | **$Log_2$ Fold change** | **$-Log_{10}$(p-value)** |
|---|---|---|
| PARP1 | 0.631482 | 1.538862 |

**Table S5.** Significantly enriched proteins in volcano plot **Fig. 4G**. Enrichment ($Log_2$ Fold change) and significance ($-Log_{10}$(p-value)) values for each protein are shown.

| **Protein** | **$Log_2$ Fold change** | **$-Log_{10}$(p-value)** |
|---|---|---|
| ACP2 | 4.179209 | 2.683746 |
| CTSA | 3.386311 | 2.744564 |
| SLC25A3 | 3.028913 | 4.652359 |
| PLD3 | 2.9319 | 2.874991 |
| PPME1 | 2.872788 | 2.416644 |
| PFKP | 2.465608 | 3.39613 |
| HEXB | 2.233832 | 4.14428 |
| PREP | 1.432713 | 2.336235 |
| HSPA5 | 1.309479 | 4.25234 |
| PDIA6 | 0.905974 | 2.81107 |
| PDIA3 | 0.696087 | 1.377676 |
| CYB5R1 | 0.61815 | 1.936696 |
| HMGB1 | 0.613416 | 1.832451 |
| HINT1 | 0.585216 | 1.428414 |

**Table S6.** Significantly enriched proteins in volcano plot Fig. 4H. Enrichment ($Log_2$ Fold change) and significance ($-Log_{10}$(p-value)) values for each protein are shown.

| Protein | $Log_2$ Fold change | $-Log_{10}$(p-value) |
|---|---|---|
| CPVL | 5.334544 | 1.772197 |
| PARP1 | 3.846672 | 1.652674 |
| ACP2 | 2.13291 | 2.373061 |
| PREP | 2.112498 | 1.95144 |
| MYL6 | 1.850293 | 1.270927 |
| MIF | 1.713501 | 1.233575 |
| CTSB | 1.681155 | 1.400796 |
| CSE1L | 1.289517 | 1.329131 |
| NMNAT1 | 1.03635 | 2.731387 |
| APMAP | 1.003174 | 1.265947 |
| ACTR2 | 0.820199 | 1.58915 |
| PDIA3 | 0.767563 | 1.325805 |

**Table S7.** Enrichment ($Log_2$ Fold change) and significance ($-Log_{10}$(p-value)) values of literature-known ADP-ribosylated proteins with $Log_2$ fold change > 0.5 in **Figure 5A** (after $H_2O_2$ treatment, proteins highlighted in green in **Figure 5A**) that match in the non-$H_2O_2$ treated samples in **Figure 5B** (in the absence of $H_2O_2$ treatment).

| Protein | $Log_2$ Fold change | $-Log_{10}$(p-value) |
|---|---|---|
| PARP1 | 3.846672 | 1.652674 |
| HIST1H1E | 2.815618 | 0 |
| MIF | 1.713501 | 1.233575 |
| HNRNPD | 1.709938 | 0.748445 |
| PPIB | 1.660015 | 0 |
| PA2G4 | 1.297882 | 0 |
| PGAM1 | 1.124841 | 0.576642 |
| EIF4A1 | 1.087206 | 0.956283 |
| HMGB1 | 1.048254 | 0 |
| ANXA2 | 1.019864 | 0.571956 |
| DDX5 | 0.985273 | 0 |
| RPL29 | 0.955983 | 0 |
| PDIA3 | 0.767563 | 1.325805 |
| RPL22 | 0.755857 | 0.672817 |
| HIST1H1C | 0.716428 | 0.998411 |
| LRRC59 | 0.713186 | 0.407246 |
| HNRNPA1 | 0.667481 | 0 |
| HSPA9 | 0.625729 | 0.58102 |
| MAPRE1 | 0.617895 | 0.29806 |
| SYNCRIP | 0.524401 | 0 |
| HNRNPH1 | 0.51946 | 0.650085 |

**Table S8.** Enrichment ($Log_2$ Fold change) and significance ($-Log_{10}$(p-value)) values of literature-known ADP-ribosylated proteins with $Log_2$ fold change > 0.5 in **Figure 5A** (after $H_2O_2$ treatment, proteins highlighted in red in **Figure 5A**) that do not match in the non-$H_2O_2$ treated samples in **Figure 5B** (in the absence of $H_2O_2$ treatment).

| Protein | Log2 Fold change | -Log10(p-value) |
|---|---|---|
| SMAP | 3.336305 | 0 |
| PRPF40A | 2.948371 | 0 |
| CDKN2A | 2.294599 | 0 |
| PAK2;PAK1;PAK3 | 2.12165 | 0.42355 |
| RRS1 | 1.889938 | 0 |
| MYL6 | 1.850293 | 1.270927 |
| CSE1L | 1.289517 | 1.329131 |
| SMARCA5 | 1.253673 | 0 |
| APEX1 | 1.129928 | 0 |
| RCC2 | 0.722015 | 0.385906 |
| SUPT5H | 0.700761 | 0.588136 |
| ILF2 | 0.598951 | 0 |
| TOMM20 | 0.56606 | 0.269199 |
| SRSF7 | 0.541213 | 0.580246 |

**Table S9.** Enrichment ($Log_2$ Fold change) and significance ($-Log_{10}$(p-value)) values of the matched (to **Figure 5A, Table S3**) literature-known ADP-ribosylated proteins in **Figure 5B** (in the absence of $H_2O_2$ treatment, highlighted in green in **Figure 5B**).

| Protein | Log2 Fold change | -Log10(p-value) |
|---|---|---|
| MIF | 0.932961 | 0.501271 |
| PPIB | 0.510748 | 0.880577 |
| HIST1H1E;HIST1H1D | 0.415435 | 0.740221 |
| HIST1H1C | 0.378419 | 0.212134 |
| HNRNPD | 0.211852 | 0.573071 |
| RPL22 | 0.073099 | 1.67071 |
| RPL29 | 0.065112 | 0.068278 |
| HNRNPH1 | 0.053975 | 0.376998 |
| LRRC59 | -0.00626 | 0.006765 |
| PARP1 | -0.04657 | 0.311125 |
| SYNCRIP | -0.01943 | 0.080263 |
| HSPA9 | -0.02136 | 0.084425 |
| MAPRE1 | -0.05238 | 0.060811 |
| PDIA3 | -0.07126 | 0.15963 |
| PGAM1;PGAM2;PGAM4 | -0.11119 | 1.644959 |
| HNRNPA1;HNRNPA1L2 | -0.1526 | 0.251962 |
| PA2G4 | -0.32071 | 0.267906 |
| DDX5 | -0.37503 | 0.402074 |
| EIF4A1;EIF4A2 | -0.58233 | 0.398695 |
| ANXA2 | -0.63286 | 0.460383 |
| HMGB1;HMGB1P1 | -0.76225 | 0.516123 |

**Table S10.** Enrichment ($Log_2$ Fold change) and significance ($-Log_{10}$(p-value)) values of the significantly enriched non-matching (to **Figure 5A, Table S4**) literature-known ADP-ribosylated proteins in **Figure 5B** (in the absence of $H_2O_2$ treatment, highlighted in red in **Figure 5B**).

| Protein | Log2 Fold change | -Log10(p-value) |
|---|---|---|
| ARHGDIA | 1.17246 | 0.546882 |
| VIM | 0.622622 | 2.706018 |
| SERBP1 | 0.547956 | 0.628423 |
| NME2;NME1;NME2P1 | 0.528581 | 0.4484 |
| STIP1 | 0.504067 | 0.358553 |

### NMR Spectra

#### $^1$H NMR of N-(prop-2-yn-1-yl)-9H-purin-6-amine (6Yn-adenine)

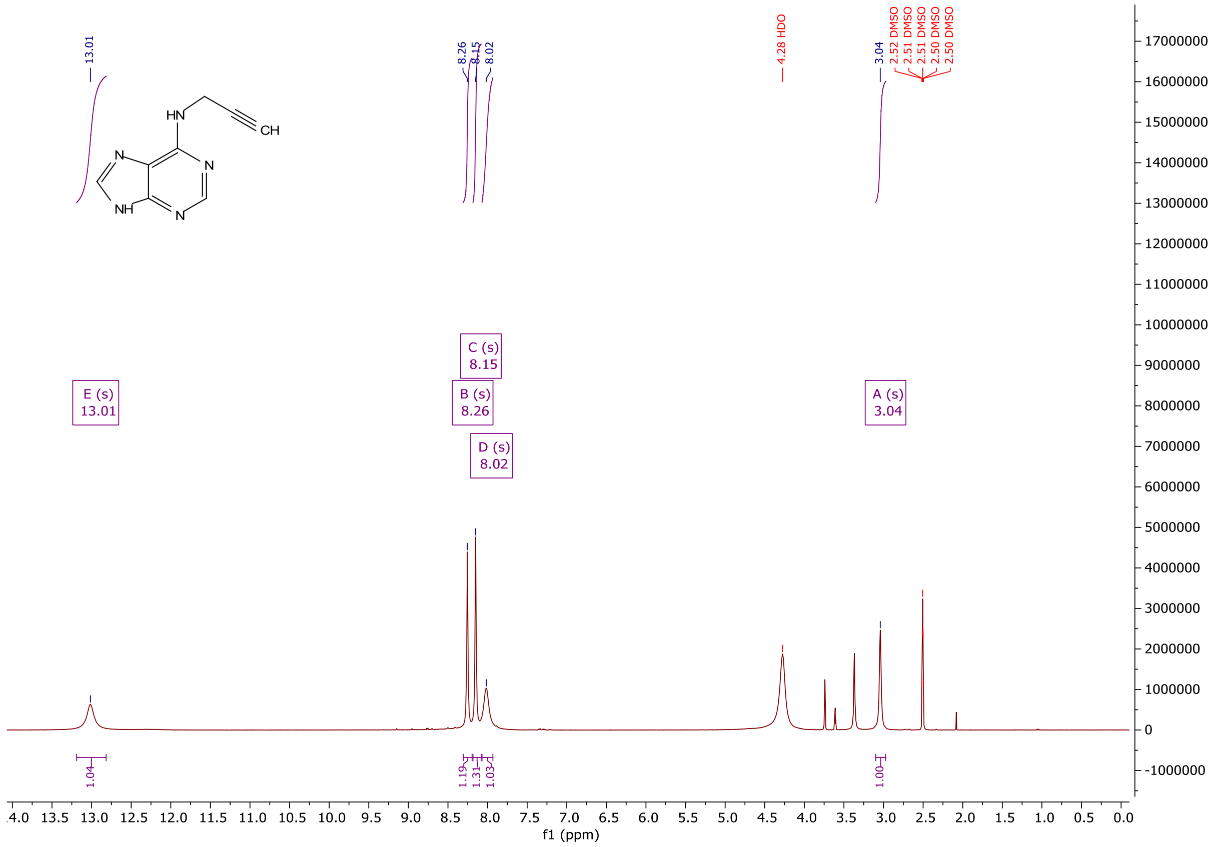

**$^{13}$C NMR of N-(prop-2-yn-1-yl)-9H-purin-6-amine (6Yn-adenine)**

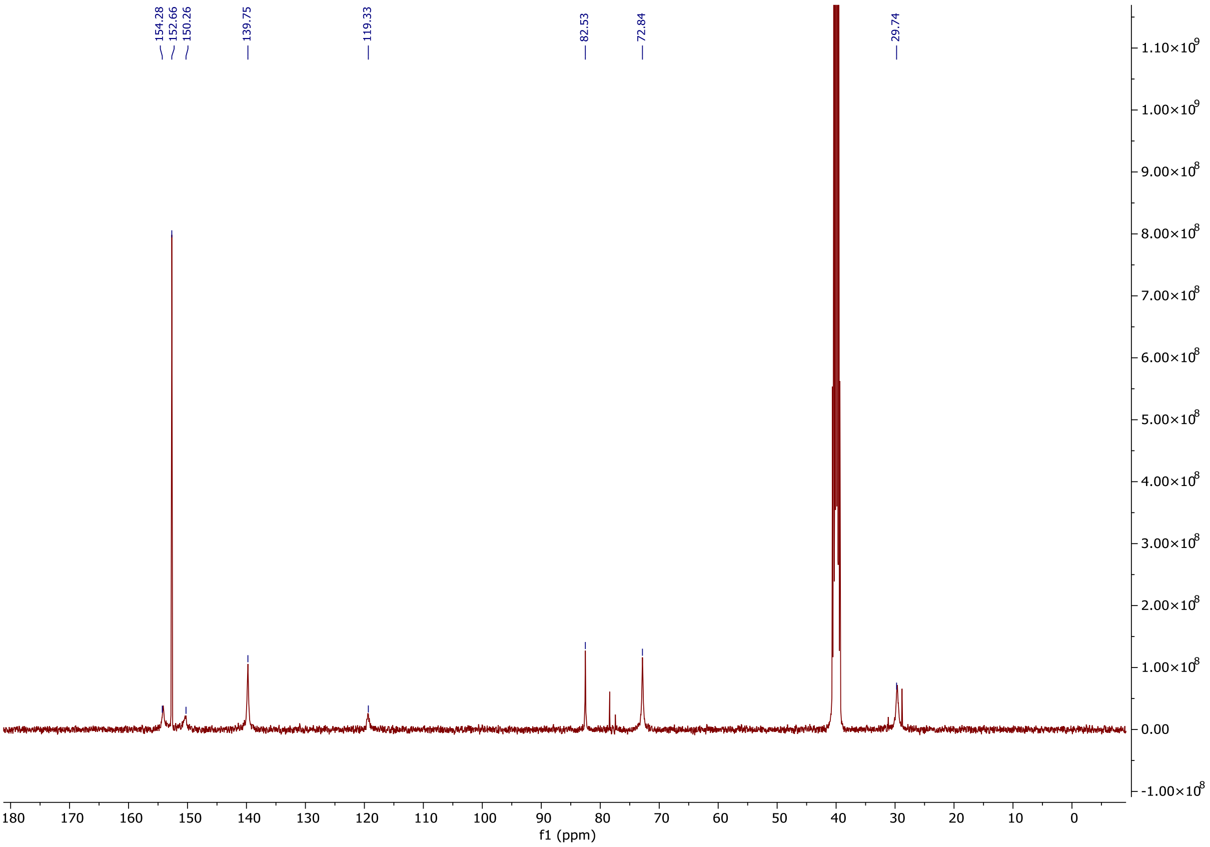

**$^{1}$H NMR of (2R,3S,4R,5R)-2-(hydroxymethyl)-5-(6-(prop-2-yn-1-ylamino)-9H-purin-9-yl)tetrahydrofuran-3,4-diol (6Yn-Ad)**

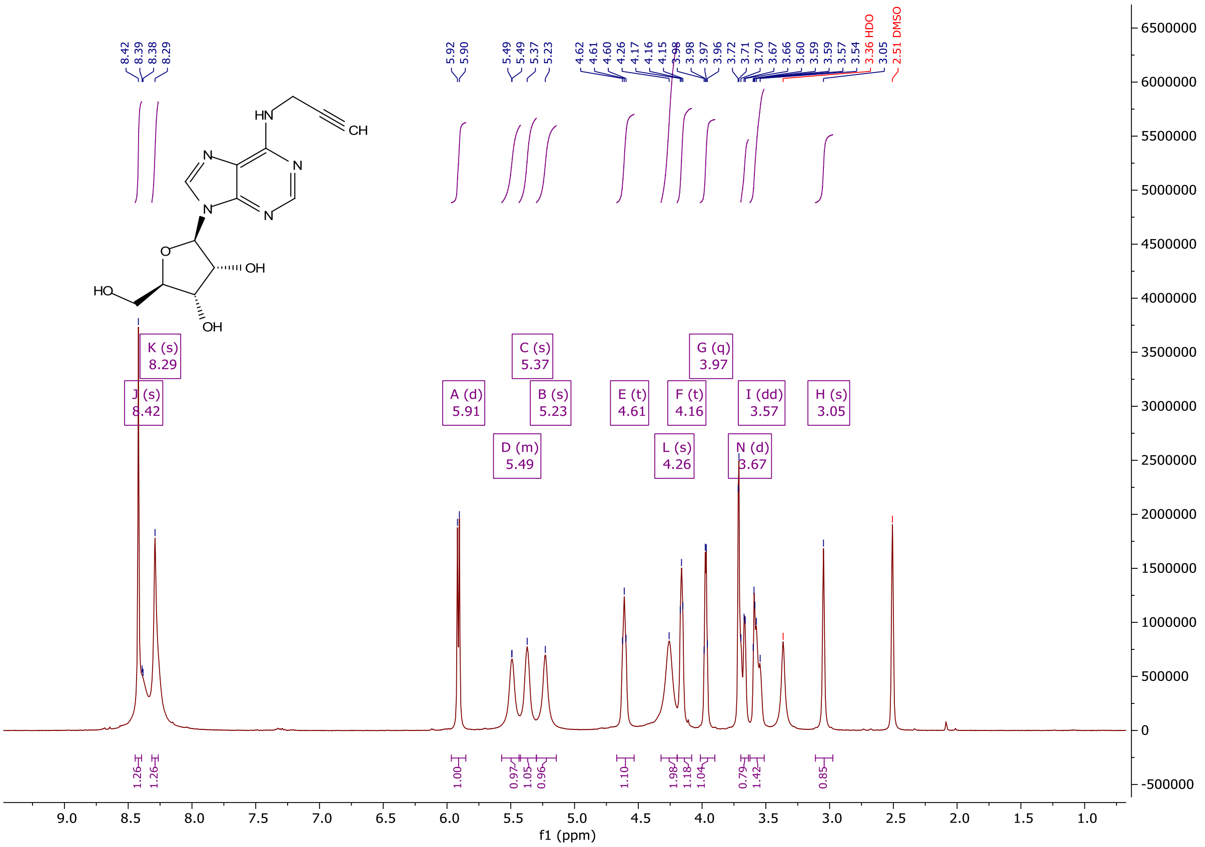

**[13]C NMR of (2R,3S,4R,5R)-2-(hydroxymethyl)-5-(6-(prop-2-yn-1-ylamino)-9H-purin-9-yl)tetrahydrofuran-3,4-diol (6Yn-Ad)**

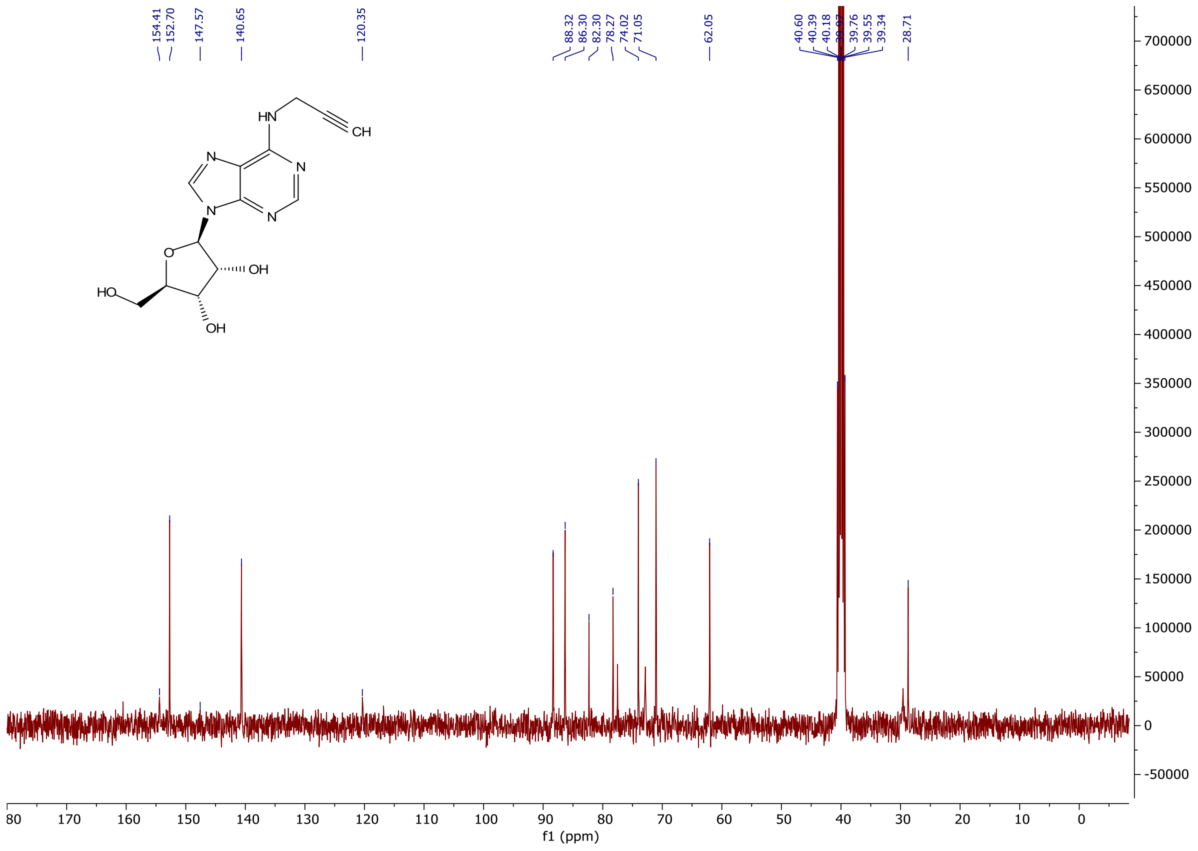

[1]H NMR of (2R,3R,4S,5R)-2-(6-amino-2-ethynyl-9H-purin-9-yl)-5-(hydroxymethyl)tetrahydrofuran-3,4-diol (2Yn-Ad)

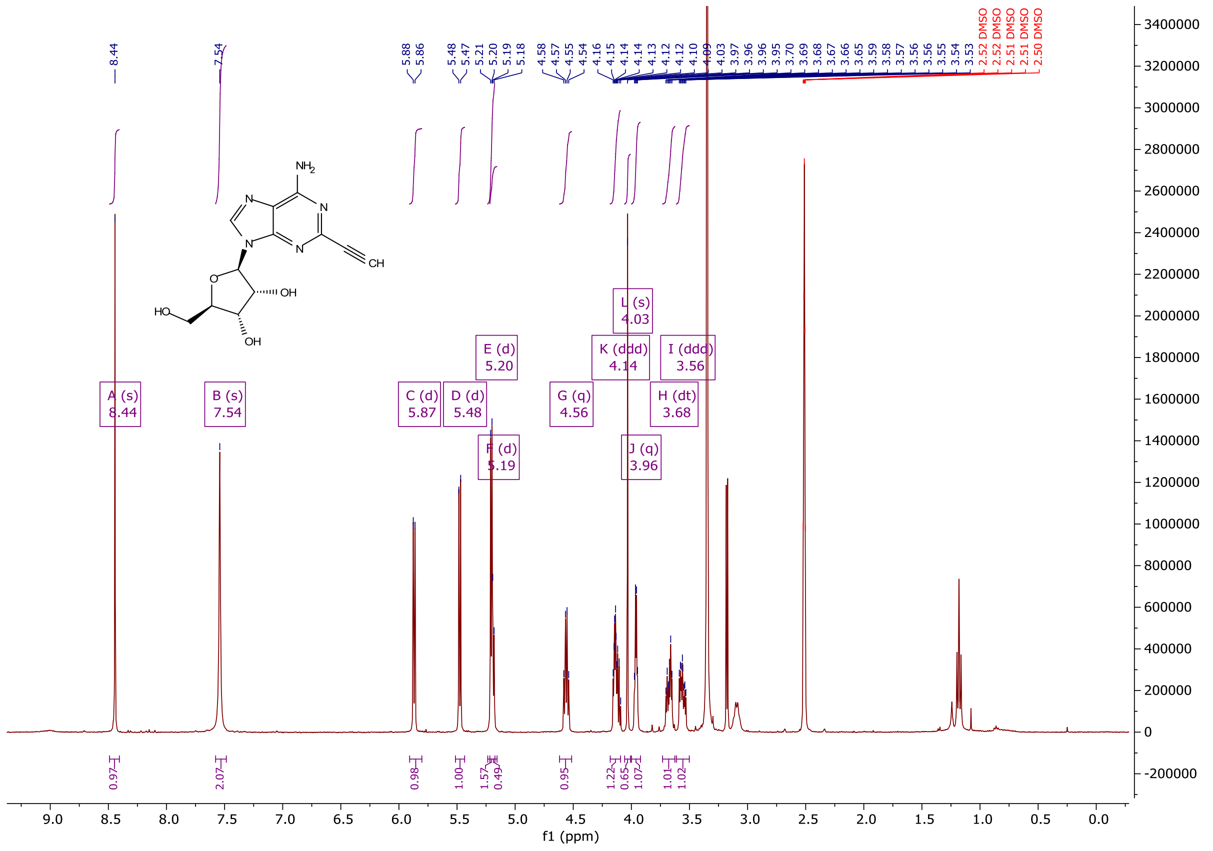

**[13]C NMR of (2R,3R,4S,5R)-2-(6-amino-2-ethynyl-9H-purin-9-yl)-5-(hydroxymethyl)tetrahydrofuran-3,4-diol (2Yn-Ad)**

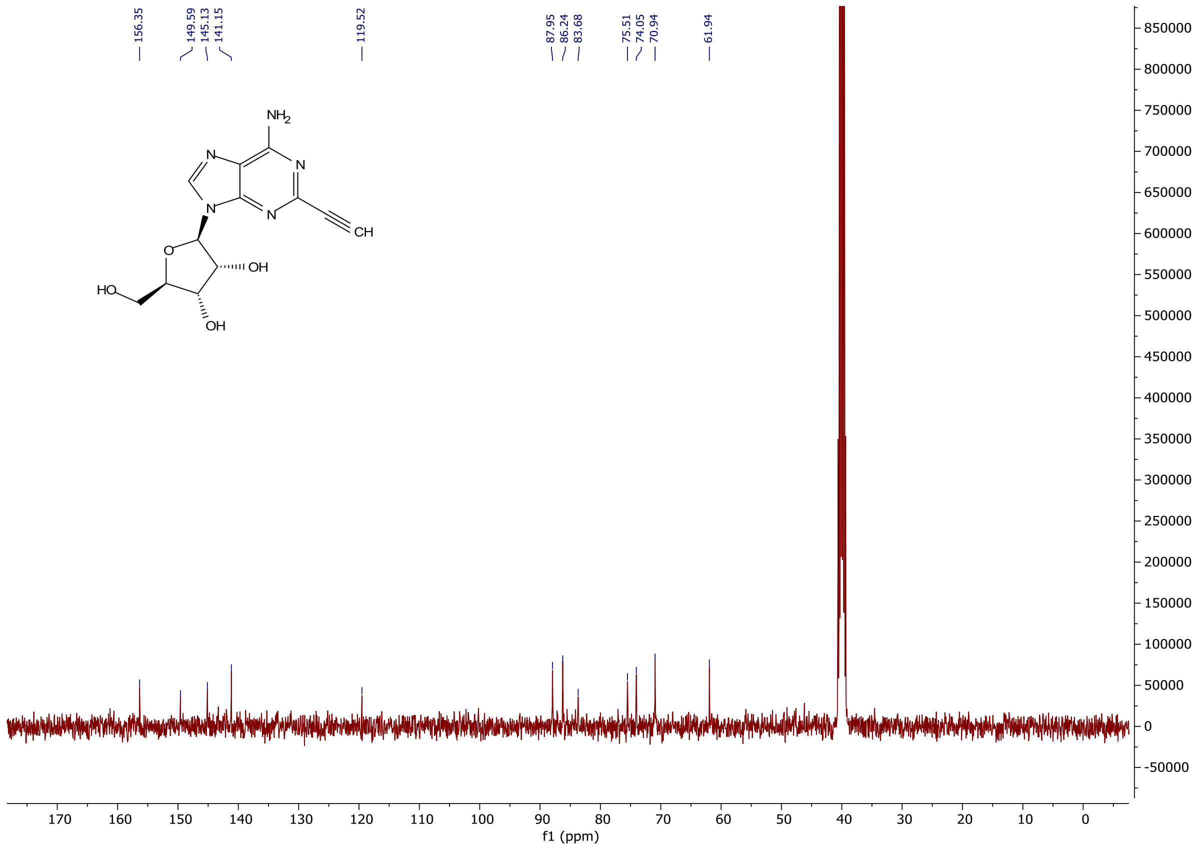

**[1]H NMR of (2R,3R,4S,5R)-2-(6-amino-8-bromo-9H-purin-9-yl)-5-(hydroxymethyl)tetra hydrofuran-3,4-diol (8Br-Ad)**

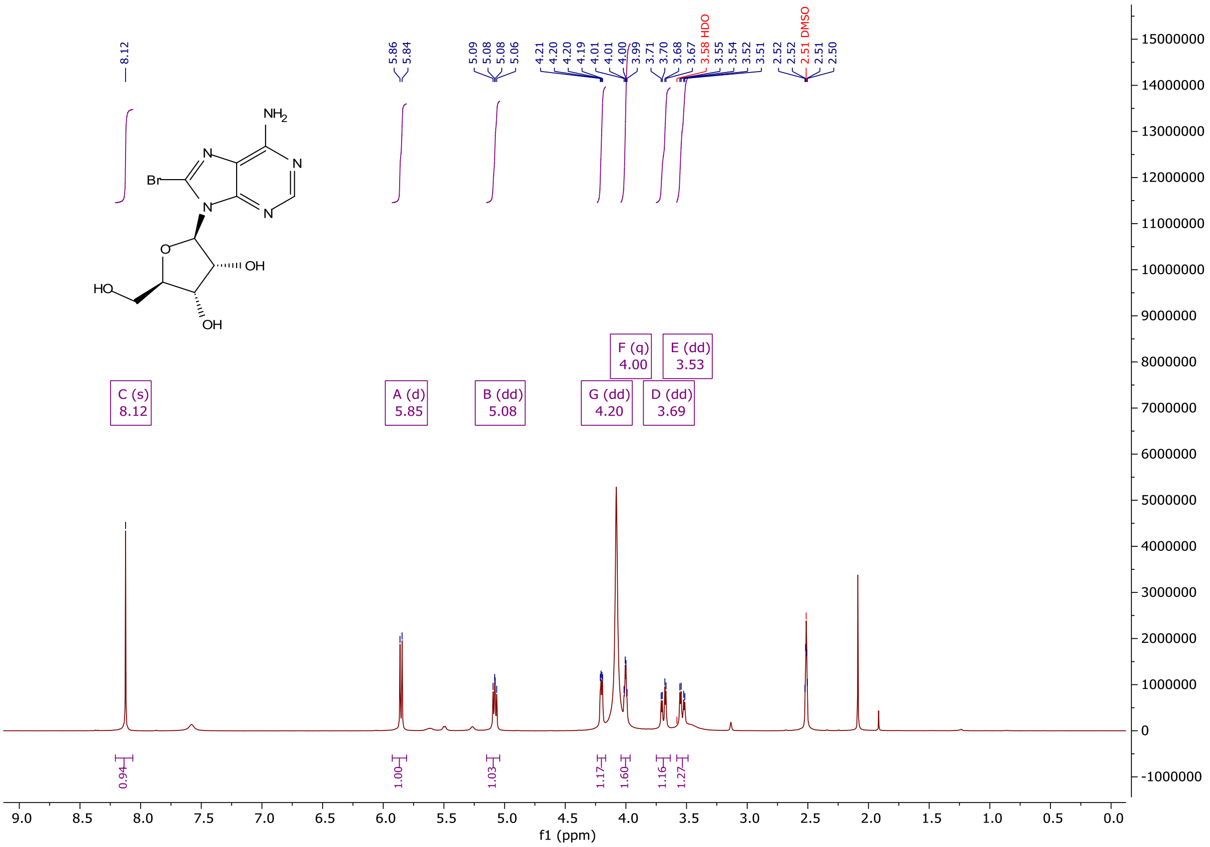

**$^{13}$C NMR of (2R,3R,4S,5R)-2-(6-amino-8-bromo-9H-purin-9-yl)-5-(hydroxymethyl)tetra hydrofuran-3,4-diol (8Br-Ad)**

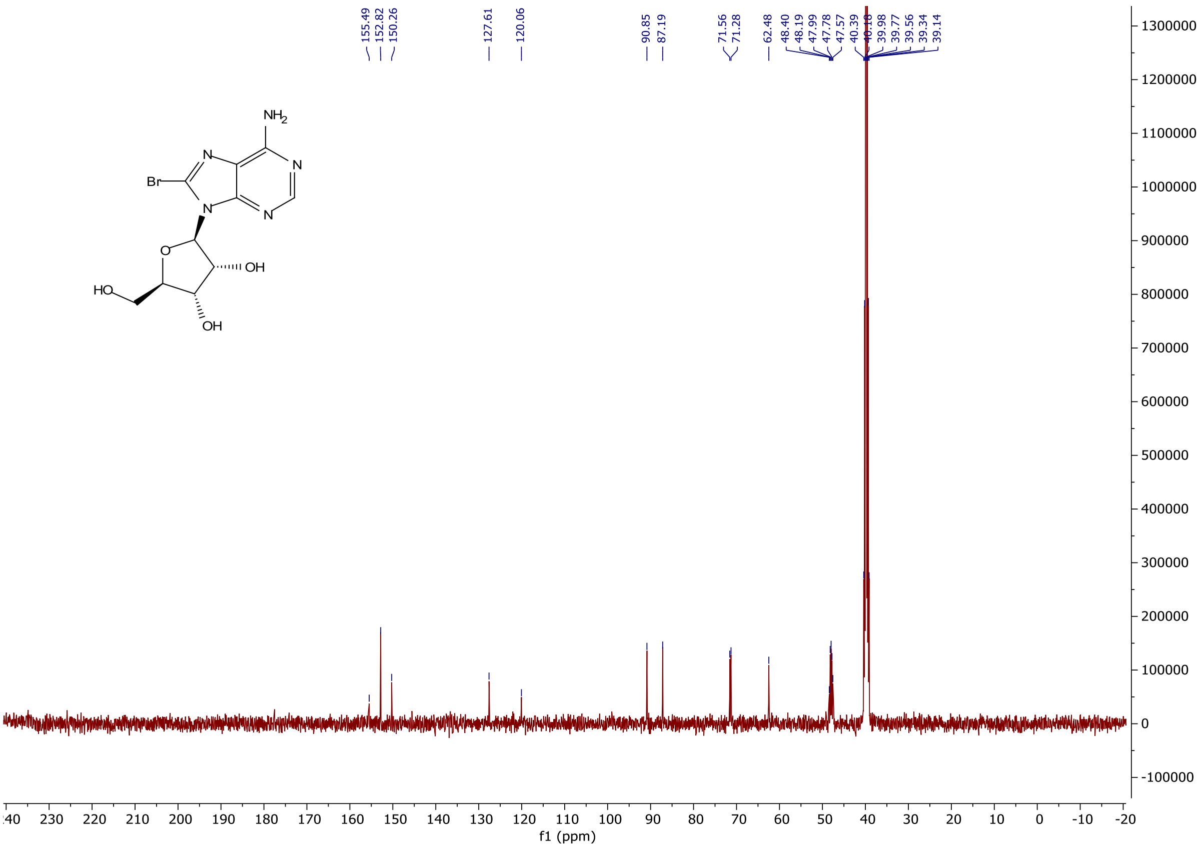

**[1]H NMR of (2R,3R,4S,5R)-2-(6-amino-8-(but-3-yn-1-ylthio)-9H-purin-9-yl)-5-(hydroxy methyl)tetrahydrofuran-3,4-diol (8But-Yn-Ad)**

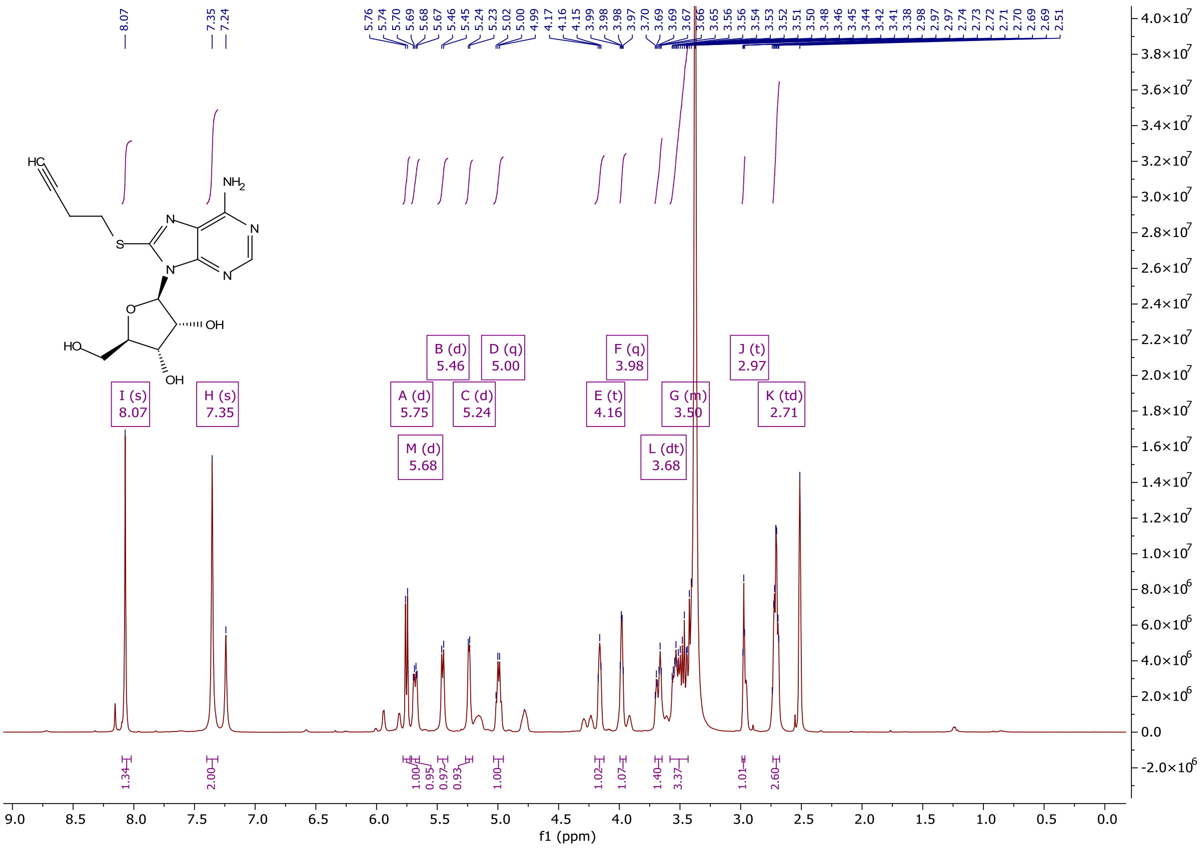

**[13]C NMR of (2R,3R,4S,5R)-2-(6-amino-8-(but-3-yn-1-ylthio)-9H-purin-9-yl)-5-(hydroxy methyl)tetrahydrofuran-3,4-diol (8But-Yn-Ad)**

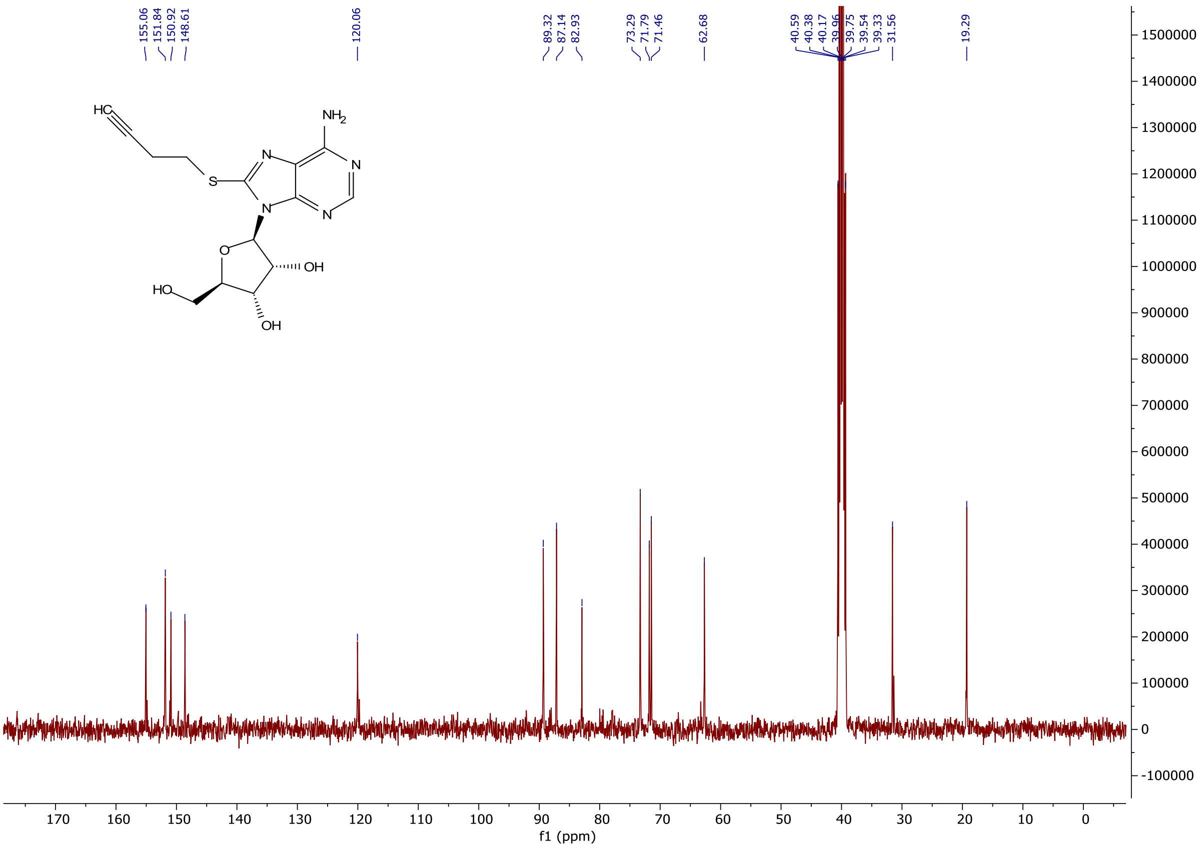

**[1]H NMR of 3-carbamoyl-1-((2R,3R,4R,5R)-3,4-diacetoxy-5-(acetoxymethyl)tetrahydrofuran-2-yl)pyridin-1-ium (NR-OAc)**

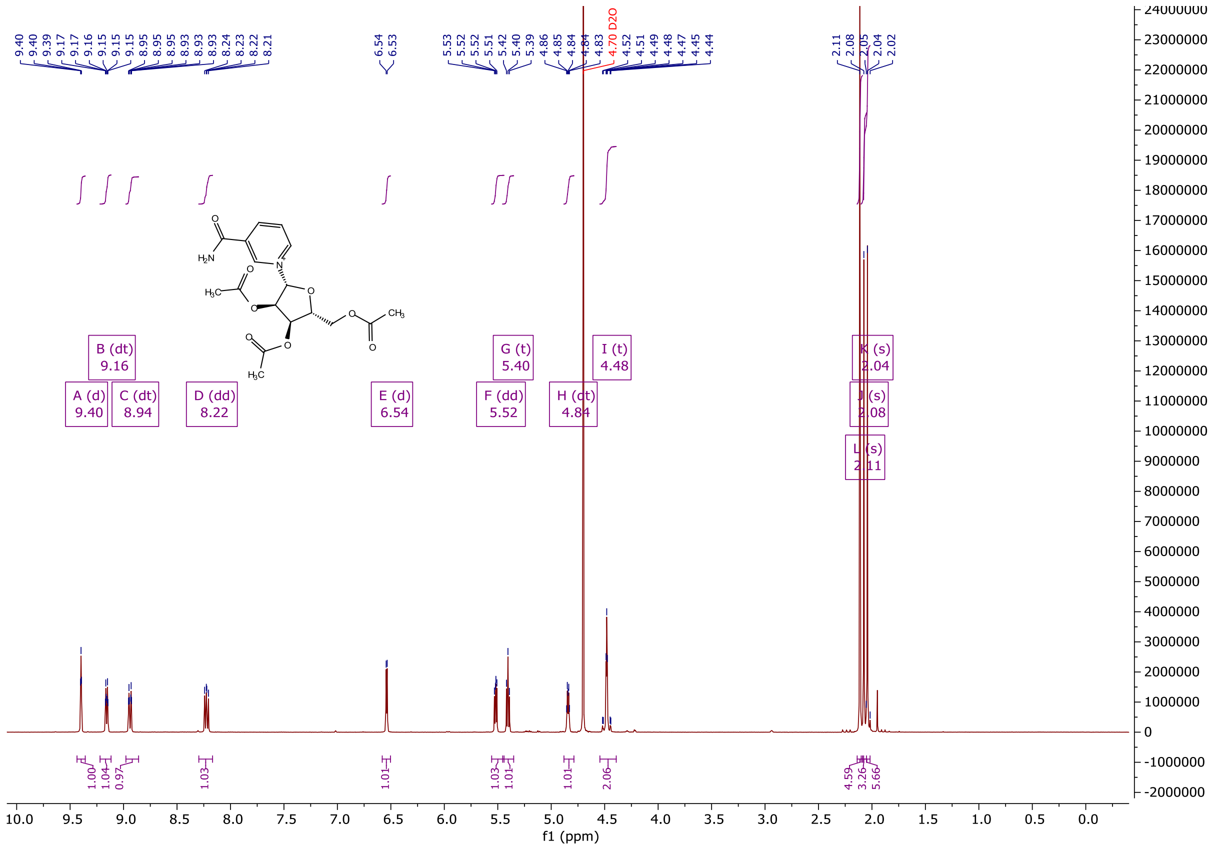

**$^{13}$C NMR of 3-carbamoyl-1-((2R,3R,4R,5R)-3,4-diacetoxy-5-(acetoxymethyl)tetrahydrofuran-2-yl)pyridin-1-ium (NR-OAc)**

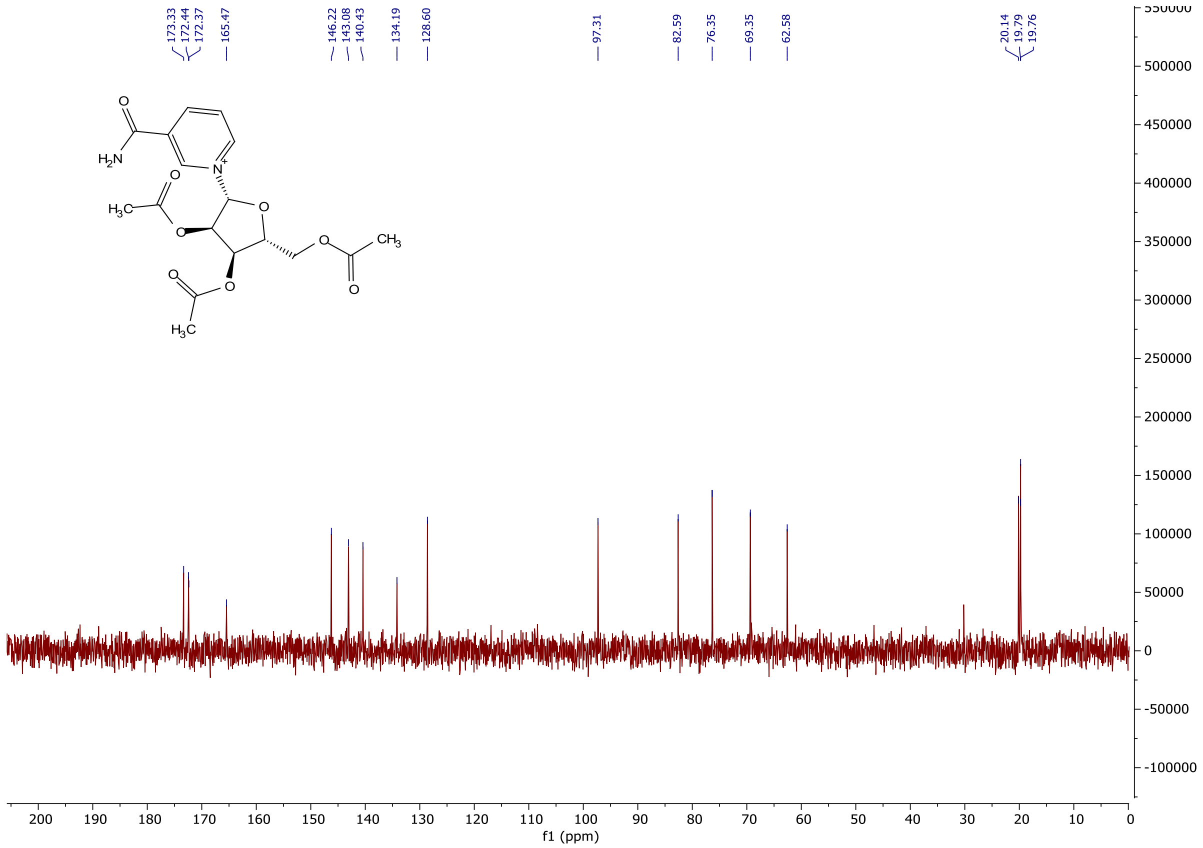

$^{1}H$ NMR of 1-((2R,3R,4S,5R)-3,4-dihydroxy-5-(hydroxymethyl)tetrahydrofuran-2-yl)-1,4-dihydropyridine-3-carboxamide (NRH)

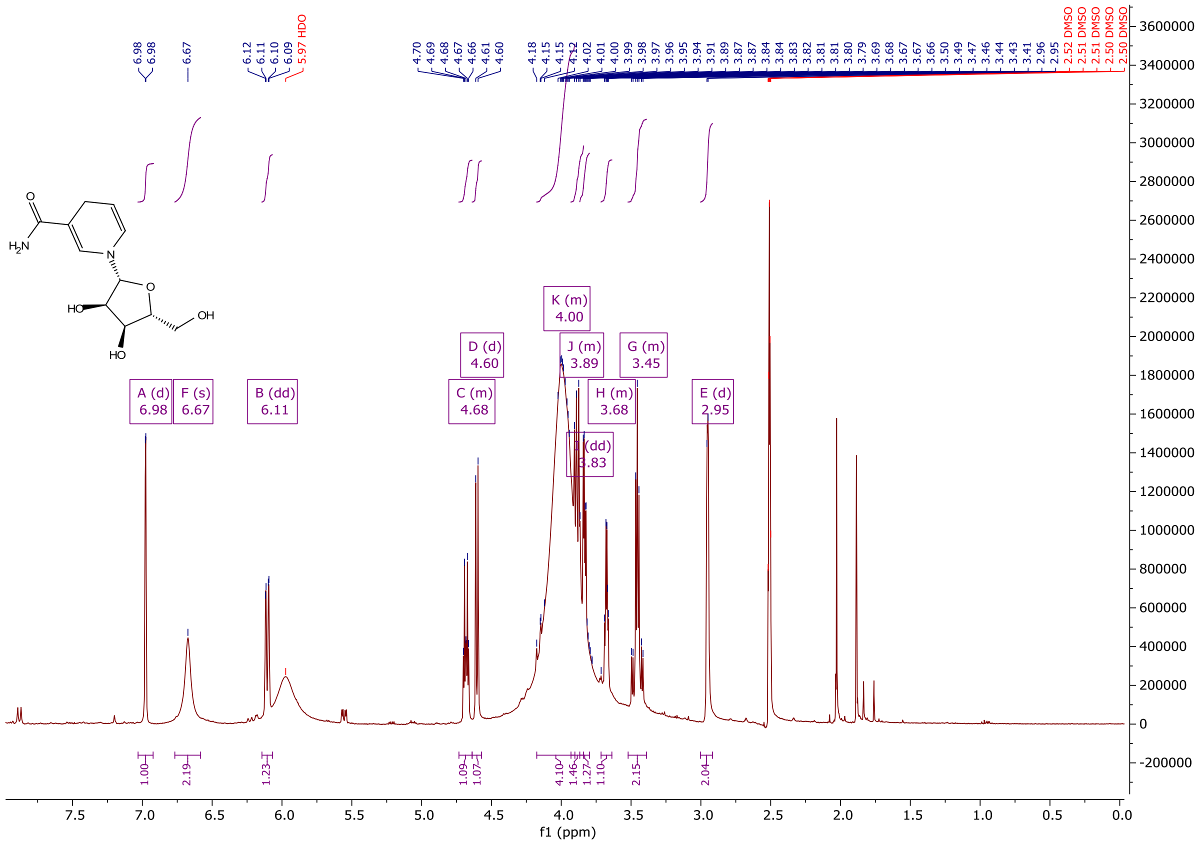

**$^{13}$C NMR of 1-((2R,3R,4S,5R)-3,4-dihydroxy-5-(hydroxymethyl)tetrahydrofuran-2-yl)-1,4-dihydro-pyridine-3-carboxamide (NRH)**

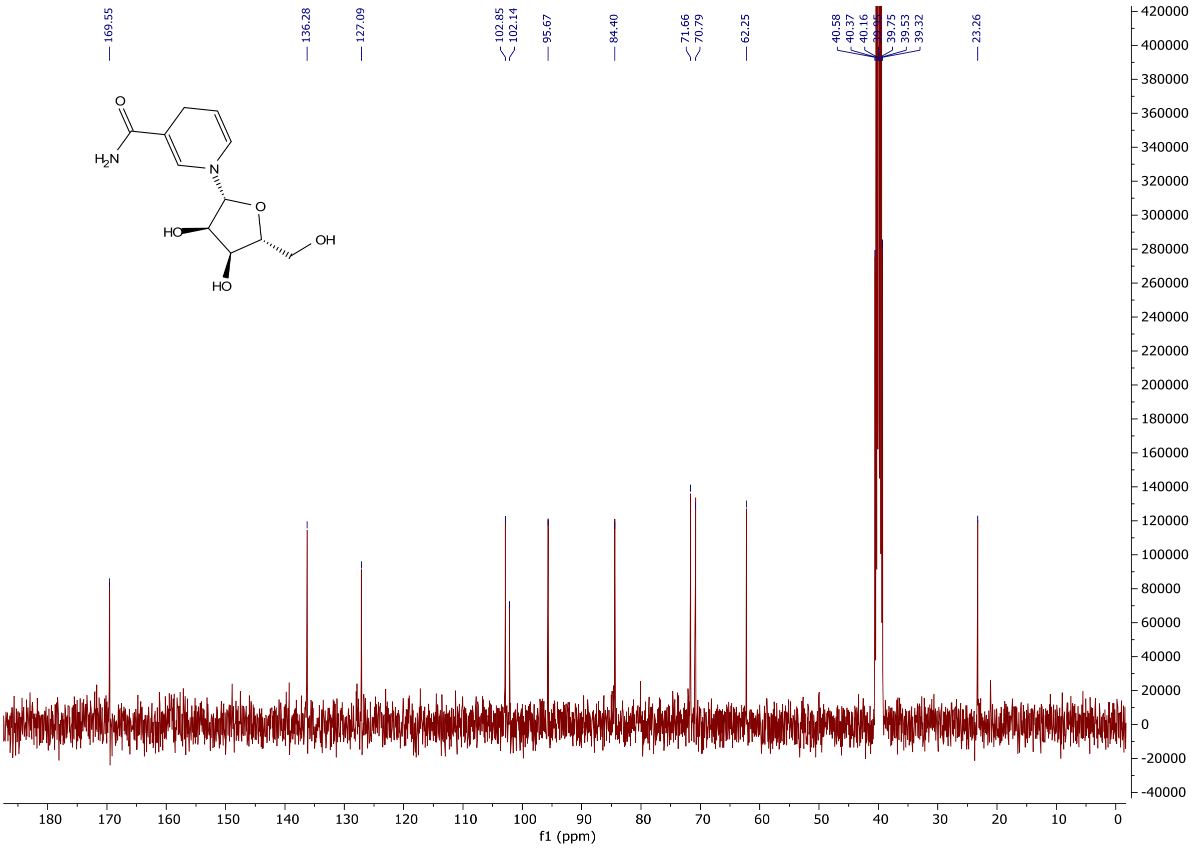

**$^{1}$H NMR of Benzyl ((((2R,3S,4R,5R)-3,4-dihydroxy-5-(6-(prop-2-yn-1-ylamino)-9H-purin-9-yl)tetrahydrofuran-2-yl)methoxy)(phenoxy)phosphoryl)-L-alaninate (6Yn-Pro)**

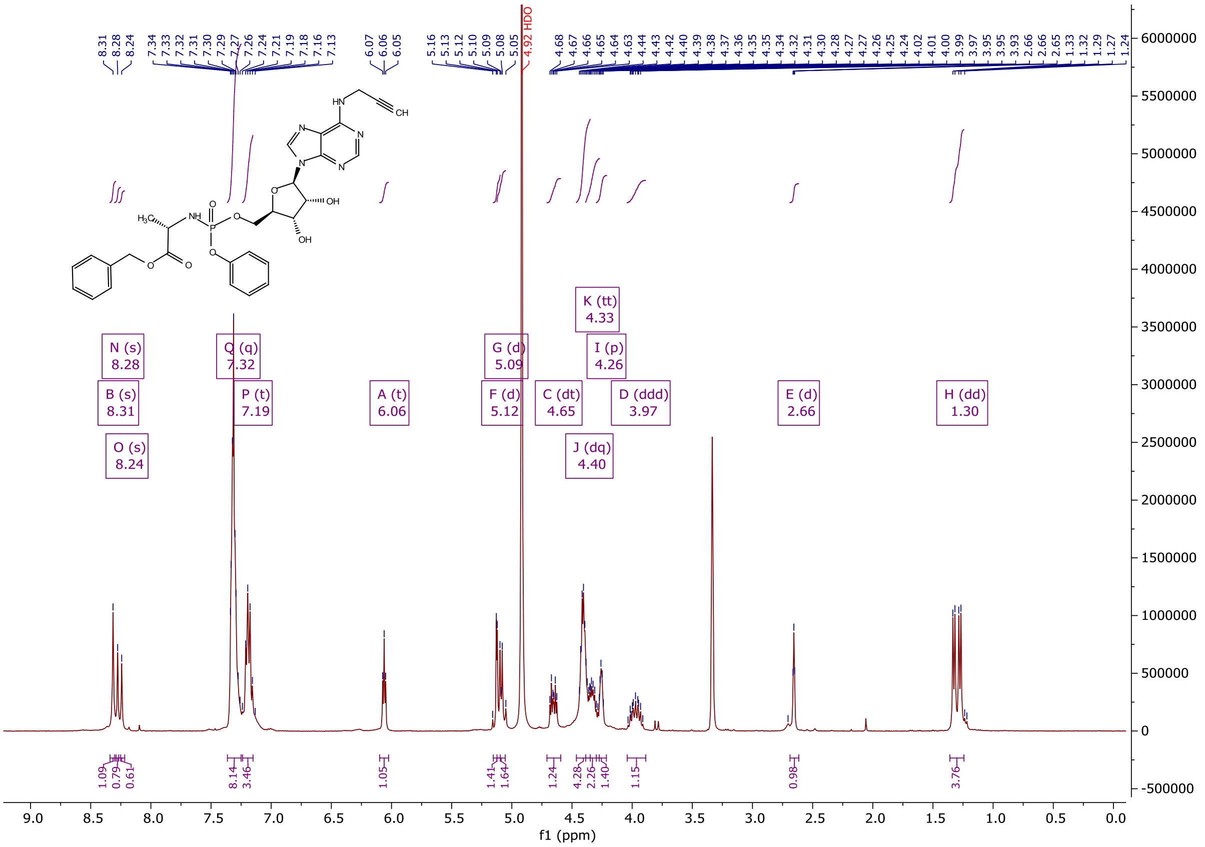

**$^{13}$C NMR of Benzyl ((((2R,3S,4R,5R)-3,4-dihydroxy-5-(6-(prop-2-yn-1-ylamino)-9H-purin-9-yl)tetrahydrofuran-2-yl)methoxy)(phenoxy)phosphoryl)-L-alaninate (6Yn-Pro)**

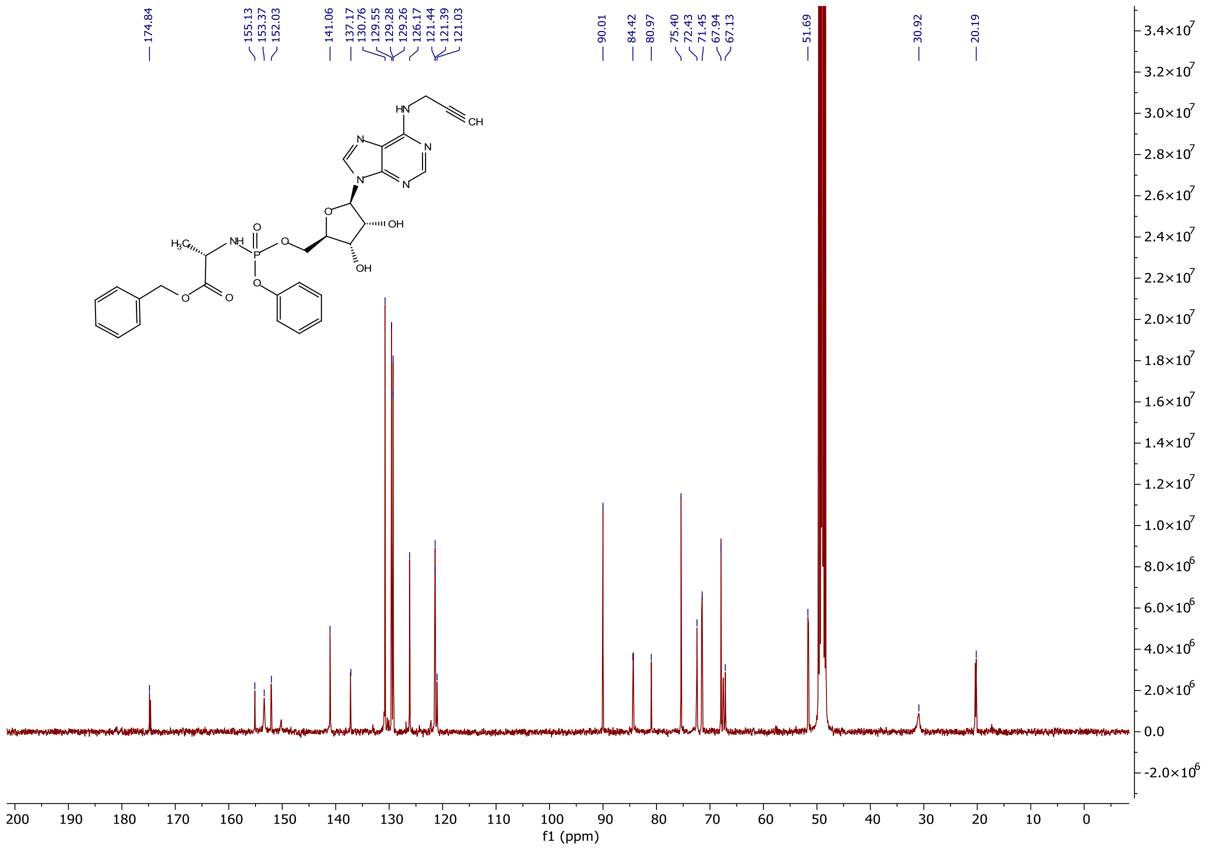

**$^{31}$P NMR of Benzyl ((((2R,3S,4R,5R)-3,4-dihydroxy-5-(6-(prop-2-yn-1-ylamino)-9H-purin-9-yl)tetrahydrofuran-2-yl)methoxy)(phenoxy)phosphoryl)-L-alaninate (6Yn-Pro)**

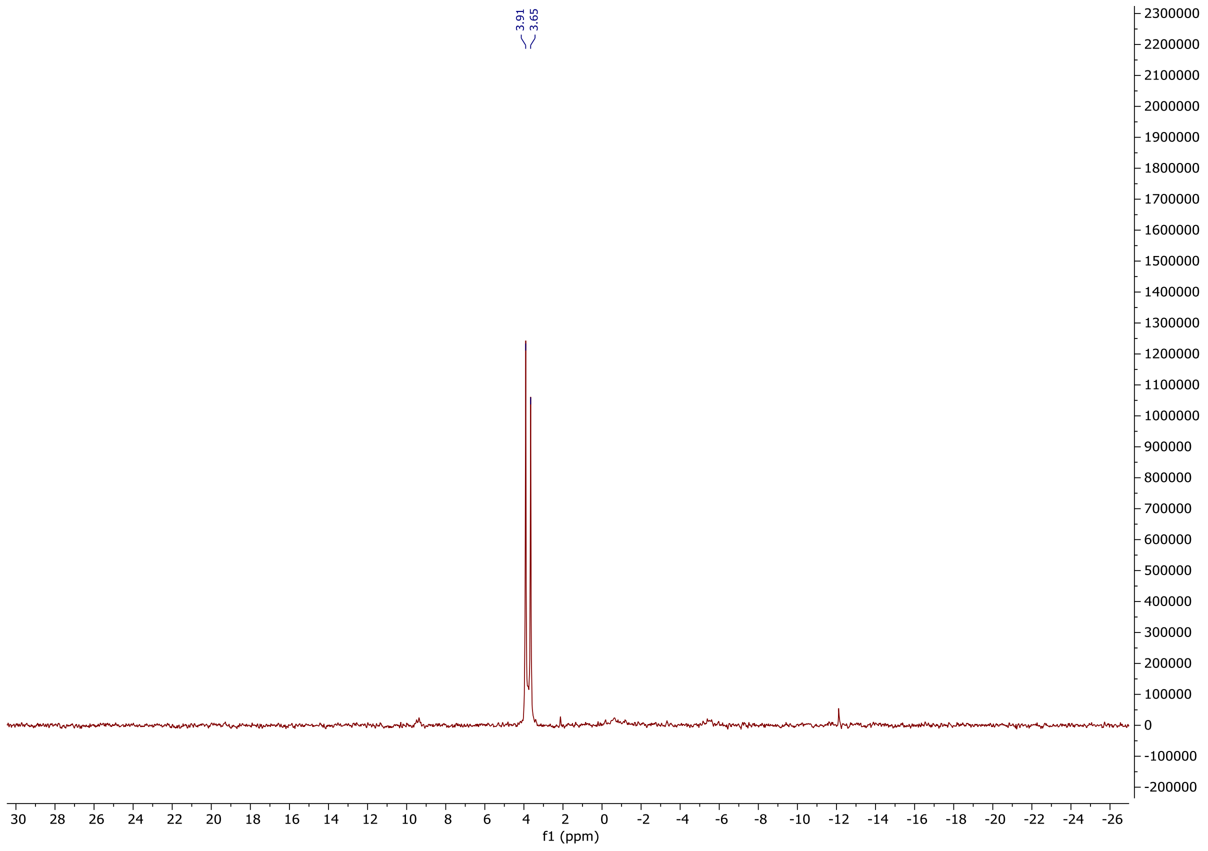

**$^{1}$H NMR of Benzyl ((((2R,3S,4R,5R)-5-(6-amino-2-ethynyl-9H-purin-9-yl)-3,4-dihydroxy tetrahydrofuran-2-yl)methoxy)(phenoxy)phosphoryl)-L-alaninate (2Yn-Pro)**

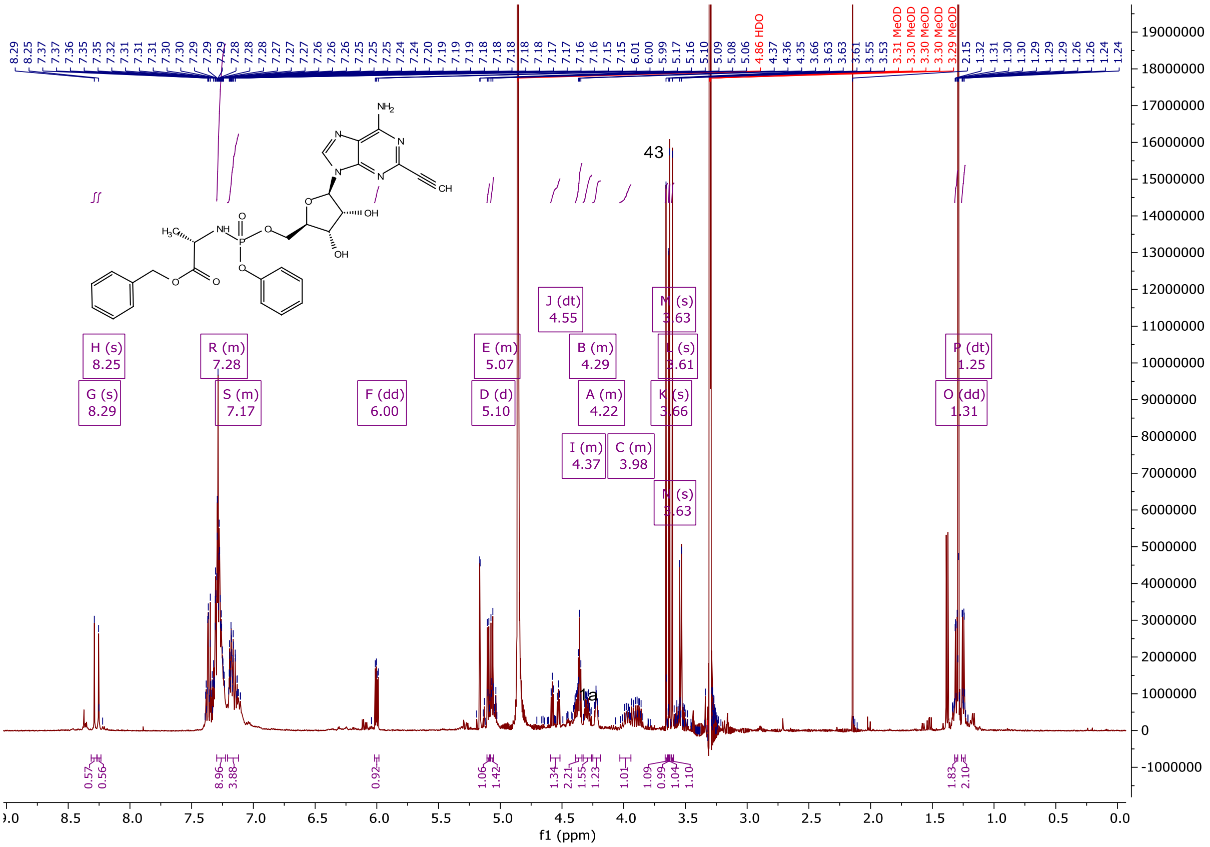

**$^{13}$C NMR of Benzyl ((((2R,3S,4R,5R)-5-(6-amino-2-ethynyl-9H-purin-9-yl)-3,4-dihydroxy tetrahydrofuran-2-yl)methoxy)(phenoxy)phosphoryl)-L-alaninate (2Yn-Pro)**

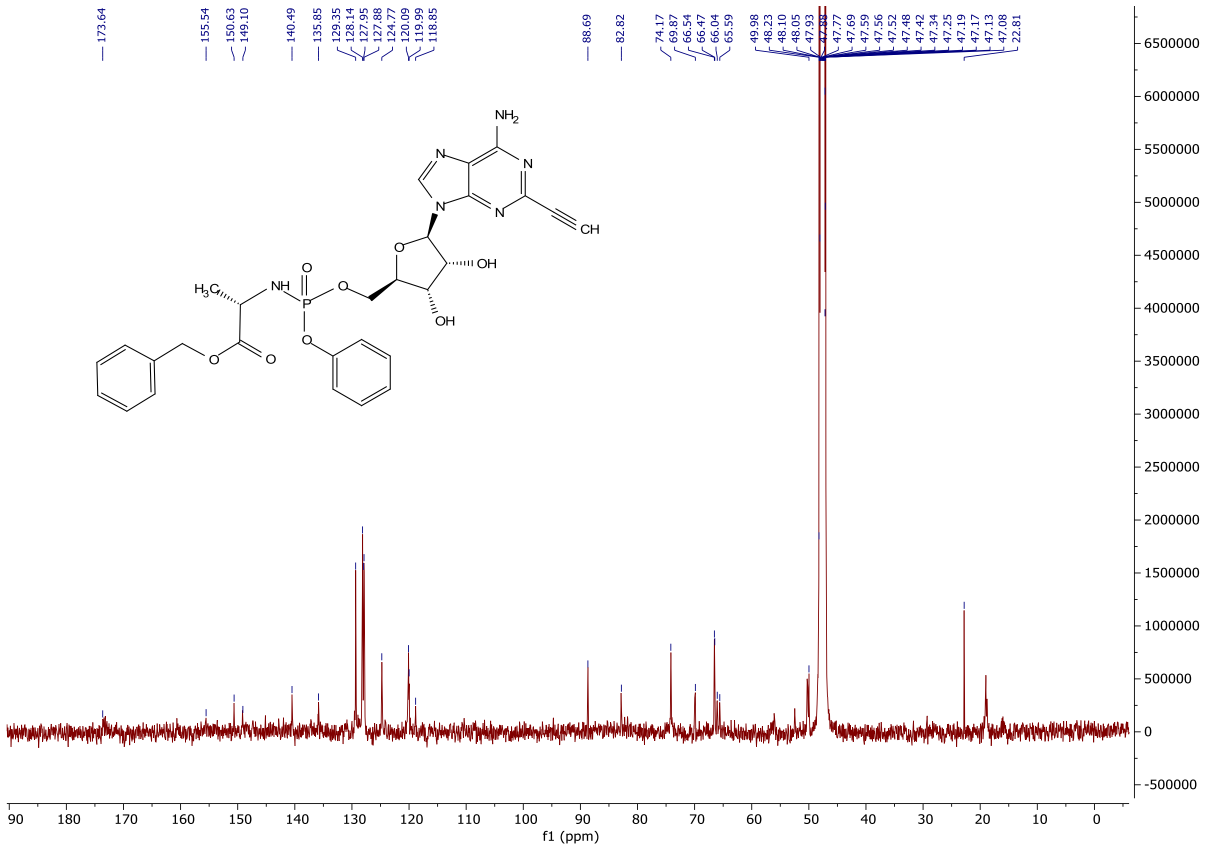

**$^{31}$P NMR of Benzyl ((((2R,3S,4R,5R)-5-(6-amino-2-ethynyl-9H-purin-9-yl)-3,4-dihydroxy tetrahydrofuran-2-yl)methoxy)(phenoxy)phosphoryl)-L-alaninate (2Yn-Pro)**

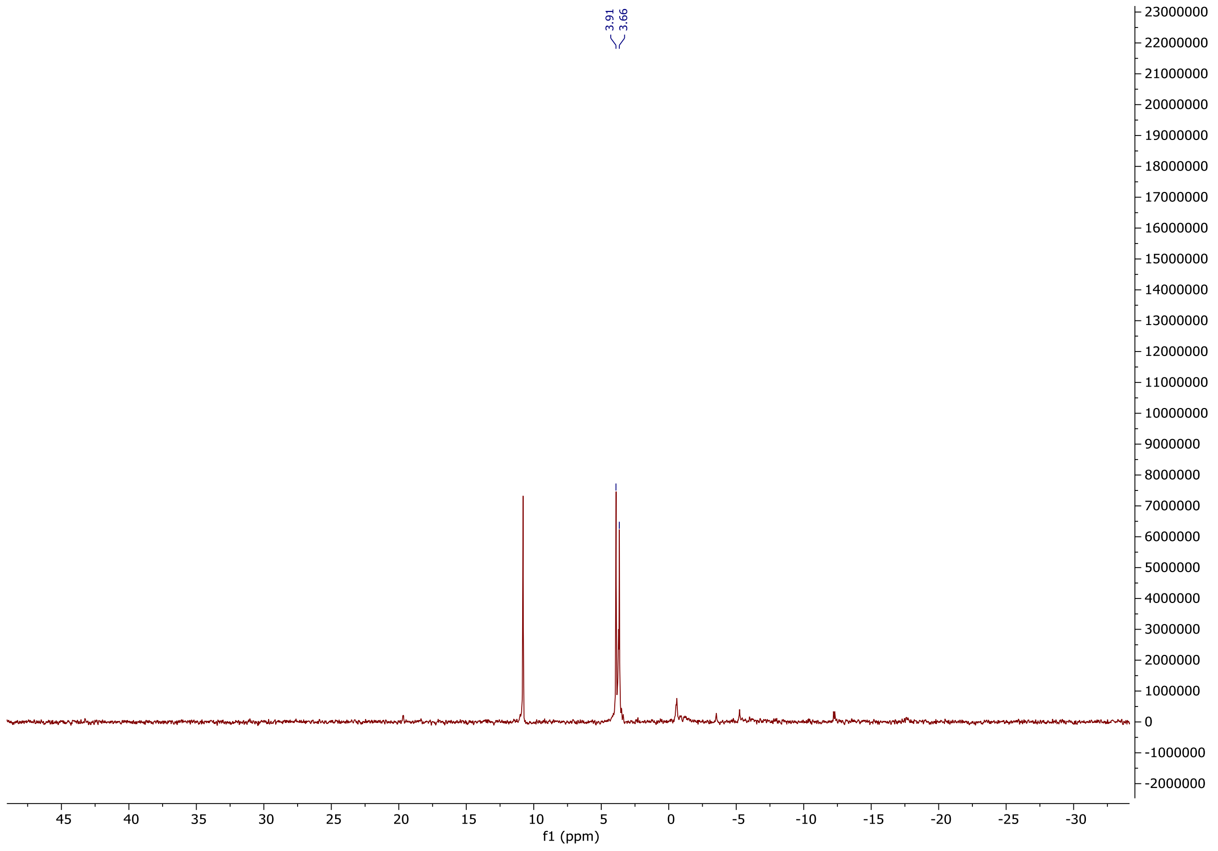

**[1]H NMR of Benzyl ((((2R,3S,4R,5R)-5-(6-amino-8-(but-3-yn-1-ylthio)-9H-purin-9-yl)-3,4-dihydroxytetrahydrofuran-2-yl)methoxy)(phenoxy)phosphoryl)-L-alaninate (8But-Yn-Pro)**

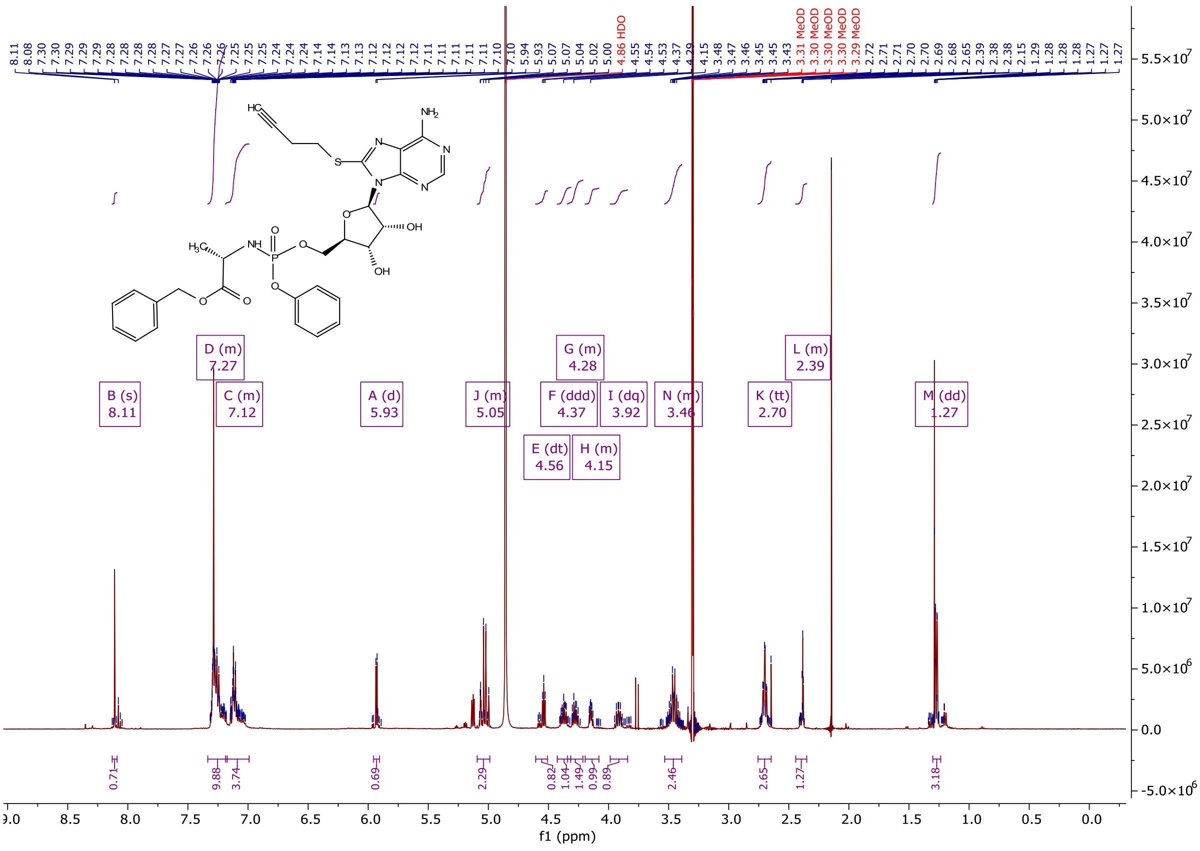

**$^{13}$C NMR of Benzyl ((((2R,3S,4R,5R)-5-(6-amino-8-(but-3-yn-1-ylthio)-9H-purin-9-yl)-3,4-dihydroxytetrahydrofuran-2-yl)methoxy)(phenoxy)phosphoryl)-L-alaninate (8But-Yn-Pro)**

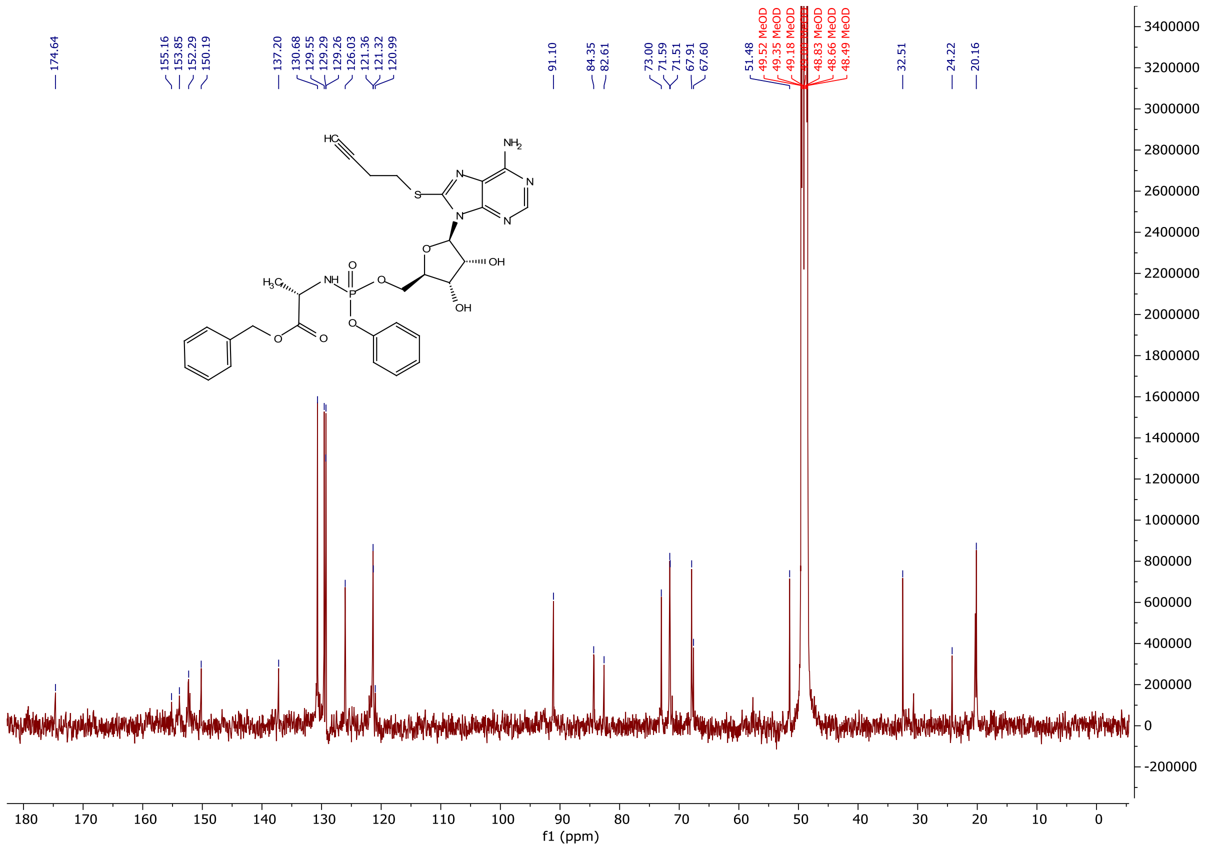

**$^{31}$P NMR of Benzyl ((((2R,3S,4R,5R)-5-(6-amino-8-(but-3-yn-1-ylthio)-9H-purin-9-yl)-3,4-dihydroxytetrahydrofuran-2-yl)methoxy)(phenoxy)phosphoryl)-L-alaninate (8But-Yn-Pro)**

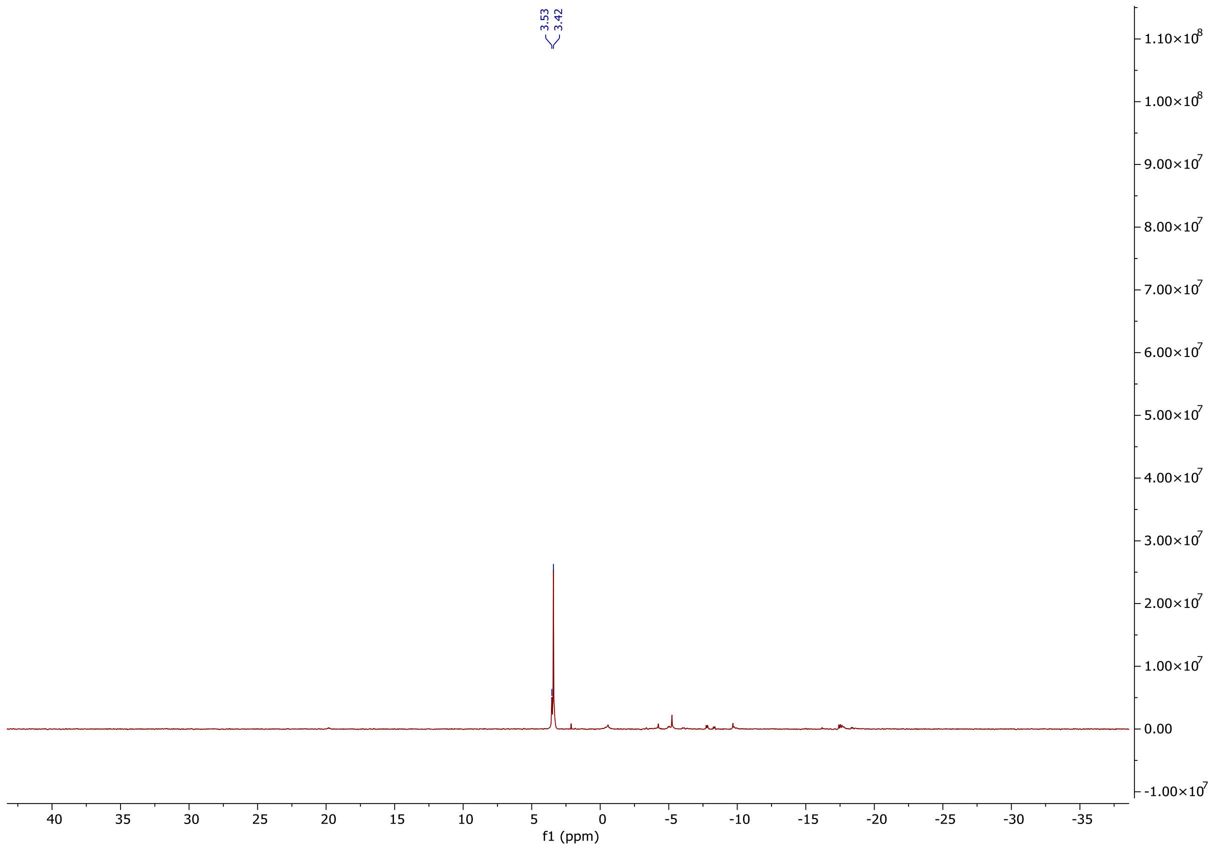

**[1]H NMR of ((2R,3S,4R,5R)-3,4-dihydroxy-5-(6-(prop-2-yn-1-ylamino)-9H-purin-9-yl)tetra hydrofuran-2-yl)methyl dihydrogen phosphate (6Yn-AMP)**

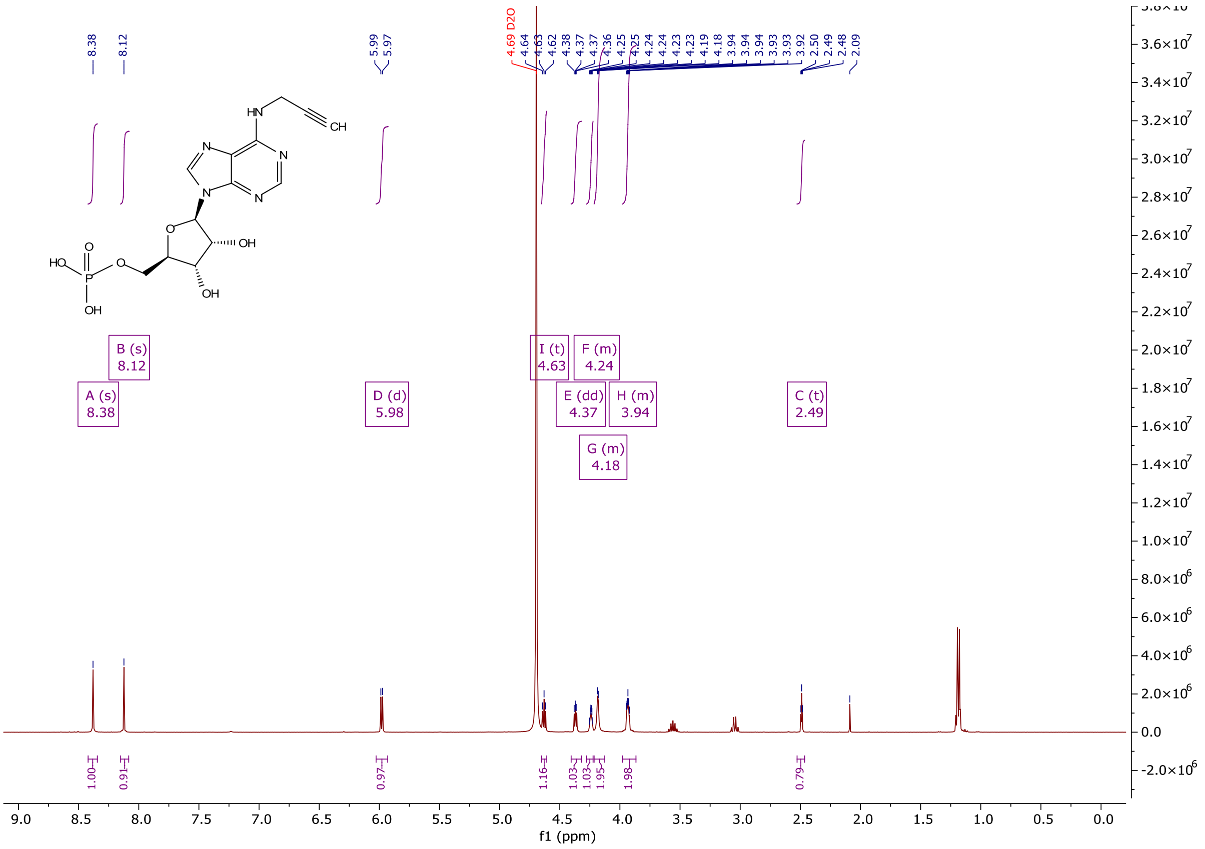

**$^{13}$C NMR of ((2R,3S,4R,5R)-3,4-dihydroxy-5-(6-(prop-2-yn-1-ylamino)-9H-purin-9-yl)tetra hydro-furan-2-yl)methyl dihydrogen phosphate (6Yn-AMP)**

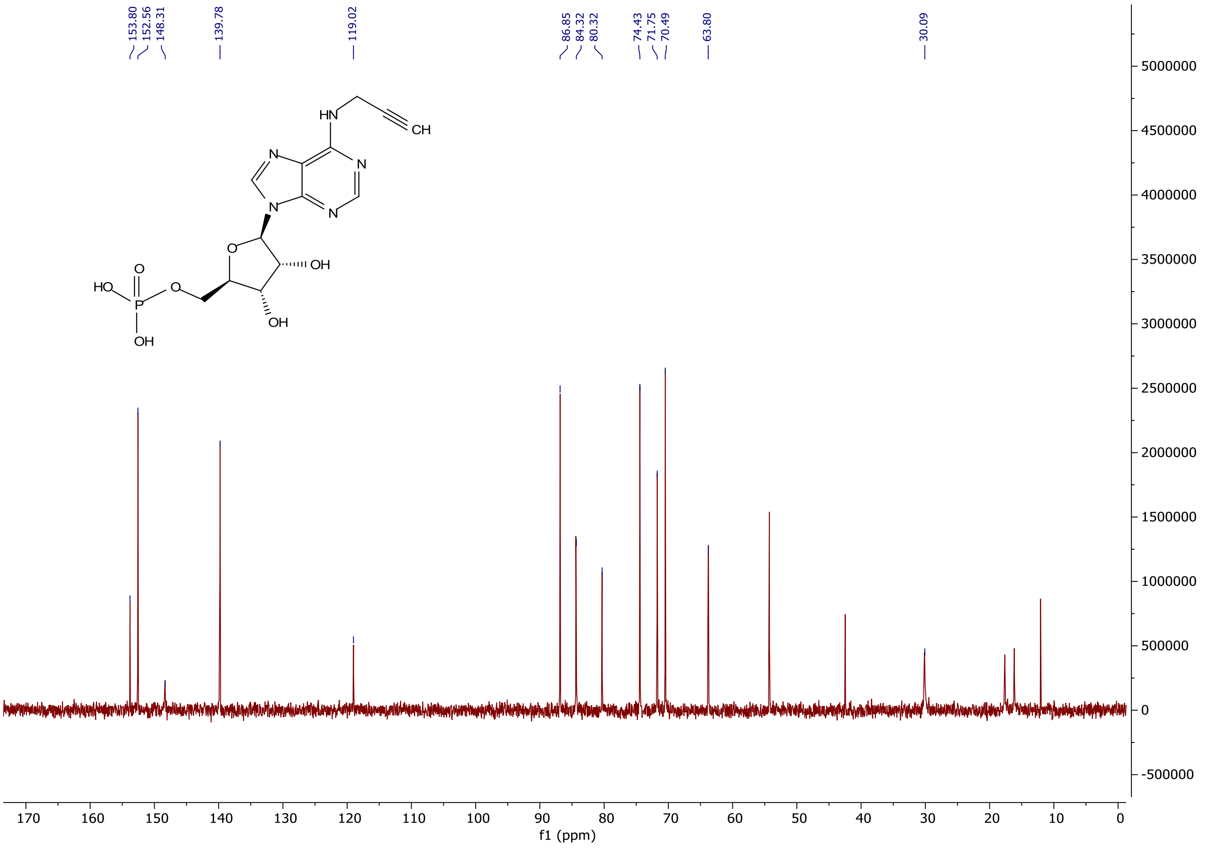

**$^{31}$P NMR of ((2R,3S,4R,5R)-3,4-dihydroxy-5-(6-(prop-2-yn-1-ylamino)-9H-purin-9-yl)tetra hydro-furan-2-yl)methyl dihydrogen phosphate (6Yn-AMP)**

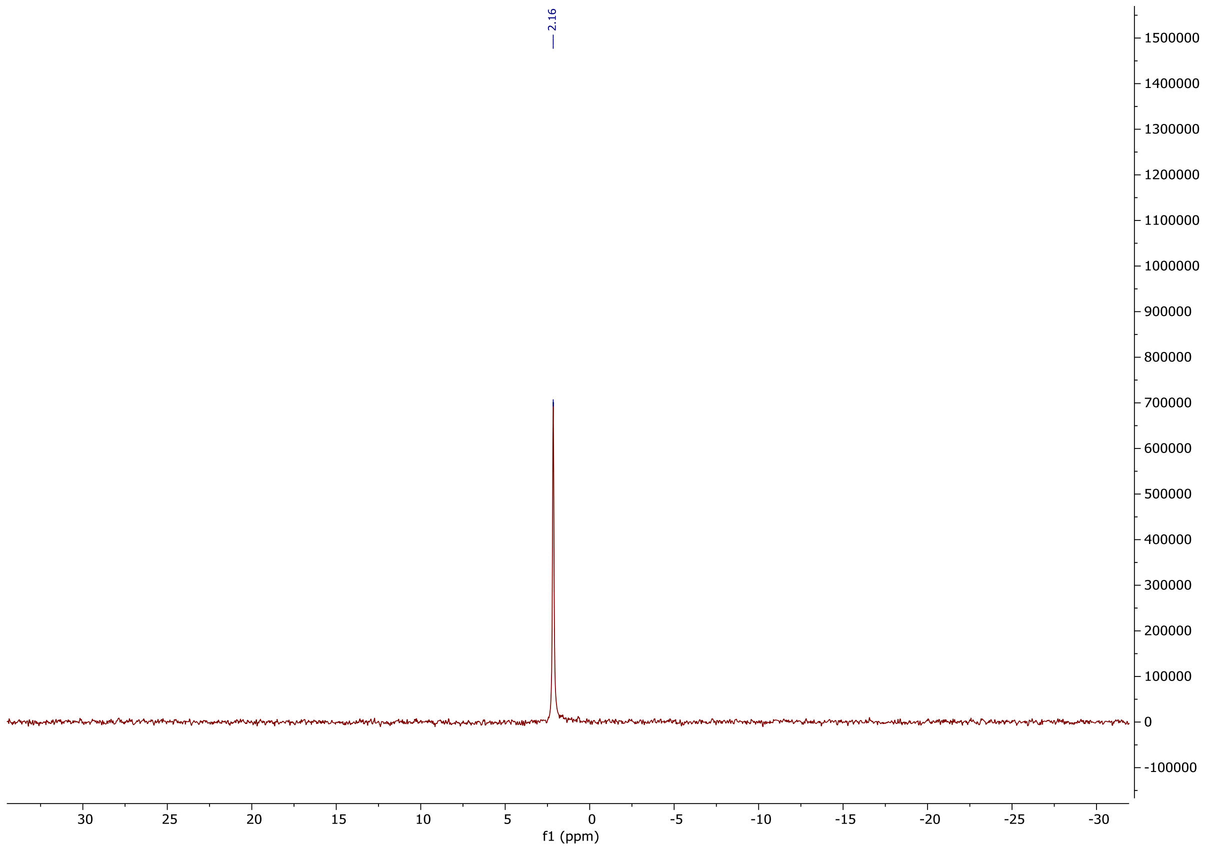

**[1]H NMR of ((2R,3S,4R,5R)-3,4-dihydroxy-5-(6-(prop-2-yn-1-ylamino)-9H-purin-9-yl)tetra hydrofuran-2-yl)methyl tetrahydrogen triphosphate (6Yn-ATP)**

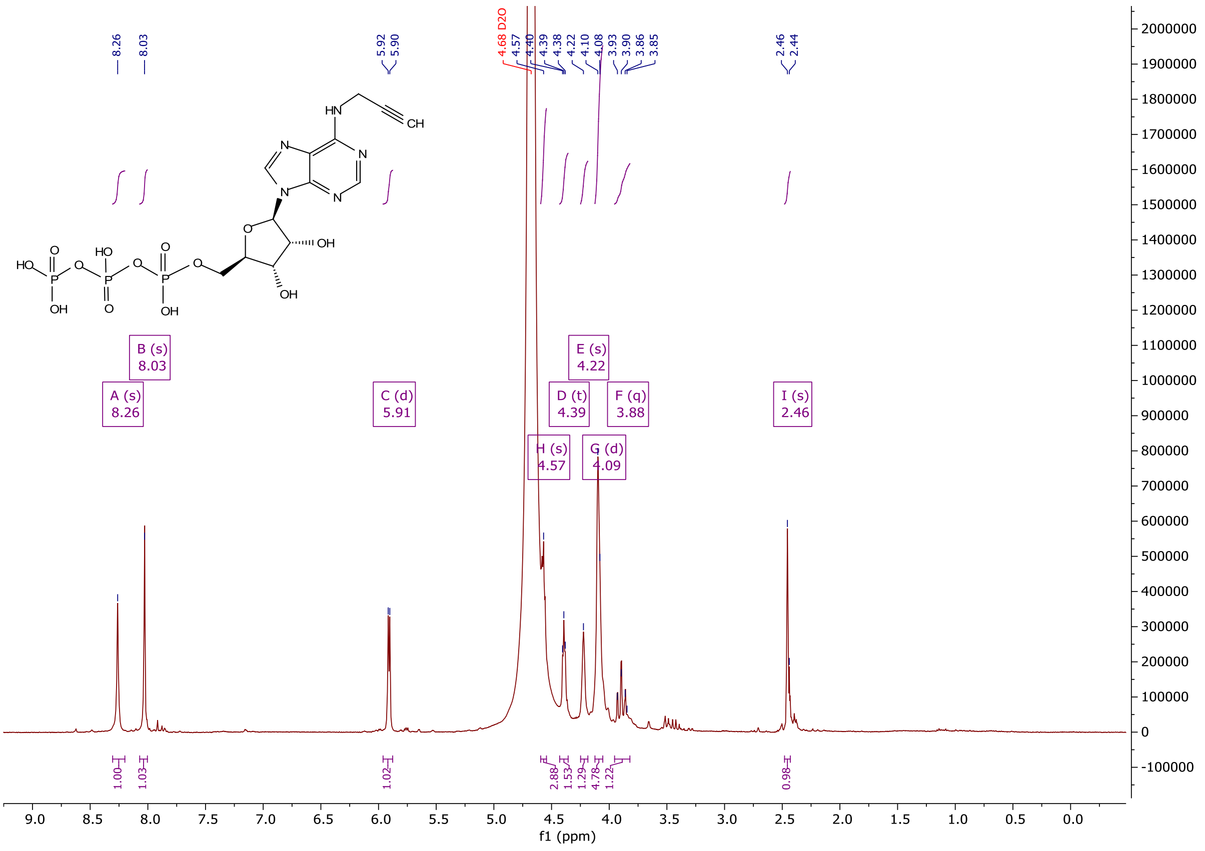

**$^{13}$C NMR of ((2R,3S,4R,5R)-3,4-dihydroxy-5-(6-(prop-2-yn-1-ylamino)-9H-purin-9-yl)tetra hydro-furan-2-yl)methyl tetrahydrogen triphosphate (6Yn-ATP)**

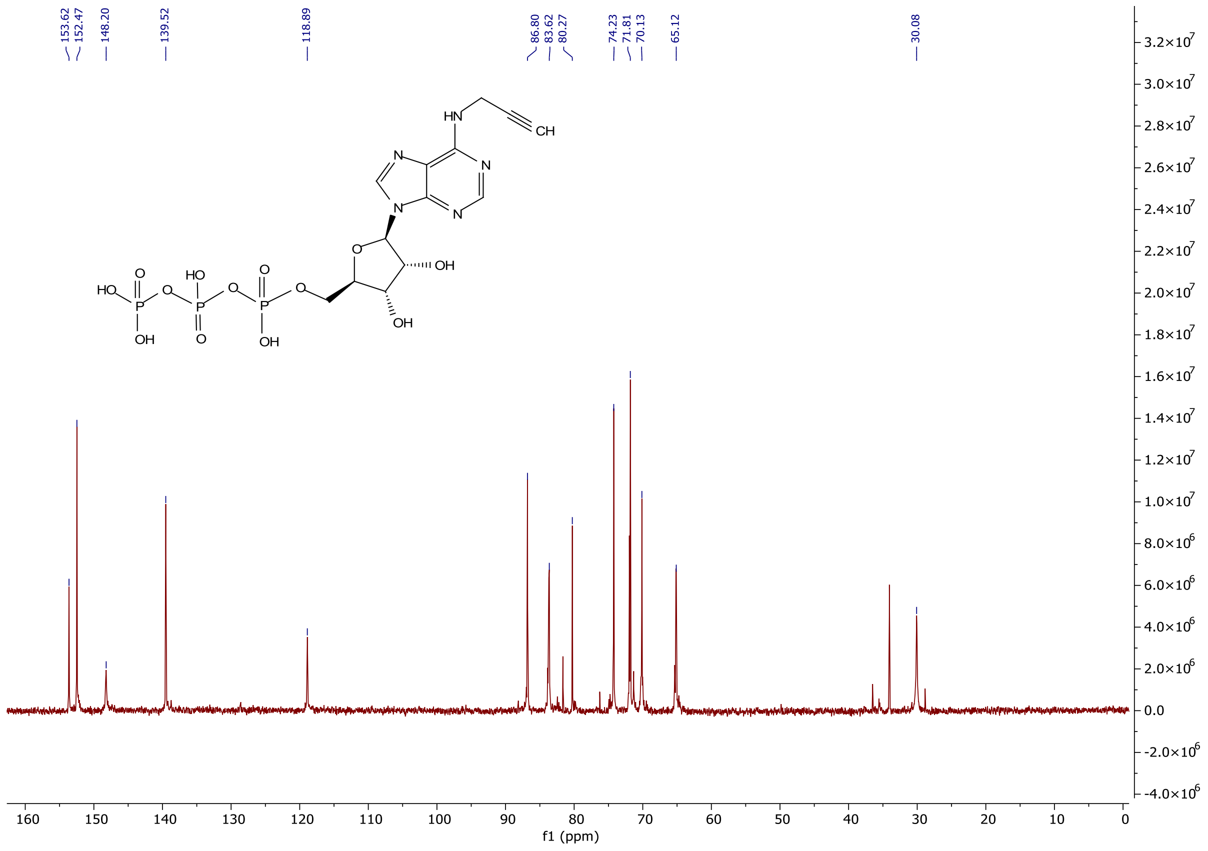

**[31]P NMR of ((2R,3S,4R,5R)-3,4-dihydroxy-5-(6-(prop-2-yn-1-ylamino)-9H-purin-9-yl)tetra hydro-furan-2-yl)methyl tetrahydrogen triphosphate (6Yn-ATP)**

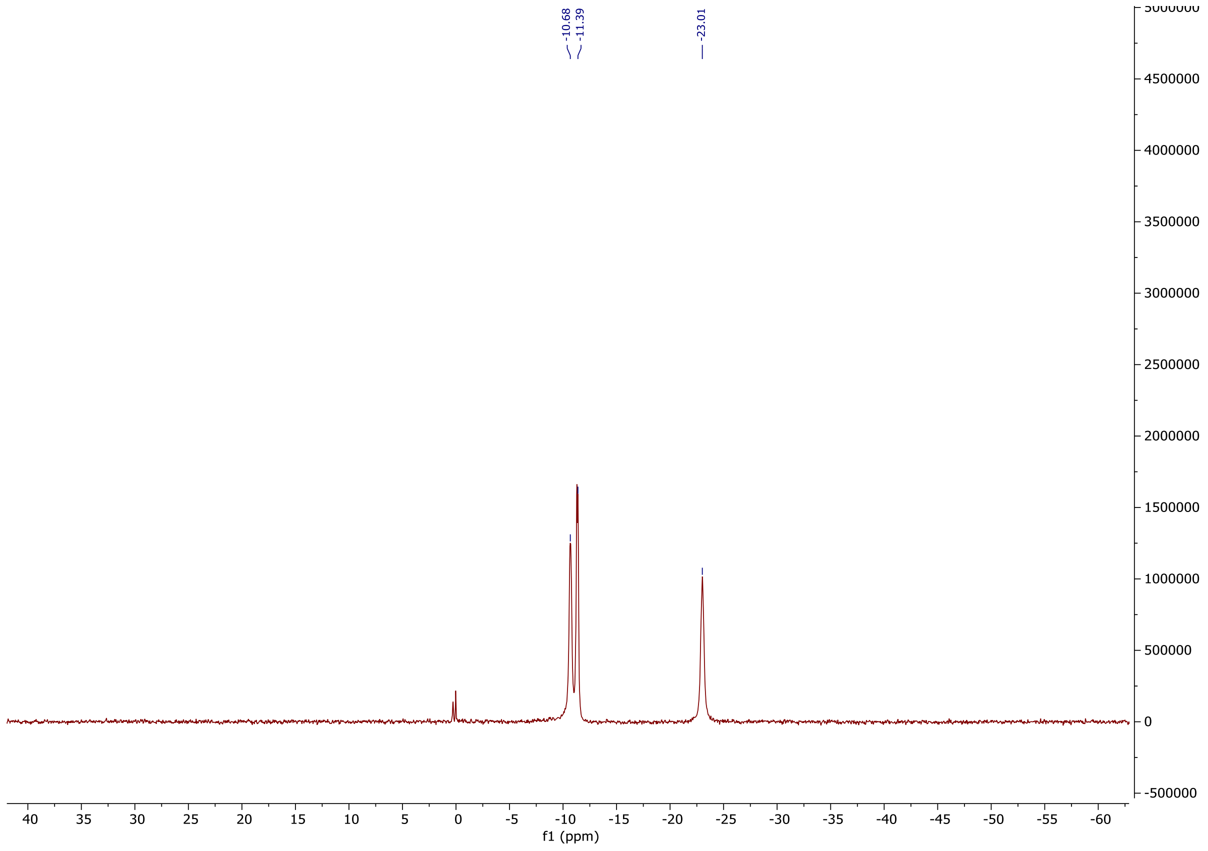
